# Connectional neuroanatomy of U-fibers in the rhesus monkey brain

**DOI:** 10.64898/2026.06.06.730597

**Authors:** Tyler J. Capen, Lauren J. O’Donnell, Fan Zhang, Edward H. Yeterian, Yogesh Rathi, Nikos Makris, Douglas L. Rosene, R. Jarrett Rushmore

**Author notes:** Corresponding author Dr. R.J. Rushmore, Ph.D., 650 Albany Street, X327, Department of Anatomy and Neurobiology Boston, University School of Medicine Boston, MA 02118, USA.

## Abstract

The superficial white matter (SWM), the region of white matter immediately beneath the gray matter-white matter (GM-WM) border, contains short cortico-cortical association fibers that interconnect neighboring cortical regions. The SWM is estimated to comprise the majority of axons in the cerebral white matter and is thus thought to play a major role in cortical information processing. Despite this prominence, and growing attention in the diffusion MRI (dMRI) field, the connectional organization of the SWM remains poorly defined. A major component of the SWM is U-shaped fiber bundles, classically depicted in brain dissections as bundles beneath sulci that interconnect adjacent gyri. The prevailing view, from both brain dissection and dMRI, considers U-fibers to be ubiquitous elements of the SWM that form symmetric connections between corresponding portions of adjacent gyri beneath all sulci, and they are routinely reconstructed in tractography as a layer of short association connections. However, their actual prevalence and organization in the primate brain have never been systematically evaluated with experimental tract tracing. To address this gap, we analyzed 28 archival macaque tract-tracing cases in which anterograde tracers were injected on gyri adjacent to major cortical sulci (intraparietal, central, principal, superior temporal), all of which have clear anatomical correspondences to sulci in the human cerebral cortex. Several organizational principles emerged. First, although short association fibers were present in every case, U-fibers - operationally defined as labeled fibers that left the injection site, passed beneath a sulcus, and terminated on the adjacent gyrus - were only observed in a minority of cases. Second, U-fiber incidence was sulcus-dependent: U-fibers were consistently present beneath the principal and central sulci, but uncommon beneath the intraparietal and superior temporal sulci. Third, when present, U-fibers did not follow a uniform trajectory - some took a course symmetric and orthogonal to the sulcus, whereas others followed oblique trajectories to more distant regions. In addition to U-fibers, we identified shorter association fiber bundles that terminated on the sulcal bank proximal to the injection, or on the opposing bank after they crossed the fundus of the sulcus, evidence that the SWM contains multiple classes of short association fiber bundles rather than a single canonical U-fiber system. We further identified a consistent 200-300 µm band of white matter beneath cortical layer 6 that contained exclusively short association fibers on approach to their cortical terminations and was selectively avoided by deeper long cortico-cortical pathways. This band was present across all examined sulci, suggesting that it is a conserved feature of the cortical white matter. Notably, U-fibers were found both within this band and in the white matter below. We also identified deeper fiber bundles that adopted a U-shaped course beneath a sulcus but did not interconnect adjacent gyri. Thus, a U-shaped trajectory alone does not define a U-fiber. Together, these findings constitute the first systematic connectional study of cortical U-fibers in major sulci of the non-human primate brain and indicate that the white matter beneath the cortex is a highly complex system in which short association fiber bundle organization reflects the selective connectivity of the cortical areas separated by a sulcus, rather than a uniform U-fiber architecture imposed by the geometry of the overlying sulcus. These results have potential consequences for dMRI and anatomical studies of white matter and provide an anatomically grounded basis to better define SWM organization in anatomy and tractography.

## 1. Introduction

The cerebral cortex consists of interconnected regions organized into structural and functional networks (Corbetta, 2012; Bullmore & Sporns, 2012; Ye et al., 2015; Glasser et al., 2016; de Pasquale et al., 2018; Siegel et al., 2022; Assem et al., 2024). These regions have been delineated by cytoarchitecture (Brodmann, 1909), laminar structure (Dombrowski et al., 2001; Hilgetag & Barbas, 2006), myelination (Vogt, 1910, 1911), neurotransmitter receptor distribution (Zilles & Palomero-Gallagher, 2017), and other features. In contrast, the organization of the underlying white matter, the large and complex compartment composed of axons that carry inputs to and outputs from cortical areas (Schmahmann & Pandya, 2006), has received comparatively less attention (Rockland & Rushmore, 2025). This is especially true for the superficial white matter (SWM), the zone immediately beneath the gray matter-white matter (GM-WM) border.

The SWM contains a large number of axons that originate from and terminate in nearby cortical regions. By some estimates, these short association fibers roughly outnumber those in the deep white matter by an order of magnitude (Schüz & Braitenberg, 2002). Classic dissection studies of the white matter beneath the cortex primarily described short cortico-cortical fiber bundles that travel beneath sulci and interconnect adjacent gyri (e.g., Luys, 1865; Dejerine & Dejerine-Klumpke, 1895). Because of their shape, these dissected bundles were referred to as arcuate fibers (Arnold, 1883) or U-fibers (Meynert, 1885). These descriptions led to the conceptualization of the SWM as an extensive layer of short association fibers oriented parallel to the GM-WM interface (Budde & Annese, 2013) and located beneath the entire cerebral cortex. Despite their abundance and the associated functional importance, the detailed connectional organization of these fibers, including their regional distribution, geometrical arrangement, and potential laminar position in the white matter, remains largely understudied and undefined at the level of their anatomical connections.

The SWM is also a region through which long-range axons projecting to or from the overlying cortex must pass through. Axons projecting from the overlying cortex include long-range cortico-cortical association pathways (e.g., the superior longitudinal fasciculus; Dejerine & Dejerine-Klumpke, 1895; Petrides & Pandya, 2002; Schmahmann & Pandya, 2006; Schmahmann et al., 2007) as well as projection pathways to telencephalic, diencephalic, brainstem, or spinal cord targets (e.g., Yeterian & Pandya, 1985; Stefanacci & Amaral, 2002; Frankle et al., 2006; Morecraft et al., 2019). Axons from brainstem, diencephalic, or telencephalic nuclei must also traverse the SWM to terminate in overlying cortical regions (e.g., Mufson et al., 1981; Mesulam & Mufson, 1984; Amaral & Price, 1984; Dermon & Barbas, 1994; Haber et al., 2000; Zikopoulos & Barbas, 2007). These axons traversing through the SWM are expected to be oriented perpendicular to the GM-WM interface and to the short association fibers in the SWM (Reveley et al., 2015).

The juxtaposition of these crossing orientations within the SWM has presented significant challenges for accurate reconstruction of fiber bundles with diffusion MRI (dMRI) (Reveley et al., 2015; Schilling et al., 2018). In regions where component fibers are not uniformly oriented, such as the SWM, tractography cannot reliably infer the trajectory of streamlines. Because the SWM has been estimated to comprise the majority of axons in the human brain (Braitenberg & Schüz, 1991; Schüz & Braitenberg, 2002), this limitation represents a major obstacle to the study of human brain connectivity and is therefore essential to consider when interpreting diffusion-based tractography.

The original anatomical depictions of U-fibers in classic literature (e.g., Arnold, 1883; Meynert, 1885; Ludwig & Klingler, 1956) have strongly influenced subsequent studies of SWM organization and led to several main principles of SWM organization. First, U-fibers are commonly depicted as ubiquitous features of the SWM and are present beneath most, if not all, sulci (e.g., Dejerine & Dejerine-Klumpke, 1895; Oishi et al., 2011; Guevara et al., 2020; Tian et al., 2025). Second, U-fibers are depicted as largely symmetrical in shape; they connect one gyrus to the corresponding portion of an adjacent gyrus by traveling in the SWM underneath a sulcus. Meynert described the arrangement of U-fibers along sulci, stating that they “resemble the half of a gun-barrel constructed of wire rings” (Meynert, 1885). Dejerine and Dejerine-Klumpke further stated that the direction of the U-fibers is perpendicular to the sulcus (“la direction des fibres en U est toujours perpendiculaire au grand axe du sillon qu’elles tapissent”; Dejerine & Dejerine-Klumpke, 1895, p. 748). Along the length of a sulcus, U-fibers are conceived to be arranged or stacked side by side, forming a smooth sheet of parallel arc-shaped fiber bundles linking corresponding portions of adjacent gyri. Finally, Meynert described the depth of U-fibers is directly related to their length such that the shortest fibers connect nearby regions and are located closest to the GM-WM boundary, while longer fibers that connect more distant regions are deeper in the white matter (Meynert, 1885). This arrangement was confirmed by Dejerine and Dejerine-Klumpke (1895) and supports a gradual progression in which U-fibers increase in length with increasing distance from the GM-WM interface.

These principles, however, were derived from gross techniques that were limited in their ability to delineate fiber bundles due to incomplete fixation and preservation of the white matter. Subsequent improvements in anatomical dissection followed the advent of the Klingler technique (Klingler, 1935), which involves repeated fixation and freezing to expand the white matter by forming ice crystals. This separation allowed for more complete white matter fixation and improved the visualization of white matter fiber bundles. Even so, the application of advanced dissection techniques has largely been limited to long association fiber tracts of the deep white matter (e.g., Sarubbo et al., 2013; Zemmoura et al., 2014, 2016; Hau et al., 2017; Yendiki et al., 2022), and the SWM is rarely studied using these methods, although more sophisticated dissection techniques and results are now beginning to be reported (e.g., Shinohara et al., 2020; Dannhoff et al., 2024; Shah et al., 2025, 2026).

There are two principal challenges in studying the neuroanatomy of the SWM. First, dissection cannot reliably locate the GM-WM border because it is a destructive technique that progresses from the pial surface downward through the cortex (but see Dannhoff et al., 2024). In addition, the progressive increase in myelination in the deep cortical layers (Ramón y Cajal, 1995) and the extensive network of intracortical layer 6 axons that run parallel to the cortical surface (e.g., Yamashita & Arikuni, 2001; Rockland & Knutson, 2001; Rockland, 2020) blur the distinction between the SWM and gray matter, so dissected bundles cannot be assigned with confidence to the SWM. Second, tract-tracing techniques have focused on long association or projection pathways and have not explicitly studied the SWM. While some studies have shown axons that travel in the SWM of the rhesus monkey brain (e.g., Cavada & Goldman-Rakic, 1989b; Seltzer et al., 1996; Morris et al., 1999; Petrides & Pandya, 2006; Muñoz-López et al., 2015), their depictions have been incidental and seldom discussed. This is also true for studies that show long-range axons in the superficial portions of the white matter adjacent to their termination, but either do not discuss the SWM at all (e.g., Jones et al., 1978; Seltzer & Pandya, 1989; Burton et al., 1995) or only mention it briefly (e.g., Friedman et al., 1986). As a result, no connectional study using experimentally labeled tract tracers has directly examined this region. Without the ability to follow specifically labeled fibers through the white matter, the organization and connectivity of the SWM remain largely assumed rather than empirically established.

We addressed this gap by analyzing archival case material from the laboratories of Dr. Deepak Pandya and Dr. Douglas Rosene (the Pandya-Rosene Neuroanatomical Archive). This archive consists of tract-tracing cases in the rhesus macaque monkey in which anterograde tracers were injected into specific cortical or subcortical areas. The primary autoradiographic slide material comprising this archive forms the basis of much of the literature of macaque connectional neuroanatomy (e.g., Rockland & Pandya, 1981; Mufson & Pandya, 1984; Petrides & Pandya, 1984, 2006; Demeter et al., 1985; Pandya & Rosene, 1985, 1993; Schmahmann & Pandya, 1997, 2006; Saunders & Rosene, 1988; Seltzer & Pandya, 1989; Barbas & Pandya, 1989; Cipolloni & Pandya, 1989; Pandya et al., 1994; Yeterian & Pandya, 1997, 2010; Morris et al., 1999; Blatt et al., 2003); however, the local projections within the SWM were not explicitly described in detail in these reports. To examine the organization and connectivity of the SWM and U-fibers, we selected 28 tract-tracing cases from the frontal, temporal, and parietal lobes of the rhesus macaque brain, as well as the posterior parietal lobe, often referred to as the preoccipital region (Yeterian & Pandya, 2010). A critical aspect of this approach was the ability to precisely define the GM-WM border by using Nissl counterstaining for each section, which allowed us to precisely determine the position of tract-tracing label in the SWM. This approach enabled direct evaluation of SWM organization using experimentally labeled fibers. We specifically asked: 1) whether U-fibers are a ubiquitous feature of sulcal white matter throughout the cortex, 2) whether U-fibers, when present, are symmetric and link matched points on adjacent gyri, 3) whether U-fibers occupy a single position in the superficial compartment of white matter and whether their depth reflects their length, as suggested by Meynert, and 4) whether U-fibers represent the only class of short association bundle beneath sulci, or whether other classes exist.

## 2. Results

### 2.1 Overview

Archival anterograde tract-tracing cases in the rhesus macaque monkey (n = 28) were evaluated for this study (see Methods). In each case, axonal bundles could be divided into several components (Schmahmann & Pandya, 2006). The most strongly labeled fibers comprise the cord (Figure 1, cord), a dense aggregation of fibers entering the white matter of the gyral core before separating into commissural (teal arrow) and subcortical bundles (green arrow). Label was also observed directly beneath the cerebral cortex in the superficial white matter (SWM), occasionally forming discrete bundles. These bundles were most compact and recognizable beneath sulci (Figure 1, red/white arrowheads), where they could adopt U-shapes. SWM labeling was typically continuous with focal patches of terminal label in the overlying cortical gray matter (Schmahmann & Pandya, 2006). Between the cord and SWM labeling were white matter regions of either little or no label, as well as label consistent with the position and characteristic stippled appearance of long-range pathways (Schmahmann & Pandya, 2006).

**Figure 1:**
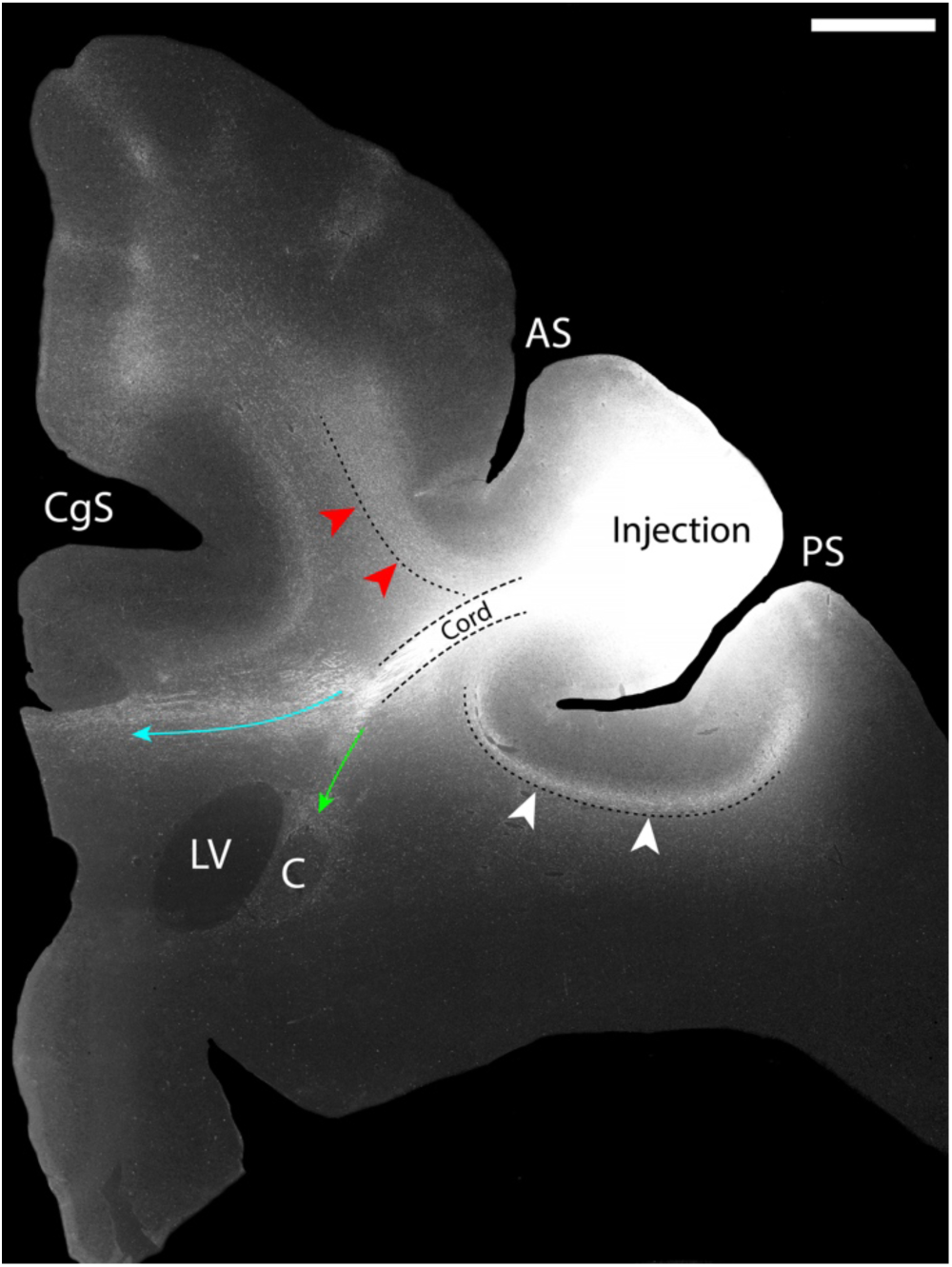
Overview of major axonal bundles after anterograde tracer injection in the cerebral cortex (imaged with darkfield microscopy). Radiolabeled amino acids were injected into dorsal area 9/46. Labeled cord fibers coursed from the injection site, then split into a commissural bundle to the corpus callosum (teal arrow) and a subcortical bundle (green arrow). Local association fibers (red and white arrowheads) also emanated from the injection site and traveled under sulci. Case BMFG. Scale bar: 2 mm. AS, arcuate sulcus; C, caudate nucleus; CgS, cingulate sulcus; LV, lateral ventricle; PS, principal sulcus.

Each section processed for autoradiography was Nissl-counterstained (Figure 2A-B) to reveal the cellular structure of the cerebral cortex. This facilitated the delineation of the gray matter-white matter (GM-WM) border under brightfield illumination (Figure 2C). This border was then transferred to the same section imaged under darkfield illumination to precisely delineate the position of fiber bundles relative to the GM-WM border (Figure 2D). Given the extensive ramifications of layer 6 intrinsic axonal systems (Rockland & Drash, 1996; Pucak et al., 1996; Yamashita & Arikuni, 2001; Rockland & Knutson, 2001; Rockland, 2020), this border is critical for accurately defining the SWM and its fibers.

**Figure 2:**
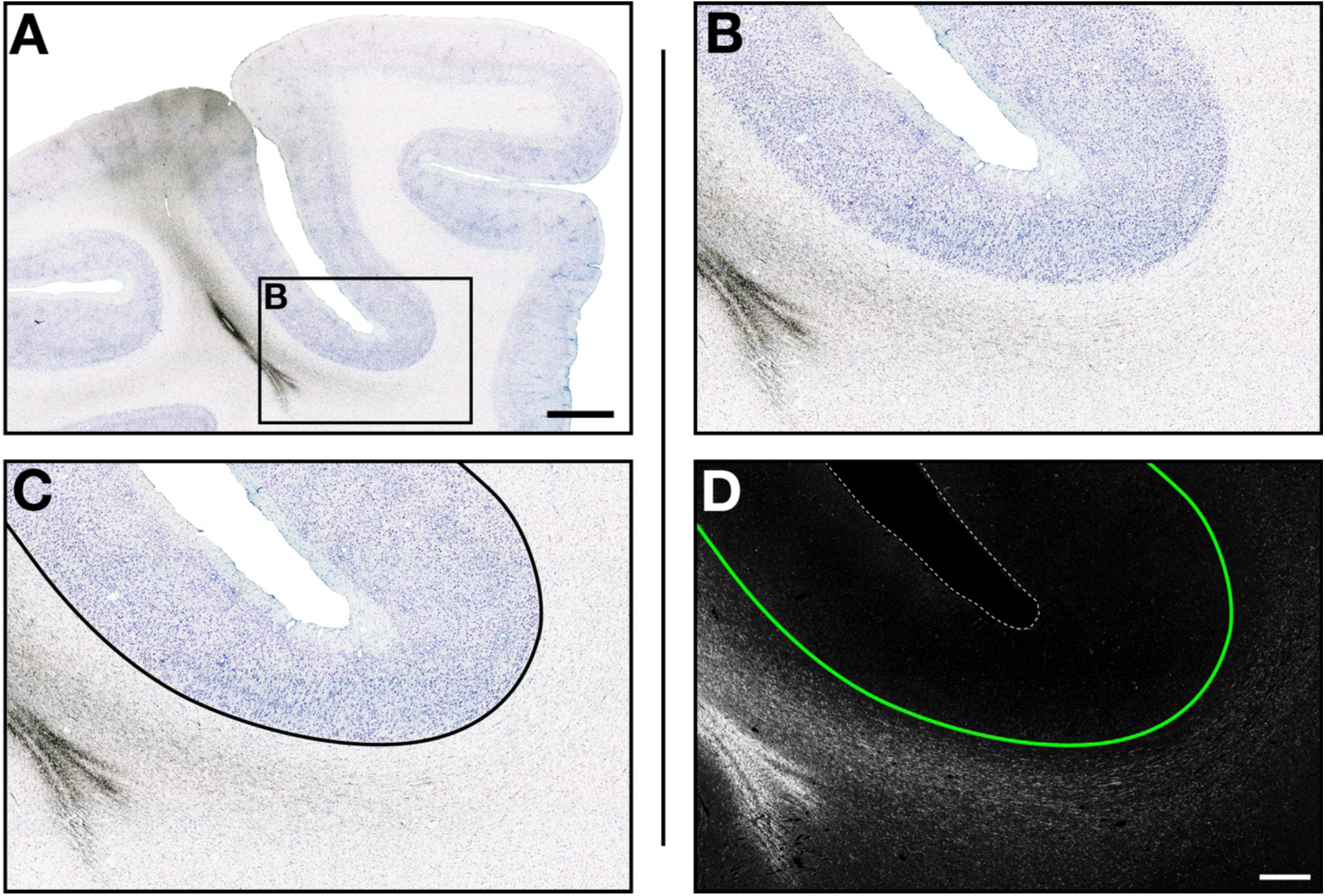
Delineation of the gray matter-white matter (GM-WM) border. A. Low-magnification brightfield image of the intraparietal sulcus in a coronal section processed for autoradiography (tracer, black) and Nissl substance (blue). B. Higher-magnification brightfield view of the box in A. C. Same field as in B with the GM-WM border indicated by the solid black line. D. Same field under darkfield illumination, with the GM-WM border from C shown by the solid green line; dashed white line represents the pial surface. Case BML. Scale bars: A = 2 mm; B-D = 500 µm.

### 2.2 U-fibers are not ubiquitous

The organization of the SWM was examined across major sulci of the macaque brain. Cases were selected from the archive in which tracer injections had been placed on gyri adjacent to four major sulci: the principal sulcus, central sulcus, intraparietal sulcus, and superior temporal sulcus. The 28 cases analyzed are listed in the methods section, and injection sites of the subset of cases presented in detail are shown in Figure 3. Detailed case-level descriptions of the those highlighted in Figure 3 are provided in the Supplementary Case Atlas.

**Figure 3:**
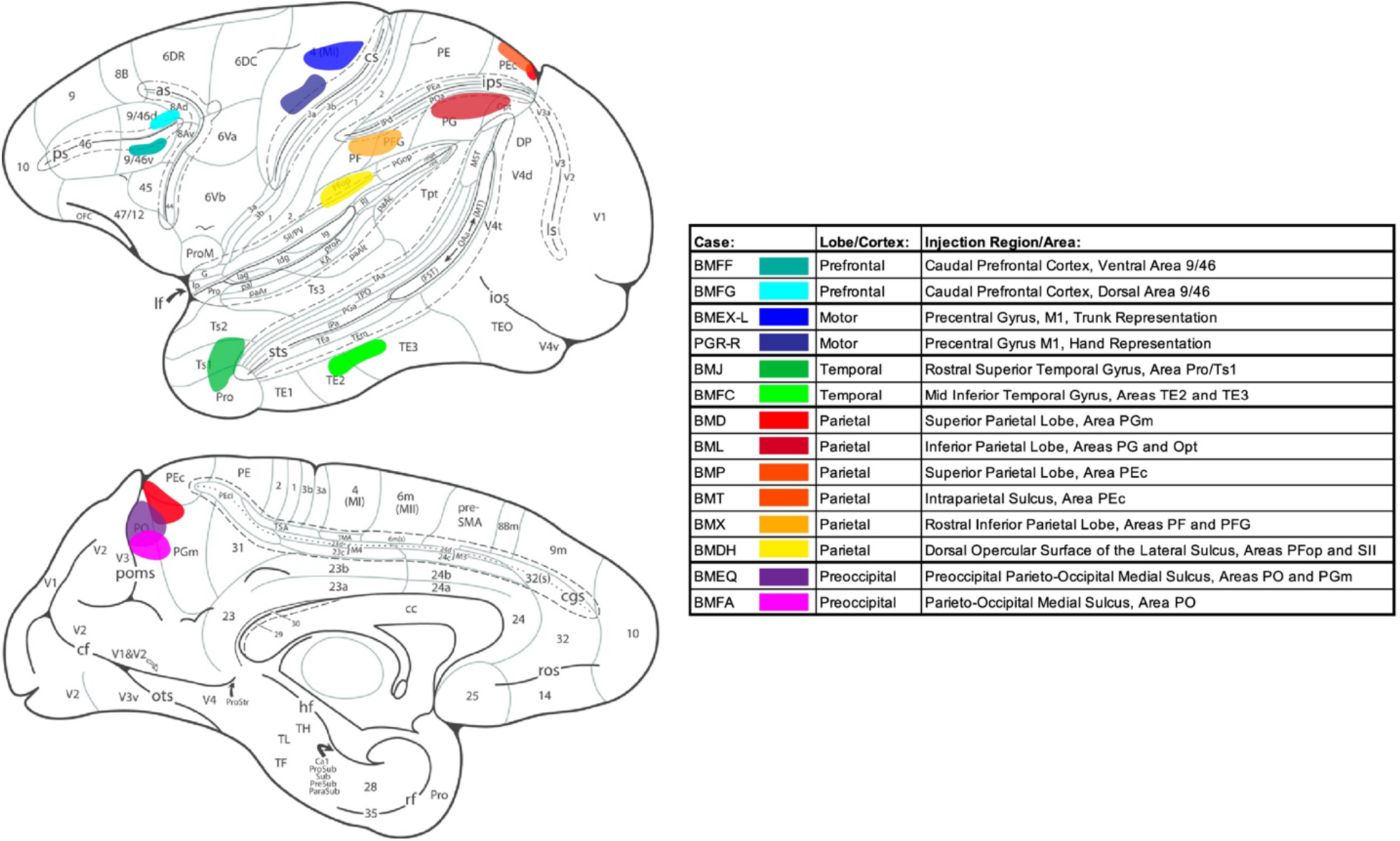
Subset of cases used in this study. Lateral (upper) and medial (lower) views of the rhesus macaque cerebral cortex showing cortical injection sites for 14 highlighted cases. Colored overlays indicate injection extent, and are color-coded by case, as shown in the table. Injections span the frontal, temporal, and parietal lobes, including gyri adjacent to major sulci (cs, central; ips, intraparietal; sts, superior temporal; ps, principal). The table lists each case ID, lobe, and injection region/area (gross designation then cytoarchitectonic area). Cortical surface diagram adapted from Morecraft et al. (2012); see Morecraft et al. (2012) for full abbreviations.

Short cortico-cortical association fibers were assessed in each case by tracing labeled fibers from the injection site to their terminations. In all cases, labeled short association fibers either: 1) formed distinct bundles that coursed along a sulcus adjacent to the injection, terminating along the sulcus and/or on the opposing gyrus, or 2) traveled as intragyral fibers that remained within the white matter of the injected gyrus and were associated with terminations along the gyrus rather than a sulcus. Tracing labeled bundles from the injection to their terminations revealed three distinct shapes (Figure 4A-C). U-fibers, consistent with classical anatomical descriptions (e.g., Meynert, 1885; Dejerine & Dejerine-Klumpke, 1895), were defined as labeled fibers forming bundles that arose from a gyral injection, coursed beneath a sulcus with a U-shaped trajectory, and terminated on the adjacent gyrus (Figure 4A). Two additional shorter association bundle types were identified that terminated along the sulcus rather than progressing to the adjacent gyrus: ‘L’-type connections, in which labeled fibers formed bundles that descended in the white matter along the proximal sulcal bank with a straight trajectory and terminated at or before the fundus (Figure 4B), and ‘J’-type connections, in which labeled bundles hooked around the fundus and terminated on the opposite sulcal bank without reaching the gyral crown (Figure 4C). Local association fibers formed distinct bundles along sulci consistent with one of the three shapes classified in 26 of the 28 cases (92.9%); in the remaining 2 cases, no short association bundles associated with sulci were observed and only intracortical fibers traveling predominantly within cortical layer 6 and/or intragyral fibers were present.

**Figure 4:**
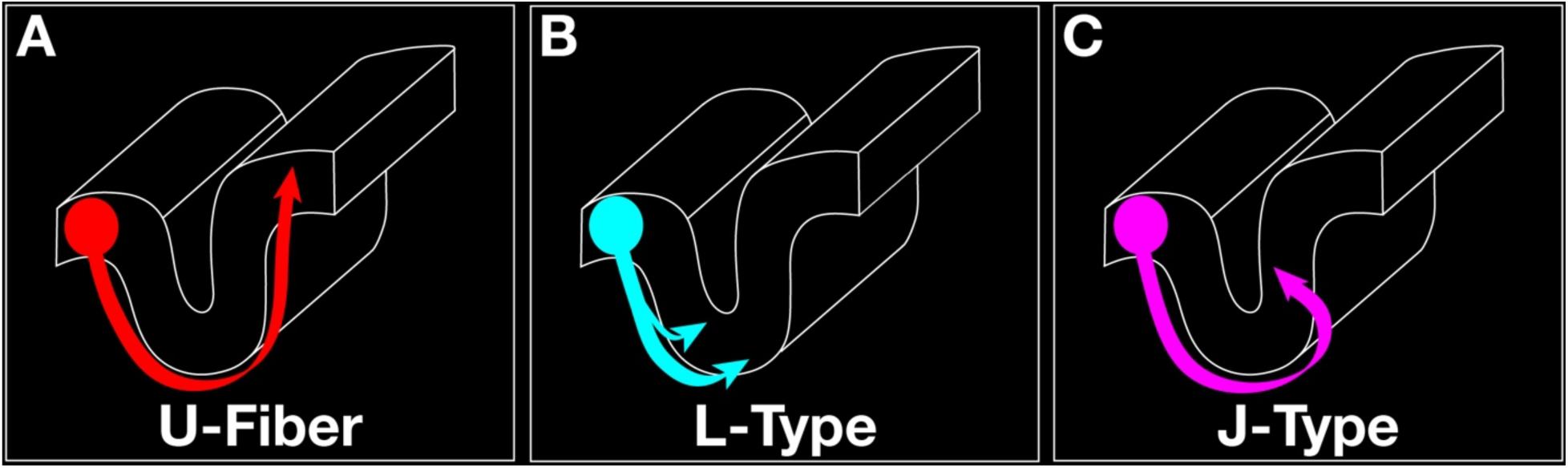
Schematic drawings of the three short association fiber bundle shapes. Each schematic shows bundles originating from a gyral injection (colored circle); this color scheme is retained in subsequent figures. A. U-shaped bundles (red) curve beneath the sulcus and terminate on the adjacent gyrus, consistent with U-fibers. B. L-type bundles (teal) follow a straight trajectory, terminating along the sulcal bank and fundus without crossing to the opposing sulcal bank. C. J-type bundles (magenta) curve beneath the sulcus and terminate on the opposing sulcal bank, but not on the adjacent gyrus.

The frequency of each bundle type was calculated for each sulcus and lobe injected (Tables 1-2). U-fibers were the least frequent, present beneath the major sulci of interest (principal, central, intraparietal, and superior temporal sulci) in only 10/28 cases (35.7%). When the criterion was broadened to include any U-fibers from the injections (and not just around the major sulci of interest), we observed additional U-fibers, increasing the total proportion of cases containing U-fibers to 14/28 (50%) (Table 1).

**TABLE 1.**
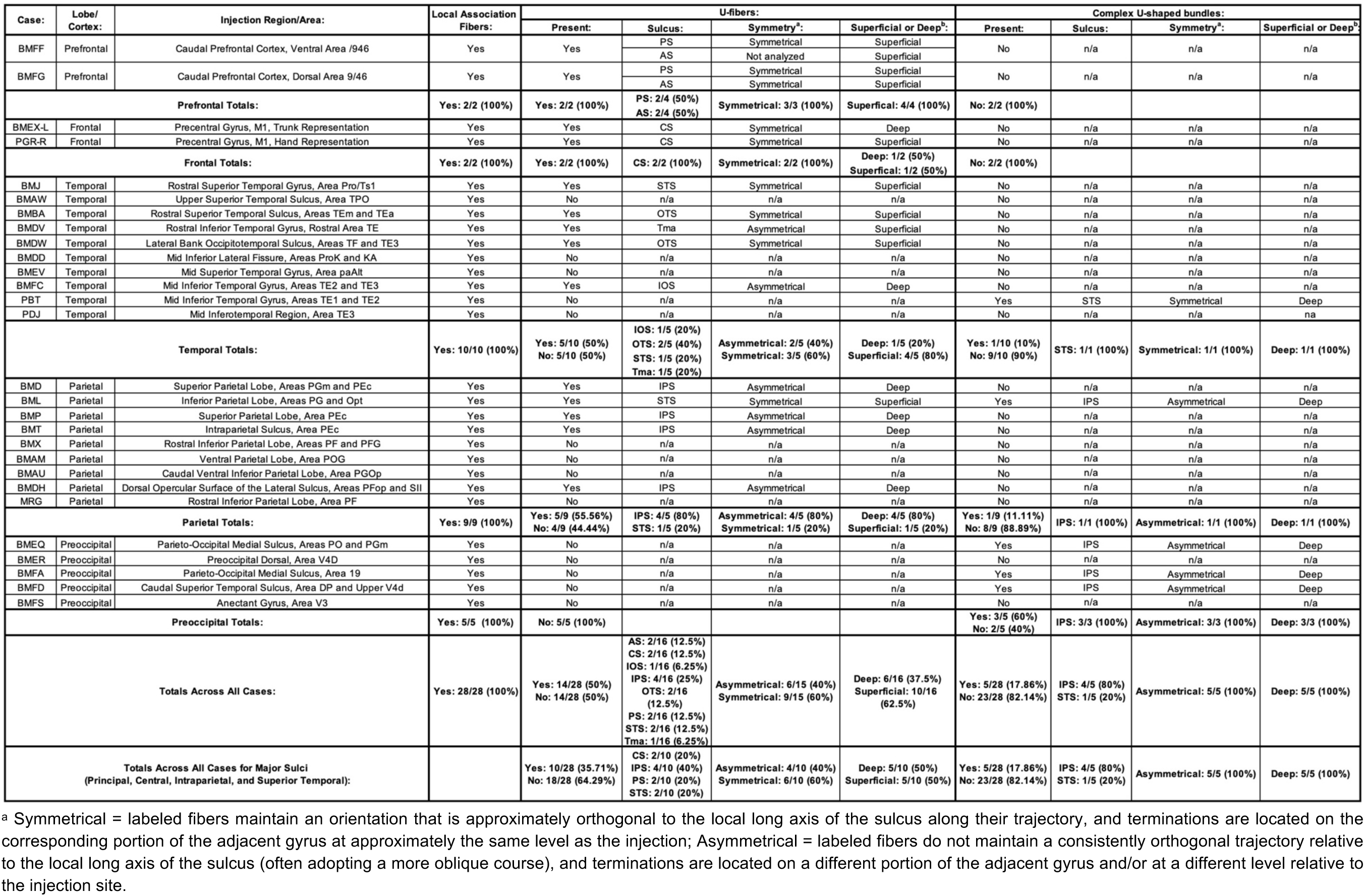
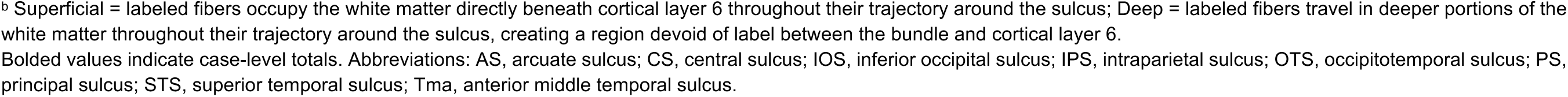
Catalog of U-fibers and complex U-shaped fiber bundles observed beneath sulci in all cases analyzed from the Pandya-Rosene Neuroanatomical Archive.

| Case: | Lobe/<br>Cortex: | Injection Region/Area: | Local Association<br>Fibers: | U-fibers: |  |  |  | Complex U-shaped bundles: |  |  |  |
| --- | --- | --- | --- | --- | --- | --- | --- | --- | --- | --- | --- |
|  |  |  |  | Present: | Sulcus: | Symmetry<sup>a</sup>: | Superficial or Deep<sup>b</sup>: | Present: | Sulcus: | Symmetry<sup>a</sup>: | Superficial or Deep<sup>b</sup>: |
| BMFF | Prefrontal | Caudal Prefrontal Cortex, Ventral Area /946 | Yes | Yes | PS | Symmetrical | Superficial | No | n/a | n/a | n/a |
|  |  |  |  |  | AS | Not analyzed | Superficial |  |  |  |  |
| BMFG | Prefrontal | Caudal Prefrontal Cortex, Dorsal Area 9/46 | Yes | Yes | PS | Symmetrical | Superficial | No | n/a | n/a | n/a |
|  |  |  |  |  | AS | Symmetrical | Superficial |  |  |  |  |
| Prefrontal Totals: |  |  | Yes: 2/2 (100%) | Yes: 2/2 (100%) | PS: 2/4 (50%)<br>AS: 2/4 (50%) | Symmetrical: 3/3 (100%) | Superficial: 4/4 (100%) | No: 2/2 (100%) |  |  |  |
| BMEX-L | Frontal | Precentral Gyrus, M1, Trunk Representation | Yes | Yes | CS | Symmetrical | Deep | No | n/a | n/a | n/a |
| PGR-R | Frontal | Precentral Gyrus, M1, Hand Representation | Yes | Yes | CS | Symmetrical | Superficial | No | n/a | n/a | n/a |
| Frontal Totals: |  |  | Yes: 2/2 (100%) | Yes: 2/2 (100%) | CS: 2/2 (100%) | Symmetrical: 2/2 (100%) | Deep: 1/2 (50%)<br>Superficial: 1/2 (50%) | No: 2/2 (100%) |  |  |  |
| BMJ | Temporal | Rostral Superior Temporal Gyrus, Area Pro/Ts1 | Yes | Yes | STS | Symmetrical | Superficial | No | n/a | n/a | n/a |
| BMAW | Temporal | Upper Superior Temporal Sulcus, Area TPO | Yes | No | n/a | n/a | n/a | No | n/a | n/a | n/a |
| BMBA | Temporal | Rostral Superior Temporal Sulcus, Areas TE<sub>m</sub> and TE<sub>a</sub> | Yes | Yes | OTS | Symmetrical | Superficial | No | n/a | n/a | n/a |
| BMDV | Temporal | Rostral Inferior Temporal Gyrus, Rostral Area TE | Yes | Yes | Tma | Asymmetrical | Superficial | No | n/a | n/a | n/a |
| BMDW | Temporal | Lateral Bank Occipitotemporal Sulcus, Areas TF and TE3 | Yes | Yes | OTS | Symmetrical | Superficial | No | n/a | n/a | n/a |
| BMDD | Temporal | Mid Inferior Lateral Fissure, Areas ProK and KA | Yes | No | n/a | n/a | n/a | No | n/a | n/a | n/a |
| BMEV | Temporal | Mid Superior Temporal Gyrus, Area paAlt | Yes | No | n/a | n/a | n/a | No | n/a | n/a | n/a |
| BMFC | Temporal | Mid Inferior Temporal Gyrus, Areas TE2 and TE3 | Yes | Yes | IOS | Asymmetrical | Deep | No | n/a | n/a | n/a |
| PBT | Temporal | Mid Inferior Temporal Gyrus, Areas TE1 and TE2 | Yes | No | n/a | n/a | n/a | Yes | STS | Symmetrical | Deep |
| PDJ | Temporal | Mid Inferotemporal Region, Area TE3 | Yes | No | n/a | n/a | n/a | No | n/a | n/a | na |
| Temporal Totals: |  |  | Yes: 10/10 (100%) | Yes: 5/10 (50%)<br>No: 5/10 (50%) | IOS: 1/5 (20%)<br>OTS: 2/5 (40%)<br>STS: 1/5 (20%)<br>Tma: 1/5 (20%) | Asymmetrical: 2/5 (40%)<br>Symmetrical: 3/5 (60%) | Deep: 1/5 (20%)<br>Superficial: 4/5 (80%) | Yes: 1/10 (10%)<br>No: 9/10 (90%) | STS: 1/1 (100%) | Symmetrical: 1/1 (100%) | Deep: 1/1 (100%) |
| BMD | Parietal | Superior Parietal Lobe, Areas PGM and PE<sub>c</sub> | Yes | Yes | IPS | Asymmetrical | Deep | No | n/a | n/a | n/a |
| BML | Parietal | Inferior Parietal Lobe, Areas PG and Opt | Yes | Yes | STS | Symmetrical | Superficial | Yes | IPS | Asymmetrical | Deep |
| BMP | Parietal | Superior Parietal Lobe, Area PE<sub>c</sub> | Yes | Yes | IPS | Asymmetrical | Deep | No | n/a | n/a | n/a |
| BMT | Parietal | Intraparietal Sulcus, Area PE<sub>c</sub> | Yes | Yes | IPS | Asymmetrical | Deep | No | n/a | n/a | n/a |
| BMX | Parietal | Rostral Inferior Parietal Lobe, Areas PF and PFG | Yes | No | n/a | n/a | n/a | No | n/a | n/a | n/a |
| BMAM | Parietal | Ventral Parietal Lobe, Area POG | Yes | No | n/a | n/a | n/a | No | n/a | n/a | n/a |
| BMAU | Parietal | Caudal Ventral Inferior Parietal Lobe, Area PGO<sub>p</sub> | Yes | No | n/a | n/a | n/a | No | n/a | n/a | n/a |
| BMDH | Parietal | Dorsal Opercular Surface of the Lateral Sulcus, Areas PFO<sub>p</sub> and SII | Yes | Yes | IPS | Asymmetrical | Deep | No | n/a | n/a | n/a |
| MRG | Parietal | Rostral Inferior Parietal Lobe, Area PF | Yes | No | n/a | n/a | n/a | No | n/a | n/a | n/a |
| Parietal Totals: |  |  | Yes: 9/9 (100%) | Yes: 5/9 (55.56%)<br>No: 4/9 (44.44%) | IPS: 4/5 (80%)<br>STS: 1/5 (20%) | Asymmetrical: 4/5 (80%)<br>Symmetrical: 1/5 (20%) | Deep: 4/5 (80%)<br>Superficial: 1/5 (20%) | Yes: 1/9 (11.11%)<br>No: 8/9 (88.89%) | IPS: 1/1 (100%) | Asymmetrical: 1/1 (100%) | Deep: 1/1 (100%) |
| BMEQ | Preoccipital | Parieto-Occipital Medial Sulcus, Areas PO and PG<sub>m</sub> | Yes | No | n/a | n/a | n/a | Yes | IPS | Asymmetrical | Deep |
| BMER | Preoccipital | Preoccipital Dorsal, Area V4D | Yes | No | n/a | n/a | n/a | No | n/a | n/a | n/a |
| BMFA | Preoccipital | Parieto-Occipital Medial Sulcus, Area 19 | Yes | No | n/a | n/a | n/a | Yes | IPS | Asymmetrical | Deep |
| BMFD | Preoccipital | Caudal Superior Temporal Sulcus, Area DP and Upper V4d | Yes | No | n/a | n/a | n/a | Yes | IPS | Asymmetrical | Deep |
| BMFS | Preoccipital | Anectant Gyrus, Area V3 | Yes | No | n/a | n/a | n/a | No | n/a | n/a | n/a |
| Preoccipital Totals: |  |  | Yes: 5/5 (100%) | No: 5/5 (100%) |  |  |  | Yes: 3/5 (60%)<br>No: 2/5 (40%) | IPS: 3/3 (100%) | Asymmetrical: 3/3 (100%) | Deep: 3/3 (100%) |
| Totals Across All Cases: |  |  | Yes: 28/28 (100%) | Yes: 14/28 (50%)<br>No: 14/28 (50%) | AS: 2/16 (12.5%)<br>CS: 2/16 (12.5%)<br>IOS: 1/16 (6.25%)<br>IPS: 4/16 (25%)<br>OTS: 2/16 (12.5%)<br>PS: 2/16 (12.5%)<br>STS: 2/16 (12.5%)<br>Tma: 1/16 (6.25%) | Asymmetrical: 6/15 (40%)<br>Symmetrical: 9/15 (60%) | Deep: 6/16 (37.5%)<br>Superficial: 10/16 (62.5%) | Yes: 5/28 (17.86%)<br>No: 23/28 (82.14%) | IPS: 4/5 (80%)<br>STS: 1/5 (20%) | Asymmetrical: 5/5 (100%) | Deep: 5/5 (100%) |
| Totals Across All Cases for Major Sulci<br>(Principal, Central, Intraparietal, and Superior Temporal): |  |  |  | Yes: 10/28 (35.71%)<br>No: 18/28 (64.29%) | CS: 2/10 (20%)<br>IPS: 4/10 (40%)<br>PS: 2/10 (20%)<br>STS: 2/10 (20%) | Asymmetrical: 4/10 (40%)<br>Symmetrical: 6/10 (60%) | Deep: 5/10 (50%)<br>Superficial: 5/10 (50%) | Yes: 5/28 (17.86%)<br>No: 23/28 (82.14%) | IPS: 4/5 (80%)<br>STS: 1/5 (20%) | Asymmetrical: 5/5 (100%) | Deep: 5/5 (100%) |

<sup>a</sup> Symmetrical = labeled fibers maintain an orientation that is approximately orthogonal to the local long axis of the sulcus along their trajectory, and terminations are located on the corresponding portion of the adjacent gyrus at approximately the same level as the injection; Asymmetrical = labeled fibers do not maintain a consistently orthogonal trajectory relative to the local long axis of the sulcus (often adopting a more oblique course), and terminations are located on a different portion of the adjacent gyrus and/or at a different level relative to the injection site.
<sup>b</sup> Superficial = labeled fibers occupy the white matter directly beneath cortical layer 6 throughout their trajectory around the sulcus; Deep = labeled fibers travel in deeper portions of the white matter throughout their trajectory around the sulcus, creating a region devoid of label between the bundle and cortical layer 6.
Bolded values indicate case-level totals. Abbreviations: AS, arcuate sulcus; CS, central sulcus; IOS, inferior occipital sulcus; IPS, intraparietal sulcus; OTS, occipitotemporal sulcus; PS, principal sulcus; STS, superior temporal sulcus; Tma, anterior middle temporal sulcus.

**TABLE 2.**
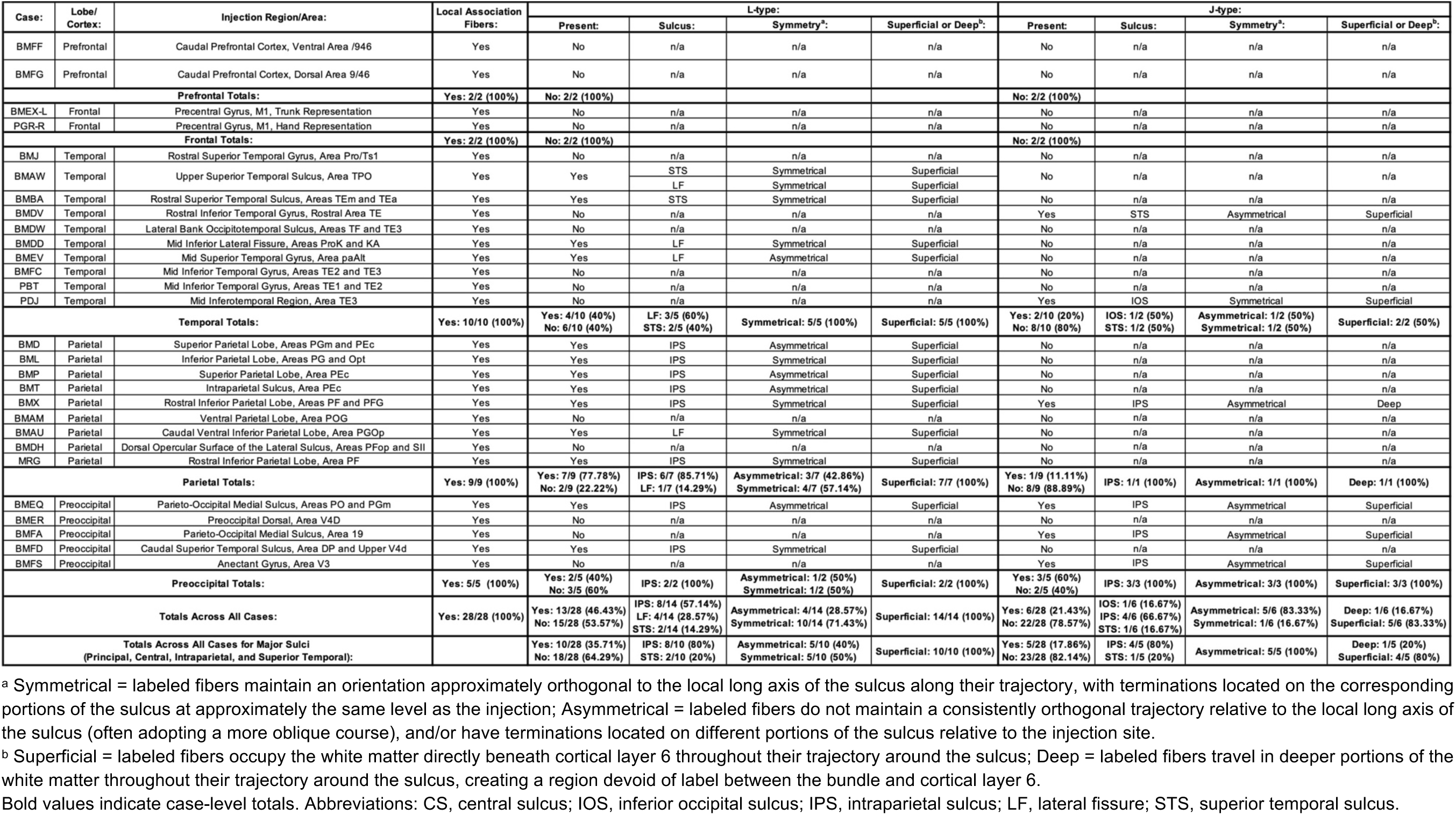
Catalog of ‘L’- and ‘J’-type connections observed beneath sulci in all cases analyzed from the Pandya-Rosene Neuroanatomical Archive.

The prevalence of U-fibers around major sulci varied by the lobe injected, with the highest rate in the frontal lobe (4/4; 100%), followed by parietal (5/9; 55.6%), temporal (1/10; 10%), and preoccipital (0/5; 0%) cortices (Table 1). In contrast, the shorter L- or J-type connections were observed more frequently, present in 60.7% (17/28) of cases overall, and were more often observed along any sulcus in the parietal (7/9; 77.8%), temporal (7/10; 70%) and pre-occipital (4/5; 80%) cortices than in the frontal lobe (0/4; 0%) (Table 2). Thus, these L- and J-type connections were associated with deeper sulci such as the intraparietal sulcus, superior temporal sulcus, and lateral (Sylvian) fissure. Individual cases often contained more than one bundle type (e.g., U-fibers with L- or J-type connections, or L- and J-type connections together), with L-type connections present in 46.4% (13/28) of cases along any sulcus, and J-type connections in 21.4% (6/28); these proportions are therefore not mutually exclusive (Tables 1-2). Representative cases from each lobe, shown in Figures 5-9, illustrate the organization of these fiber bundle types across the major sulci examined.

**Figure 5:**
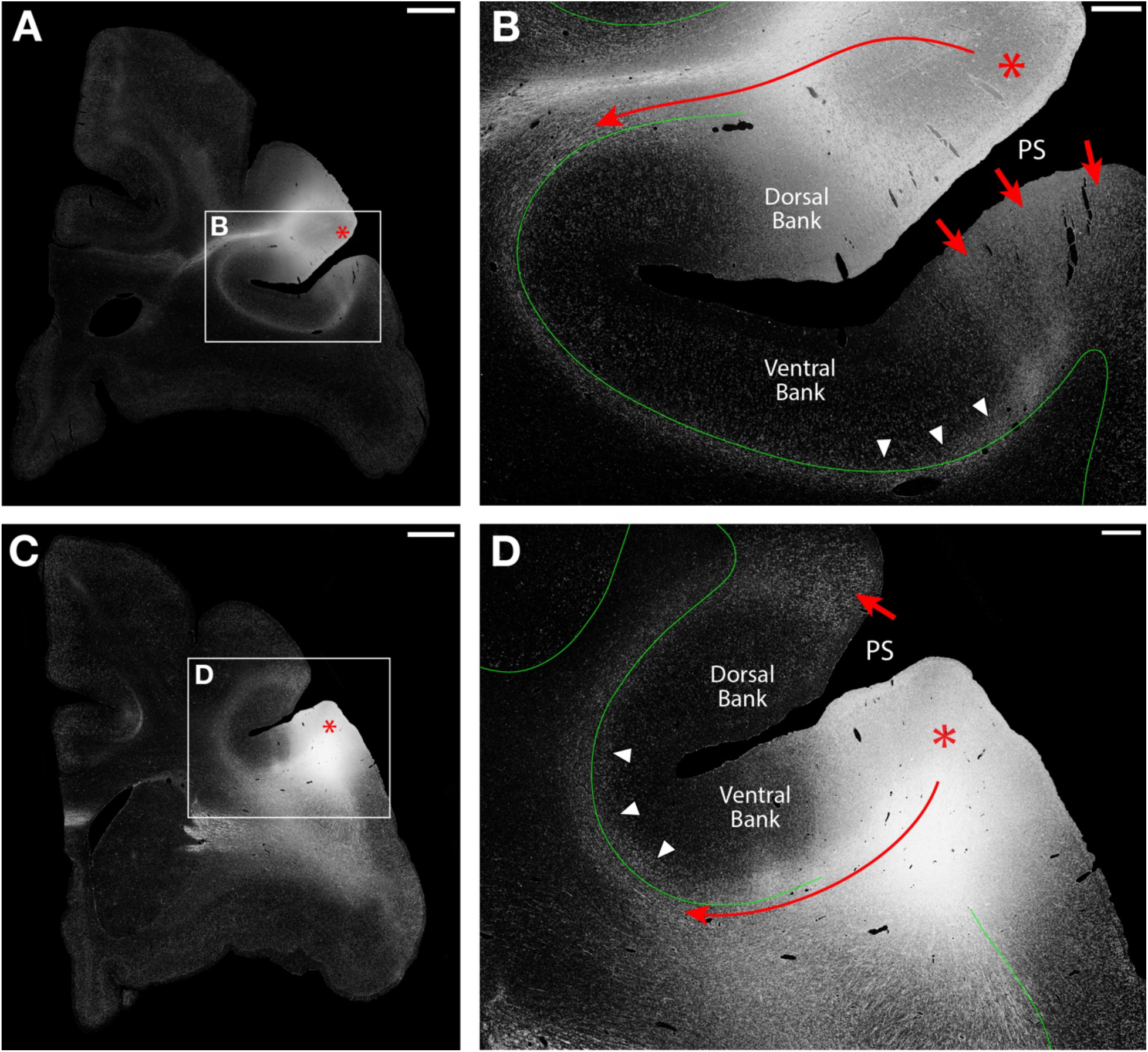
Autoradiographs of short association fibers in the macaque prefrontal cortex (imaged with darkfield microscopy). A. Low-magnification image following an injection in dorsal area 9/46 (red asterisk). B. Higher-magnification view of A; a U-fiber (solid red line) courses beneath the principal sulcus (PS) in the white matter immediately deep to the gray matter-white matter border (solid green line); labeling extends across the border into cortex (white arrowheads) before forming columnar terminations on the ventral PS bank and gyrus (red arrows). C. Low-magnification view of an injection in ventral area 9/46 (red asterisk). D. Higher-magnification view of C: a U-fiber (solid red line) courses beneath the PS, with label extending into cortex (white arrowheads) before terminating in a columnar fashion on the adjacent gyral crown (red arrow). A-B, case BMFG; C-D, case BMFF. Scale bars: A, C = 2 mm; B, D = 500 µm.

### Prefrontal Cortex

In both prefrontal cases (BMFG, BMFF), injections on the gyri dorsal or ventral to the principal sulcus produced prominent labeled U-shaped bundles (Figure 5B, D; solid red lines) that coursed beneath the sulcus and terminated on the opposing gyrus (Figure 5B, D; red arrows). Thus, both labeled bundles met the criteria for U-fibers and were classified accordingly. In both cases, labeling was also present in cortical layer 6 and was continuous with label in the underlying white matter (Figure 5B, D; arrowheads), a finding that highlights the importance of delineating the GM-WM border (Figure 5B, D; solid green line).

### Motor Cortex

Labeled fibers consistent with U-shaped bundles were identified beneath the central sulcus in both motor cortex cases (PGR-R and BMEX-L; Figure 6). Because the central sulcus is oriented dorsoventrally, labeled fibers were followed across coronal sections as they curved beneath the sulcal fundus (Figure 6B, D; dashed red lines) and continued into the postcentral gyral white matter. In both cases, the organization of labeled bundles was more complex than that observed along the principal sulcus, described above. In PGR-R, labeled fibers formed a U-shaped bundle associated with label in the lower cortical layers at the fundus (Figure 6B, white arrowheads) that was continuous with label in the underlying white matter. Labeled fibers then entered the postcentral white matter and fanned out before terminating on the caudal bank of the central sulcus (Figure 6B, red arrows) and further caudally, on the postcentral gyral crown. Because this bundle had a U-shaped trajectory and interconnected opposing gyri, it was classified as a U-fiber. In BMEX-L, label beneath the central sulcus resolved into two bands of differing intensity (Figure 6D; Supplementary Figure 3D). The deeper band consisted of labeled fibers forming a U-shaped bundle (Figure 6D, dashed red line) that coursed caudally around the central sulcus, terminating on the opposing gyrus (red arrows), consistent with a U-fiber. The more superficial band coursed predominantly within cortical layer 6 along the banks and fundus of the central sulcus (Figure 6D, white arrowheads), extending only slightly into the subjacent white matter.

**Figure 6:**
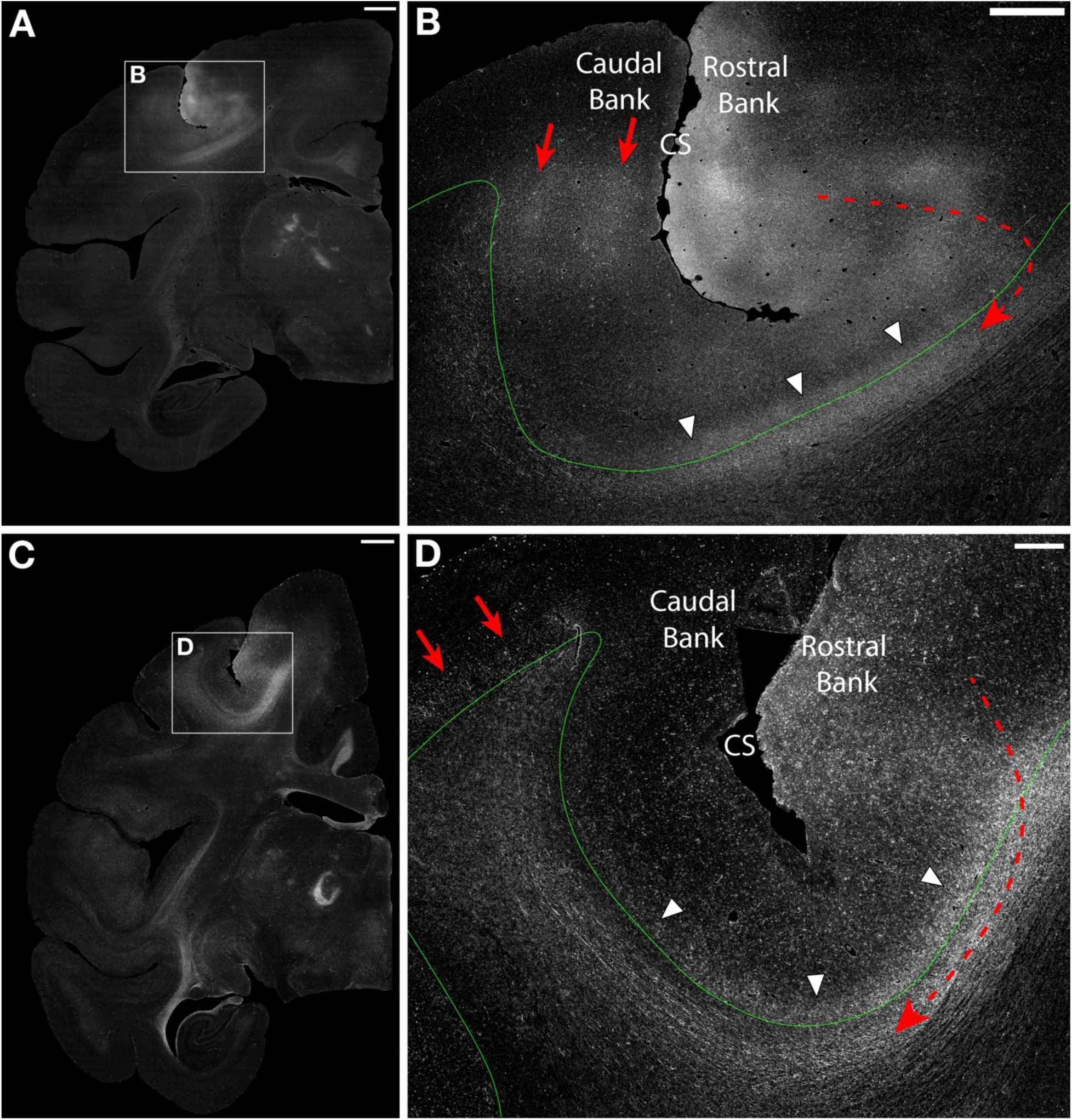
Autoradiographs of short association fibers in peri-Rolandic (motor/somatosensory) cortex (imaged with darkfield microscopy). A. Low-magnification image ∼4 mm caudal to an injection in the M1 hand representation (precentral gyrus). B. Higher-magnification view of A; a U-shaped bundle (dashed red line) courses caudally around the central sulcus (CS) deep to the gray matter-white matter (GM-WM) border (solid green line); this bundle was associated with label in the lower cortical layers at the fundus (white arrowheads) before fanning out and forming columnar terminations on the caudal CS bank (red arrows) and further caudally on the postcentral gyrus. Based on its gyrus-to-gyrus connectivity, this bundle was classified as a U-fiber. C. Low-magnification image ∼6 mm caudal to an injection in the M1 trunk representation. D. Higher-magnification view of C: two distinct intensity bands of labeling beneath the CS are observed; an intracortical band within cortical layer 6 along the sulcal banks and fundus (white arrowheads), and a U-fiber (dashed red line) with terminations on the postcentral gyrus (red arrows). A-B, case PGR-R; C-D, case BMEX-L (reflected for comparison with PGR-R). Scale bars: A, C = 2 mm; B = 1 mm; D = 500 µm.

### Parietal Lobe and Preoccipital Region

Along the intraparietal sulcus (IPS), U-shaped fiber bundles linking the superior (SPL) and inferior parietal lobules (IPL) were identified in only 4 of 14 cases (28.6%) (Table 1). At the rostral IPS, where the sulcus is shallow, U-shaped bundles were observed in only one of three cases (BMDH; Figure 7A-B). In BMDH, labeled fibers formed a U-shaped bundle (Figure 7B, solid red line; outlined with dashed red lines) that passed beneath the IPS fundus, entered the SPL, and terminated on the medial IPS bank and on the adjoining gyrus (red arrows). Therefore, this fiber bundle was classified as a U-fiber. In the other two rostral IPS cases (BMX, MRG), fibers terminated along the sulcal banks or fundus without reaching the opposing gyrus, consistent with L- and J-type connections. In case BMX, a superficial bundle (Figure 7D, solid teal line) coursed directly beneath the GM-WM border and terminated on the lateral IPS bank and at the fundus (teal arrowheads), consistent with an L-type connection. A second deeper bundle (Figure 7D, solid magenta line; outlined with dashed magenta lines) continued around the fundus and terminated on the medial bank of the IPS (magenta arrow), consistent with a J-type connection. In this case, the L- and J-type connections were observed together, and the J-type was deeper than the L-type. When the L-type bundle ended, the J-type bundle maintained its deeper position beneath the GM-WM border, continuing around the IPS fundus before passing through the more superficial portions of the white matter to terminate on the medial IPS bank.

**Figure 7:**
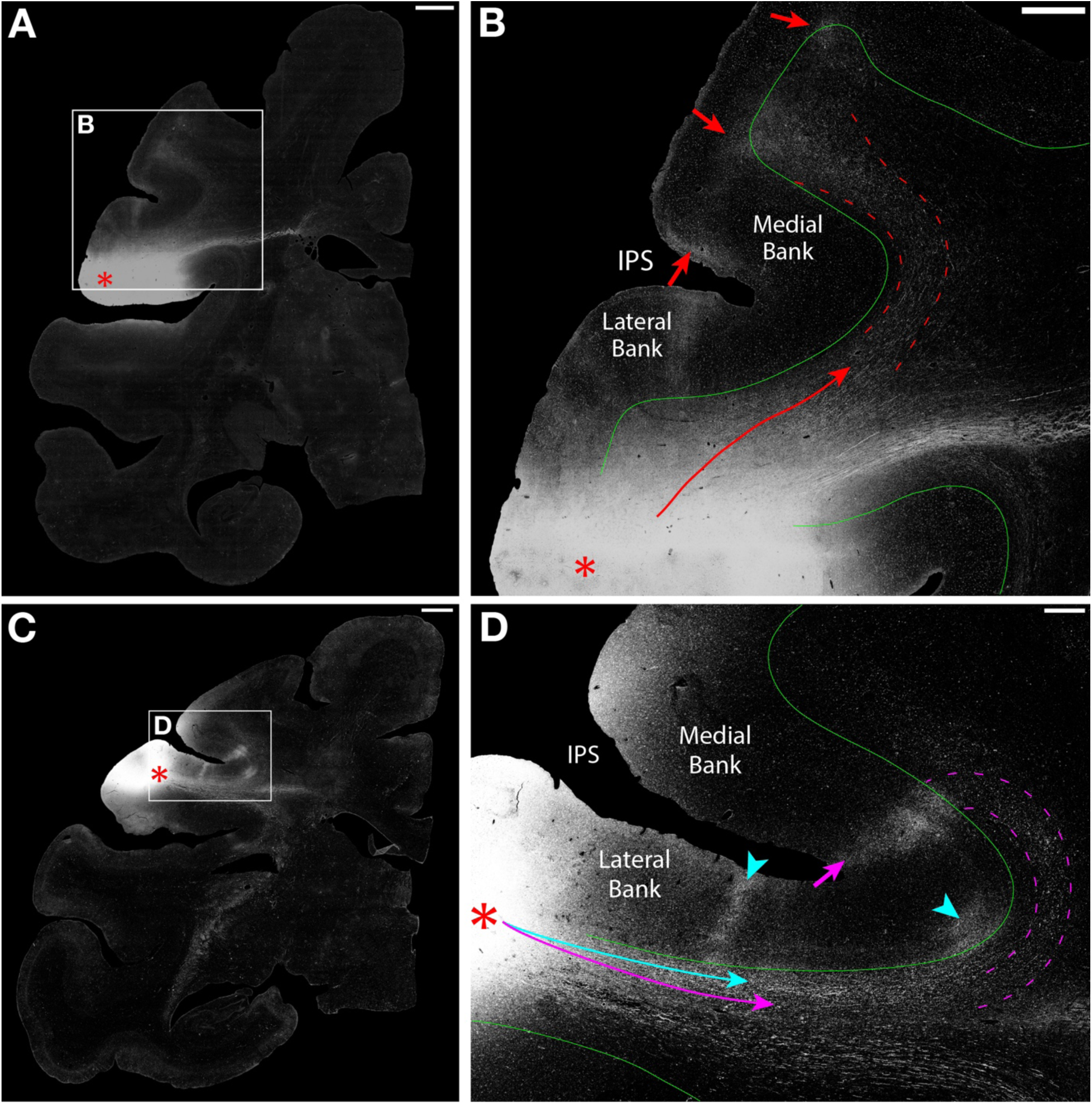
Autoradiographs of short association fibers in the macaque rostral parietal lobe (imaged under darkfield). A. Low-magnification image following an injection in the rostral inferior parietal lobule (IPL) on the dorsal opercular surface of the lateral sulcus (red asterisk). B. Higher-magnification view of A; a U-shaped bundle (solid red line; outlined by dashed red lines) courses beneath the intraparietal sulcus (IPS) in deeper portions of the white matter, largely avoiding the zone beneath the gray matter-white matter (GM-WM) border (solid green line); after crossing the fundus, labeling extends into cortex on the medial IPS bank and superior parietal lobule (SPL) crown (red arrows). C. Low-magnification image following an injection in the rostral IPL (red asterisk). D. Higher-magnification view of C: an L-type bundle (solid teal line) courses from the injection immediately deep to the GM-WM border (solid green line), terminating along the lateral IPS bank and fundus (teal arrowheads); a deeper J-type bundle (solid magenta line; outlined by dashed magenta lines) travels along the lateral IPS bank, hooks around the sulcus, and terminates on the medial IPS bank (magenta arrow). A-B, case BMDH; C-D, case BMX. Scale bars: A, C = 2 mm; B = 1 mm; D = 500 µm.

At the mid-IPS, where the sulcus is deeper and contains multiple distinct cortical areas, U-shaped fiber bundles were identified in all three cases with SPL injections (BMD, BMP, BMT; Figure 8). Labeled fibers coursed rostrally from the injection site through the SPL white matter, curved ventrally around the depth of the IPS with a U-shaped trajectory (Figure 8B, dashed red line), entered the IPL white matter, and ascended dorsally to terminate on the gyral crown (red arrow). As these fiber bundles had a U-shaped configuration beneath the IPS and interconnected adjacent gyri, they were classified as U-fibers in all three SPL cases. In contrast to the U-fibers, which avoided the white matter directly beneath the GM-WM border, shorter fiber bundles consistent with L-type connections (Figure 8B, dashed teal line) were also observed in all three SPL cases, coursing directly below the GM-WM border and terminating along the medial IPS bank (teal arrowheads). No U-fibers were observed in the three mid-IPS cases with IPL injections or in the five preoccipital cases (Table 1). However, shorter L- and J-type connections, similar to those in the SPL cases, were present in these cases (Table 2).

**Figure 8:**
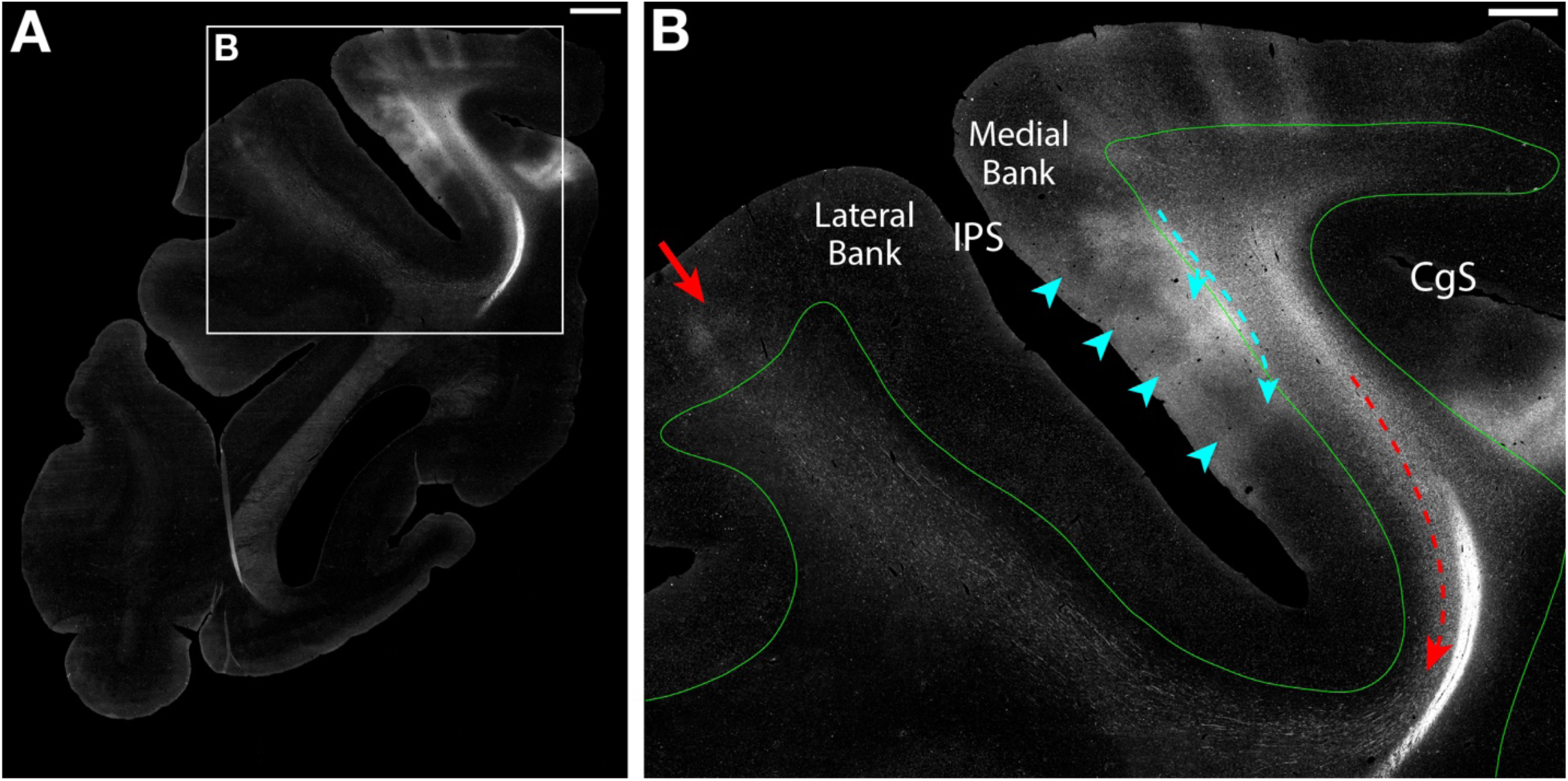
Autoradiographs of short association fibers in the macaque parietal lobe (imaged with darkfield microscopy). A. Low-magnification coronal section ∼4 mm rostral to a superior parietal lobule injection. B. Higher-magnification view of A; a U-shaped bundle (dashed red line) courses ventrally around the intraparietal sulcus (IPS) in deeper portions of the white matter and terminates on the crown of the inferior parietal lobule (red arrow); an L-type bundle (dashed teal line) courses beneath the gray matter-white matter border (solid green line), terminating along the medial IPS bank (teal arrowheads). Case BMP. Scale bars: A = 2 mm; B = 1 mm. CgS, cingulate sulcus.

### Temporal Lobe

Labeled fibers forming U-shaped bundles beneath the superior temporal sulcus (STS) were rare and present in only 1 of 10 temporal cases (10%) (BMJ; Figure 9). In BMJ, labeled fibers formed a U-shaped bundle (Figure 9B, solid red line) that coursed ventrally around the rostral STS and terminated on the opposing sulcal bank and inferior temporal gyrus (red arrows), consistent with a U-fiber. In the remaining nine temporal cases, short association bundles terminated more often along the sulcal banks or fundus of the STS, rather than reaching the adjacent gyrus, consistent with shorter L- and J-type bundles (Table 2). STS labeling was, however, observed in one parietal case (BML) with an injection in the caudal IPL near the junction of the STS and IPS (see Methods and Figure 3). In this case, labeled fibers curved around the caudal STS and terminated on the adjacent gyrus, consistent with a U-fiber. Although temporal cases were selected for injections on the gyri adjacent to the STS, 4 of 6 cases (66.7%) with inferotemporal injections produced U-shaped bundles that coursed around other neighboring sulci rather than the STS and terminated on the adjacent gyrus, consistent with U-fibers (Table 1).

**Figure 9:**
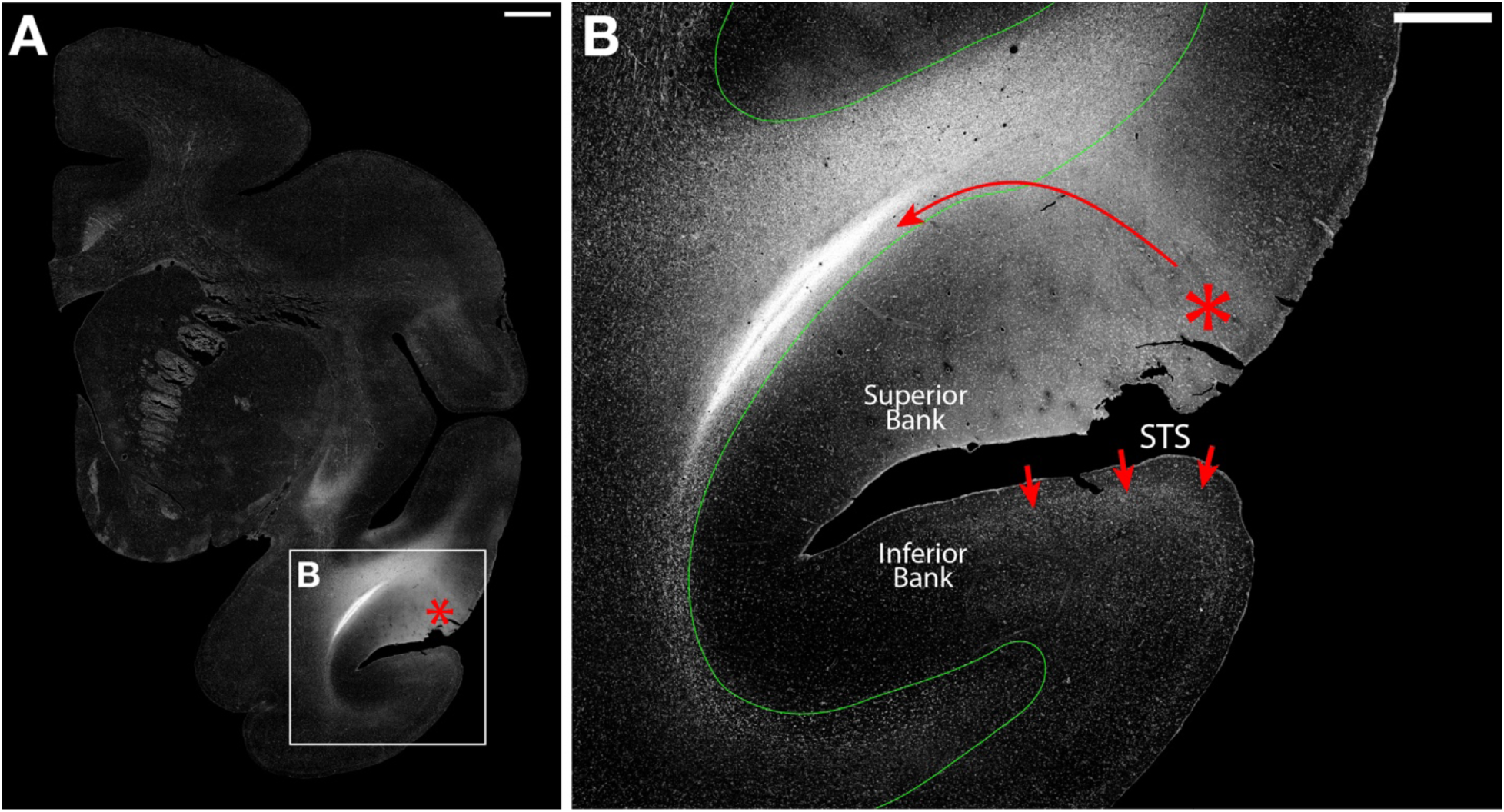
Autoradiographs of short association fibers in the rostral temporal lobe (imaged with darkfield microscopy). A. Low-magnification image following an injection in the rostral superior temporal gyrus (red asterisk). B. Higher-magnification view of A; a U-fiber (solid red line) courses ventrally around the rostral portion of the superior temporal sulcus (STS) within the white matter directly beneath the gray matter-white matter border (solid green line) and forms terminations along the inferior STS bank and inferior temporal gyral crest (red arrows). Case BMJ. Scale bars: A = 2 mm; B = 1 mm.

### Summary

Overall, these observations indicate that U-fibers are neither ubiquitous nor uniformly distributed across cortical sulci and their incidence was sulcal-dependent. Furthermore, the identification of association fibers shorter than U-fibers (e.g., L- and J-type connections) demonstrates that the sulcal white matter contains multiple classes of short association fiber bundles rather than a single canonical U-fiber system.

### 2.3 U-fiber Trajectories do not Always Reflect Sulcal Geometry

Meynert’s classical anatomical descriptions of U-fibers as “wire rings” (Meynert, 1885, p. 38) have been taken to mean that U-fibers are arranged along the length of a sulcus in a largely symmetrical fashion and interconnect corresponding portions of opposing gyri (Figure 10A). To investigate this organizational principle in tract-tracing material, fiber bundle symmetry was evaluated in the 14 cases with U-fibers by examining the position of the injection site and terminations on the gyri, as well as the trajectory of labeled fibers throughout their course around the sulcus. Symmetrical U-fibers (Figure 10A) were defined as bundles that: 1) maintained an orientation approximately orthogonal to the local long axis of the sulcus along their trajectory; 2) terminated on the corresponding portion of the opposing gyrus at approximately the same level as the injection; and 3) maintained a consistent orientation throughout the bundle’s trajectory around the sulcus. Asymmetrical bundles (Figure 10B-E) violated one or more of those criteria. Evaluation of fiber bundle symmetry showed that U-fibers did not always traverse sulci in the symmetrical fashion that Meynert predicted. Overall, 60% (9/15) of all U-fibers were symmetrical and 40% (6/15) were asymmetrical, a proportion that remained unchanged when considering only bundles beneath the major sulci (Table 1). The distribution was not uniform across the cortex: symmetrical bundles were observed in all U-fibers around the principal sulcus (2/2), central sulcus (2/2), and STS (2/2), whereas all U-fibers along the IPS (4/4) were asymmetrical.

**Figure 10:**
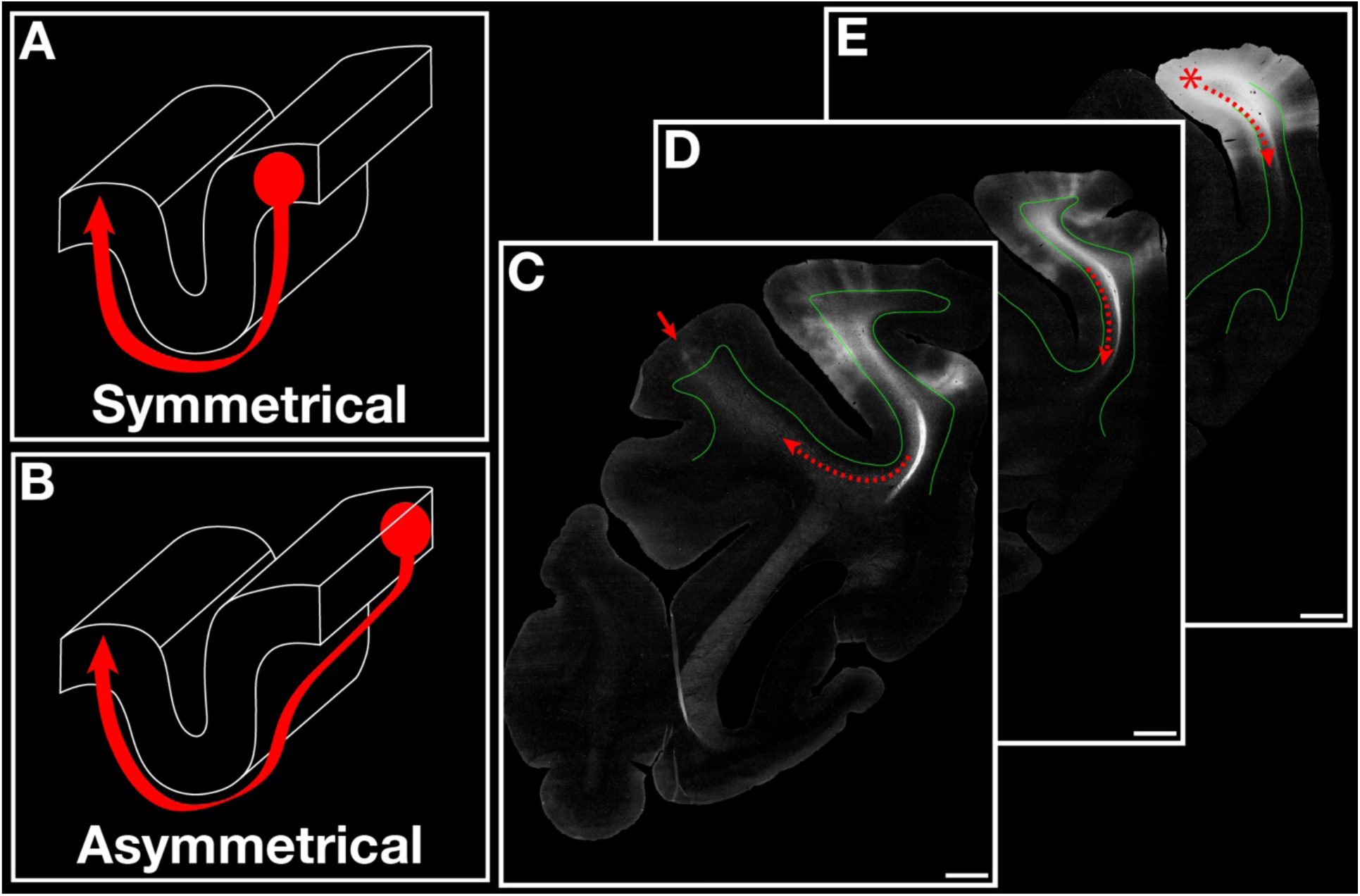
Symmetric versus asymmetric U-fibers. A. Schematic of a symmetric U-fiber terminating on the corresponding portion of the adjacent gyrus with a trajectory approximately orthogonal to the local long axis of the sulcus. B. Schematic of an asymmetric U-fiber terminating on a different portion of the adjacent gyrus relative to the injection site (red circle) with an oblique trajectory. C-E. Coronal sections through the posterior parietal lobe, rostral (C) to caudal (E), showing a rostrally directed asymmetric labeled U-fiber (dashed red line). In E, it descends within the superior parietal lobule white matter from the gyral injection (red asterisk); in D it curves beneath the intraparietal sulcus fundus; in C it completes the turn, enters the inferior parietal lobule gyral white matter, and proceeds to its termination (red arrow) at a different rostrocaudal level than the injection. Case BMP. Scale bars: C-E = 2 mm.

The U-fibers identified in the principal sulcus cases (BMFG, BMFF) illustrate U-fibers with a symmetric organization. In both cases, labeled fibers maintained a consistent orientation orthogonal to the long axis of the sulcus and terminated on the corresponding portion of the adjacent gyrus (Figure 5B, D; red arrows) at approximately the same rostrocaudal level as the injection (Figure 5B, D; red asterisks). Labeled fibers within these U-fibers consisted of dense, longitudinally oriented fibers beneath the proximal sulcal bank and fundus and gradually fanned out beneath the opposing bank before they entered the cortical gray matter (Figure 14B, D). Both U-fibers observed beneath the central sulcus (PGR-R, BMEX-L) were also classified as symmetric; however, because the central sulcus is oriented dorsoventrally rather than rostrocaudally, the trajectory and orientation of labeled fibers were confirmed by tracing the U-fibers across coronal sections. In both cases, terminations on the postcentral gyrus were located on the corresponding portion of the gyrus directly caudal to the precentral injection and labeled fiber profiles were consistent with a symmetric organization orthogonal to the local long axis of the central sulcus.

In contrast, U-fibers around the IPS exhibited an asymmetric organization. In the three cases with SPL injections (BMD, BMP, BMT), U-fibers first traveled rostrally and ventrally (∼4 mm) within the gyrus before they curved beneath the IPS to terminate on a specific cortical area on the crown of the IPL (Figure 10C-E). The terminal field (Figure 10C, red arrow) was therefore located approximately 4 mm rostral to the injection site (Figure 10E, red asterisk), rather than directly across the sulcus. Accordingly, the bundle traveled at a considerable angle relative to the long axis of the IPS rather than adopting a perpendicular orientation (Figure 10B). Notably, fibers around the fundus of the IPS did not remain in the coronal plane; fiber profiles transitioned from transverse (consistent with a rostrocaudal trajectory out of the plane of section) to longitudinal (consistent with a trajectory within the plane of section) as the bundle curved beneath the fundus and entered the IPL white matter (see Figure 17B). A similar transition was observed at the rostral IPS (BMDH), where profiles that emerged from the injection were predominantly longitudinal before they shifted to transverse beneath the fundus (see Figure 17D). This transition indicates that fibers that crossed the fundus adopted a trajectory parallel to the long axis of the IPS rather than maintaining the orthogonal orientation observed proximal to the injection in the IPL, a pattern inconsistent with symmetrical classification. Labeled fibers also continued caudally beyond the main injection foci, curving beneath the IPS and terminating on the opposing gyrus.

### 2.4 Local Connections Define a Specific Region of the White Matter

The precise delineation of the GM-WM border enabled depth measurements of U-fibers and local association fiber bundles relative to the cortex. This analysis revealed that labeled short association fiber bundles display two main depth patterns relative to the overlying sulcal cortex. In some instances, U-fibers (e.g., principal sulcus; Figure 5) and other local bundles (L- and J-type) occupied the white matter directly beneath cortical layer 6 (Figure 11B, G). In other instances (e.g., mid-IPS; Figure 8), bundles were not directly beneath the GM-WM border and instead maintained a consistent distance from that border as they curved around the sulcus, producing a label-free region or ‘gap’ between the fiber bundle and the GM-WM border (Figure 11E, H).

**Figure 11:**
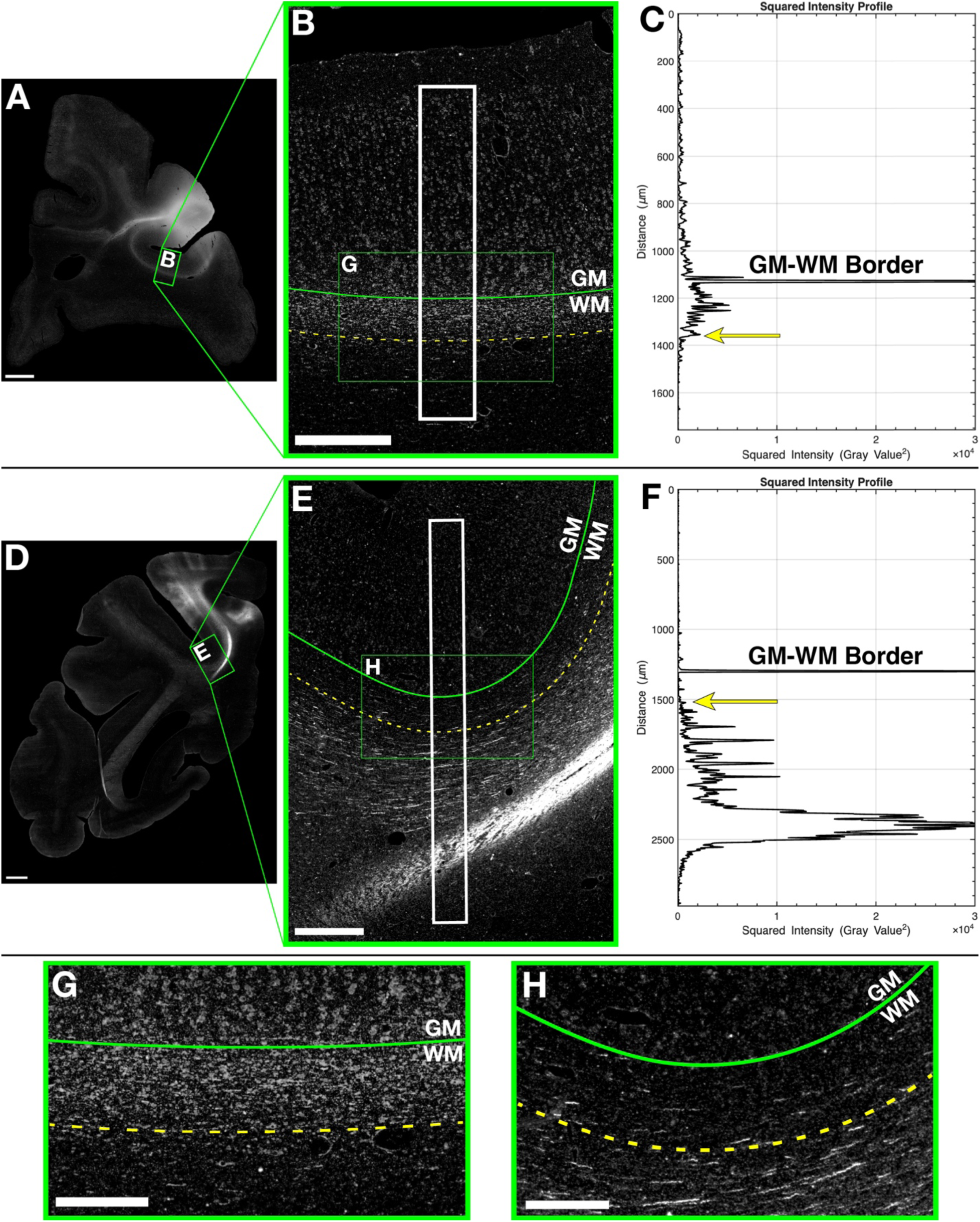
Quantification of bundle depth relative to the gray matter-white matter (GM-WM) border using darkfield intensity profiles. A. Low-magnification darkfield autoradiograph, macaque prefrontal cortex. B. Higher-magnification view of the boxed region in A, showing the GM-WM border (solid green line), rectangular ROI (white box), deep margin of the labeled bundle (dashed yellow line), and area enlarged in G (green box). C. Squared intensity profile from the ROI in B (squared gray value vs. depth) showing the GM-WM border and the deep margin of the labeled band, defined as the deepest local maximum within the high-intensity band (yellow arrow). D. Low-magnification darkfield autoradiograph, macaque parietal lobe. E. Higher-magnification view of the boxed region in D, showing the GM-WM border (solid green line), ROI (white box), traced superficial margin of the deeper labeled bundle (dashed yellow line), and area enlarged in H (green box). F. Squared intensity profile from the ROI in E; here the yellow arrow marks the first peak above the label-free region beneath the GM-WM border. G-H. Enlarged views of the green boxes in B and E, showing the dashed yellow line relative to superficial (G) and deeper (H) fibers. A-C, G, case BMFG; D-F, H, case BMP. Scale bars: A, D = 2 mm; B, E = 500 µm; G-H = 250 µm.

To determine the relationship between these two patterns, fiber bundle depth relative to the GM-WM border was measured using a two-stage approach. For fiber bundles occupying the white matter directly beneath cortical layer 6 (Figure 11A-C, G), the distance from the GM-WM border (Figure 11B, G; solid green line) to the deep margin of the labeled fiber bundle (Figure 11B, G; dashed yellow line) was measured. For fiber bundles positioned in deeper portions of the white matter (Figure 11D-F, H), the distance from the GM-WM border (Figure 11E, H; solid green line) to the superficial margin of the labeled fiber bundle (Figure 11E, H; dashed yellow line) was measured instead. When labeled fiber bundles were located superficially, directly beneath cortical layer 6, the deep margin was located at a mean depth of 256 µm from the GM-WM border (SD 24.5 µm; range 207-297 µm; 27 ROIs across 6 cases; Figure 12). When fiber bundles were present in deeper portions of the white matter, the superficial margin was located at a mean depth of 249 µm from the GM-WM border (SD 36.9 µm; range 173-327 µm; 31 ROIs across 6 cases; Figure 12).

**Figure 12:**
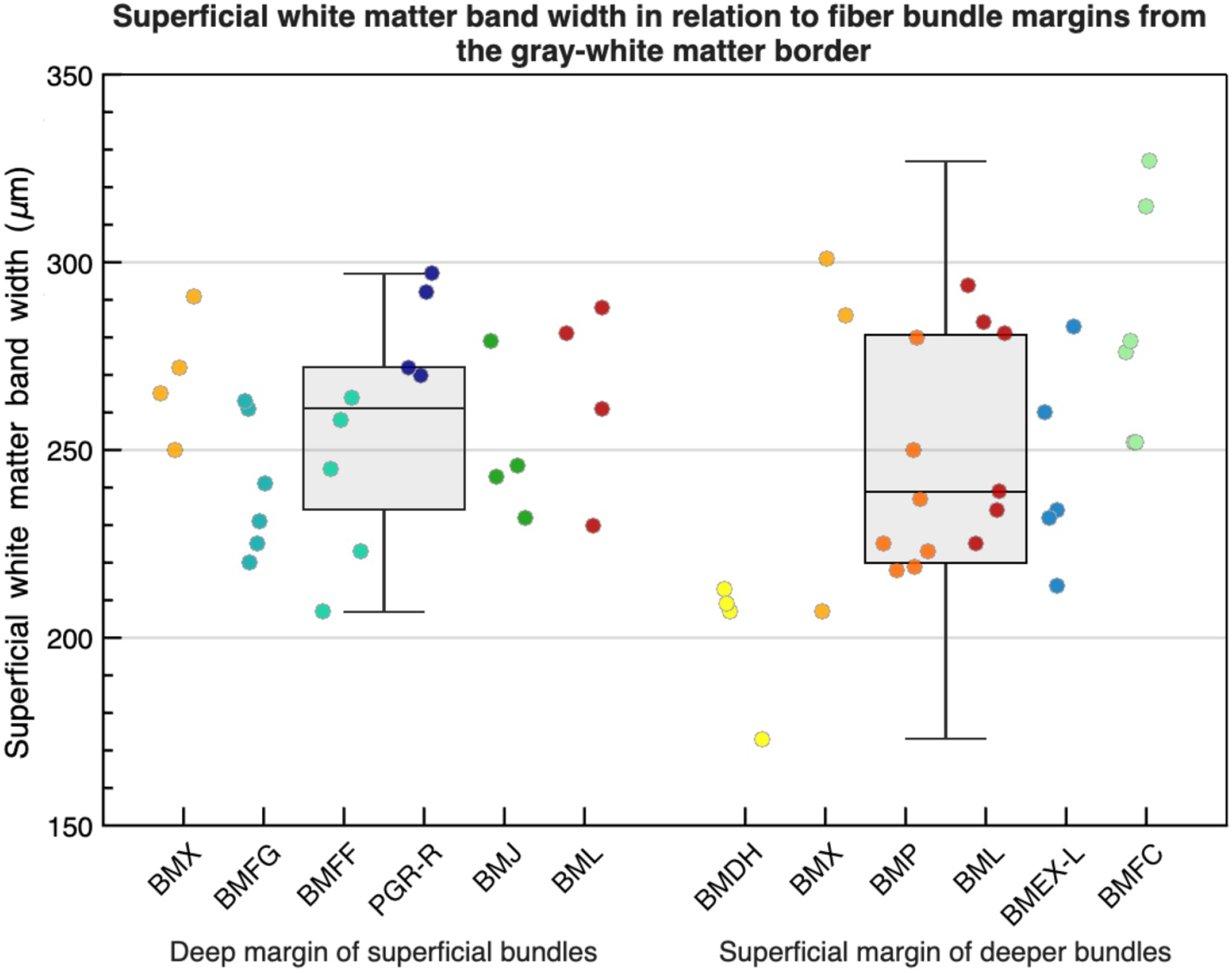
Depth measurements relative to the gray matter-white matter (GM-WM) border for the two labeling patterns. Scatter plot with overlaid box-and-whiskers showing white matter depth (µm) from the GM-WM border (y-axis) by case, grouped by labeling pattern (x-axis): deep margin of superficially labeled bundles (left) and superficial margin of deeply labeled bundles (right). Points are color-coded as in Figure 3; overlaid box-and-whisker plots show median, interquartile range, and full range.

Despite the two distinct labeling patterns, a statistical comparison of these two depth profiles (GM-WM border to deep margin of superficial bundles vs. GM-WM border to superficial margin of deeper bundles) revealed no significant difference (Welch’s t(52.55) = 0.799, p = 0.428). Figure 12 plots individual ROIs by case and labeling pattern with overlaid boxplots. Although measurement of the position of fibers in the deeper compartment showed higher variability, the distributions overlapped, with medians for both labeling patterns falling within ∼235-265 µm (Figure 12). Together, these results indicate that the two fiber bundle labeling patterns define a consistent superficial white matter band extending approximately 200-300 µm beneath the GM-WM border (i.e., immediately below cortical layer 6), with superficial bundles occupying the band and deeper bundles coursing just beneath it. Across all cases, labeled fibers entered this band only to terminate in the overlying cortex; none traveled within it and then re-entered the subjacent white matter. This band, therefore, likely represents a region in which association fibers enter as they approach their cortical targets.

In all regions sampled, the superficial band was characterized as a discrete 200-300 µm structure immediately beneath cortical layer 6 (mean width 252 µm; SD 31.6 µm; range 173-327 µm; 58 ROIs total). See Table 3 for per-sulcus and per-case values. When depth measurements were grouped by sulcus (regardless of labeling pattern), there were no significant differences in the width of the superficial band (one-way ANOVA: F(6, 51) = 1.88, p = 0.103). Boxplots showed broadly overlapping distributions of width across sulci, with medians between ∼240-280 µm (Figure 13).

**Figure 13:**
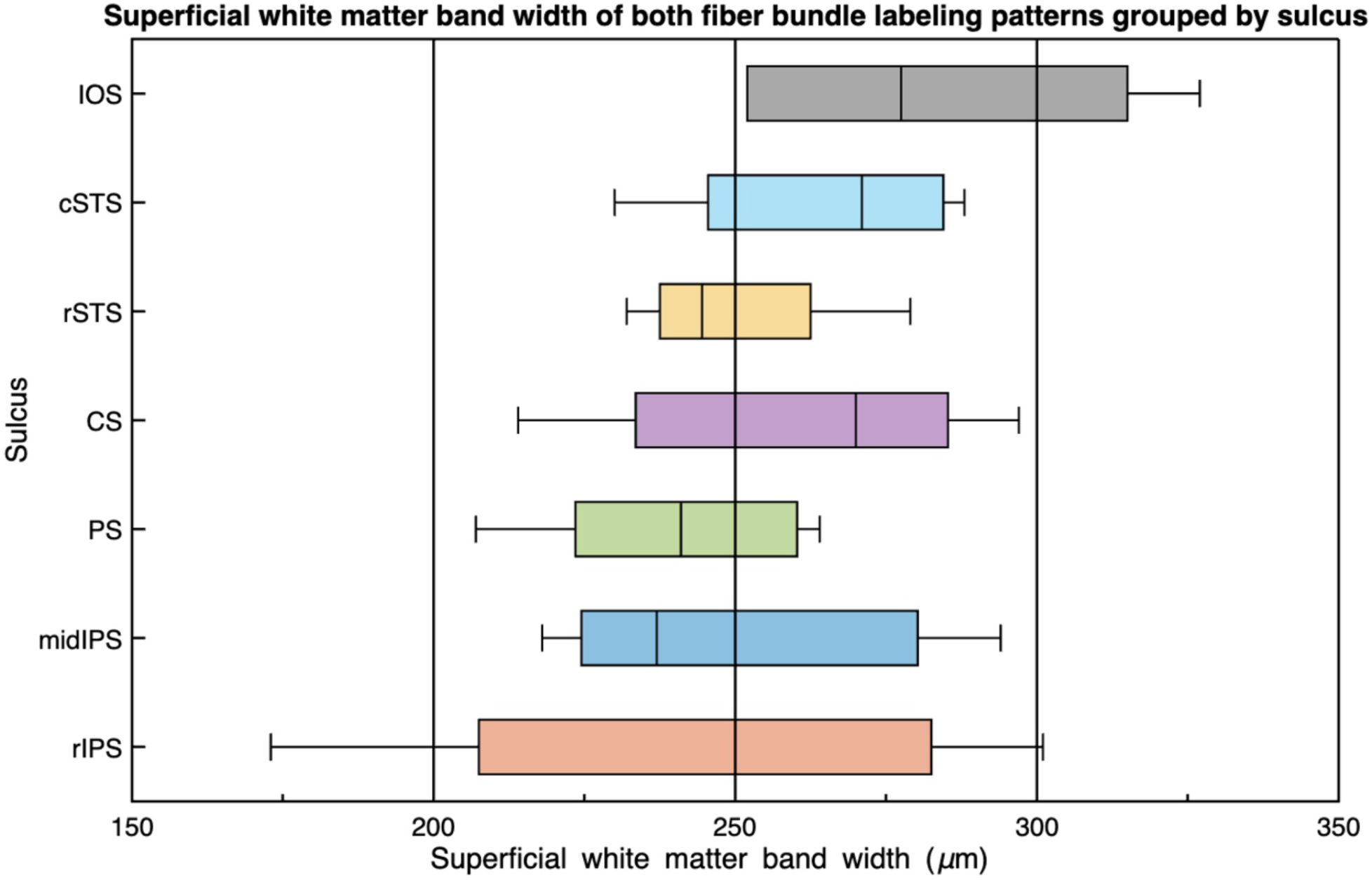
Superficial white matter band width by sulcus. Boxplots of the superficial band width (µm; y-axis) across cases grouped by sulcus (x-axis): rostral intraparietal (rIPS), mid-intraparietal (midIPS), principal (PS), central (CS), rostral superior temporal (rSTS), caudal superior temporal (cSTS), and inferior occipital (IOS). Boxes show median and interquartile range; whiskers show the observed range. No significant difference was observed across sulci (see text).

**TABLE 3.** Width summary of the superficial white matter band by sulcus and case.

| Sulcus | Case | Relationship to GM-WM border<sup>a</sup> | Fiber Bundle Type | Superficial Band Width ($\mu\text{m}$)<sup>b</sup> |
| --- | --- | --- | --- | --- |
| PS | BMFF | No-gap | U-Fiber | 239 ± 23.99 (207-264; n = 5) |
| | BMFG | No-gap | U-Fiber | 240 ± 18.31 (220-263; n = 6) |
| **Total** |  |  |  | **240 ± 19.95 (207-264; n = 11)** |
| CS | BMEX-L | Gap | U-Fiber | 245 ± 27.01 (214-283; n = 5) |
| | PGR-R | No-gap | U-Fiber | 283 ± 13.74 (270-297; n = 4) |
| **Total** |  |  |  | **262 ± 28.98 (215-297; n = 9)** |
| rSTS | BMJ | No-gap | U-Fiber | **250 ± 20.25 (232-279; n = 4)** |
| cSTS | BML | No-gap | U-Fiber | **265 ± 25.99 (230-288; n = 4)** |
| IOS | BMFC | Gap | U-Fiber | **284 ± 31.45 (252-327; n = 6)** |
| rIPS | BMX | No-gap | L-Type | 270 ± 17.02 (250-291; n = 4) |
| | | Gap | J-Type | 265 ± 50.5 (207-301; n = 3) |
| | BMDH | Gap | U-Fiber | 201 ± 18.5 (173-213; n = 4) |
| **Total** |  |  |  | **243 ± 42.94 (173-301; n = 11)** |
| midIPS | BMP | Gap | U-Fiber | 236 ± 22.49 (218-280; n = 7) |
| | BML | Gap | Complex U-Shaped | 260 ± 30.04 (225-294; n = 6) |
| **Total** |  |  |  | **247 ± 27.89 (218-294; n = 13)** |

<sup>a</sup> Gap = distance from gray matter-white matter (GM-WM) border to the superficial margin of a deeper labeled fiber bundle (i.e., a layer devoid of label); No-gap = distance from GM-WM border to the deep margin of a superficially labeled fiber bundle.
<sup>b</sup> Values pooled across ROIs within each case/sulcus.
Bold values indicate sulcus-level totals (pooled across cases within each sulcus). Abbreviations: PS, principal sulcus; CS, central sulcus; rSTS, rostral superior temporal sulcus; cSTS, caudal superior temporal sulcus; IOS, inferior occipital sulcus; rIPS, rostral intraparietal sulcus; midIPS, mid intraparietal sulcus.

### 2.5 U-fibers Occupy Two Depth Compartments

Having established the existence of a consistent superficial white matter band directly beneath the GM-WM border, we next asked whether U-fibers coursed within this band or avoided it throughout their full trajectory around the sulcus. The position of U-Fibers relative to this band was evaluated in the 10 cases in which they were present beneath the targeted major sulcus, with depth measurements obtained in 8 of the cases (Table 3) and confirmed qualitatively in the remaining 2 (BMD, BMT), which had labeled fibers at a similar portion of the mid-IPS as case BMP. Surprisingly, only 50% (5/10) of U-fibers occupied the superficial band throughout their trajectory, while the remaining 50% (5/10) avoided this band around the sulcus (see Table 1). U-fibers could therefore be divided into two distinct groups based on depth; this depth-based division also revealed a strong correspondence between depth and symmetry.

#### 2.5.1 U-fibers in the more Superficial White Matter Compartment

Superficial U-fibers (or ‘true’ U-fibers) were defined as possessing all of the following three characteristics: 1) labeled fibers adopted a U-shaped course beneath a sulcus to link adjacent gyri; 2) the fiber bundle was symmetric, terminating on the corresponding portion of the opposing gyrus while maintaining a consistent trajectory approximately orthogonal to the local long axis of the sulcus (i.e., Meynert’s “wire-rings”) and 3) the bundle occupied the ∼200-300 µm-thick superficial band throughout its trajectory.

The U-fibers beneath the principal sulcus in both prefrontal cases (BMFF, BMFG; Figure 14) met all three criteria. Each bundle (Figure 14A, C; solid red line) adopted a U-shaped trajectory, emanating from the injection site on one gyrus (Figure 14A, C; red asterisks) and coursing around the sulcus to the opposing gyrus. Columnar terminations (Figure 14A, C; red arrows) were located on the corresponding portion of the opposing gyral crown at approximately the same rostrocaudal level as the injection. In both bundles, fiber profiles maintained relatively consistent orientations within the plane of section throughout their trajectories (Figure 14B, D), consistent with an orientation perpendicular to the local long axis of the sulcus. Depth measurements confirmed that each U-fiber coursed within the ∼200-300 µm superficial band (Figure 14A-D, dashed yellow line) directly beneath the GM-WM border (BMFG: 240 ± SD 18.3 µm; range 220-263 µm (Figure 11A-C, G); BMFF: 239 ± SD 24.0 µm; range 207-264 µm). These values fell within the range established earlier for the superficial white matter band and were comparable to measurements in other sulci.

**Figure 14:**
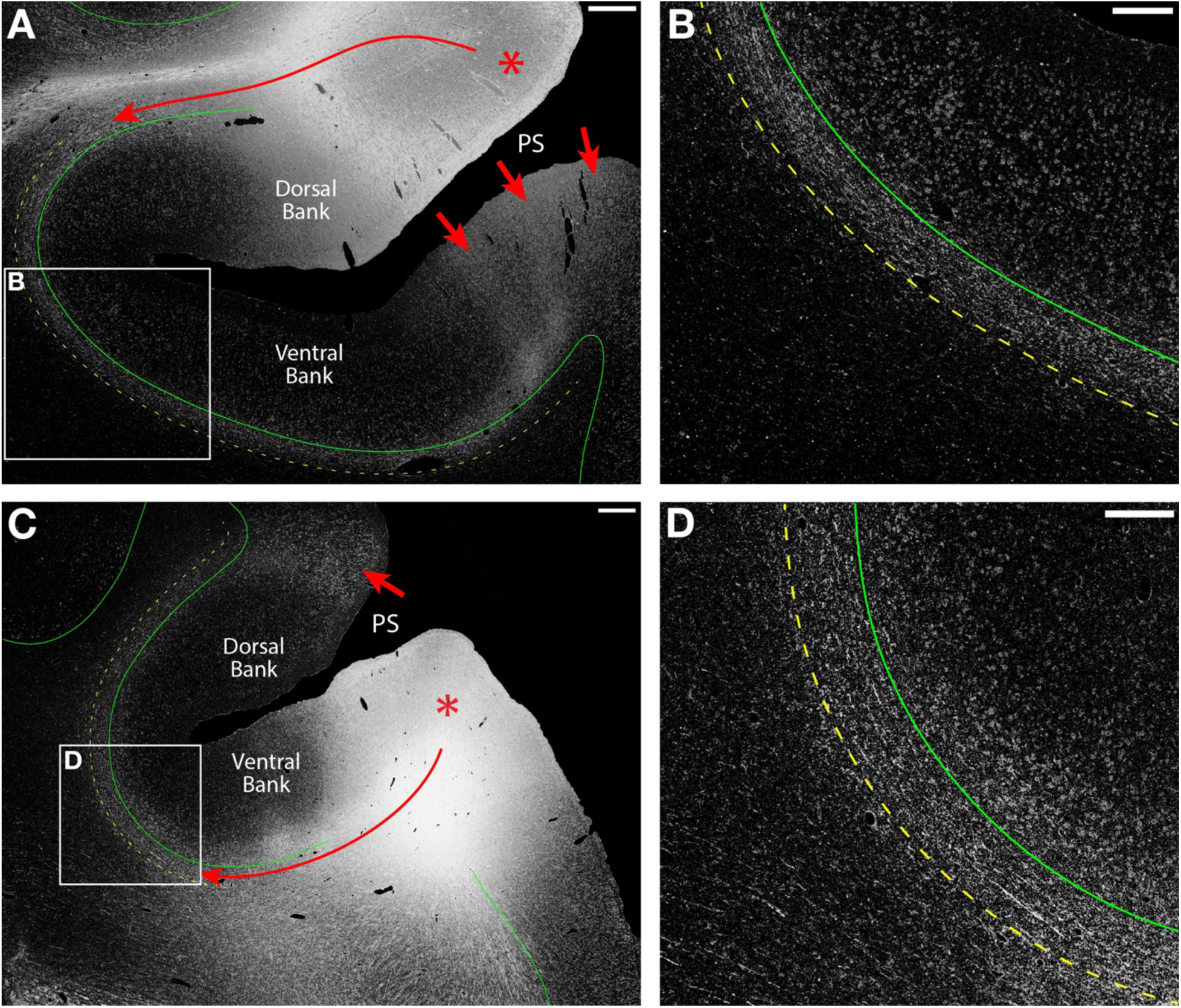
U-fibers confined to the superficial white matter band in the prefrontal cortex. Cases are from Figure 5 and are shown to highlight U-fibers restricted to the superficial band (dashed yellow line) immediately beneath the gray matter-white matter (GM-WM) border (solid green line). A, C. Higher-magnification darkfield images: U-fibers (solid red lines) course from the injections (red asterisks) around the principal sulcus (PS) to terminate on the adjacent gyral crowns (red arrows), remaining within the superficial bands throughout. B, D. Enlarged views of the boxed regions in A and C, showing the U-fibers coursing parallel to the GM-WM borders within the superficial bands. A-B, case BMFG; C-D, case BMFF. Scale bars: A, C = 500 µm; B, D = 250 µm.

Along the central sulcus, the U-fiber in case PGR-R had a similar organization and met all three criteria. Labeled fibers adopted a U-shaped trajectory (Figure 15, dashed red line), coursing caudally beneath the central sulcus to interconnect the precentral gyrus with the corresponding portion of the opposing postcentral gyrus, with additional terminations on the opposing sulcal bank (red arrows). Throughout the fiber bundle’s trajectory, labeled fiber profiles were consistent with a symmetric, caudally directed U-fiber positioned directly beneath the GM-WM border. At the sulcal fundus, the bundle’s deep margin was located at a mean depth of 283 µm from the GM-WM border (SD 13.7 µm; range 270-297 µm), confirming that the fibers fully occupied the superficial band (Figure 15, dashed yellow line) at that position. After passing around the fundus, the compact fibers fanned out and turned superiorly toward the postcentral gyrus but remained within the superficial band.

**Figure 15:**
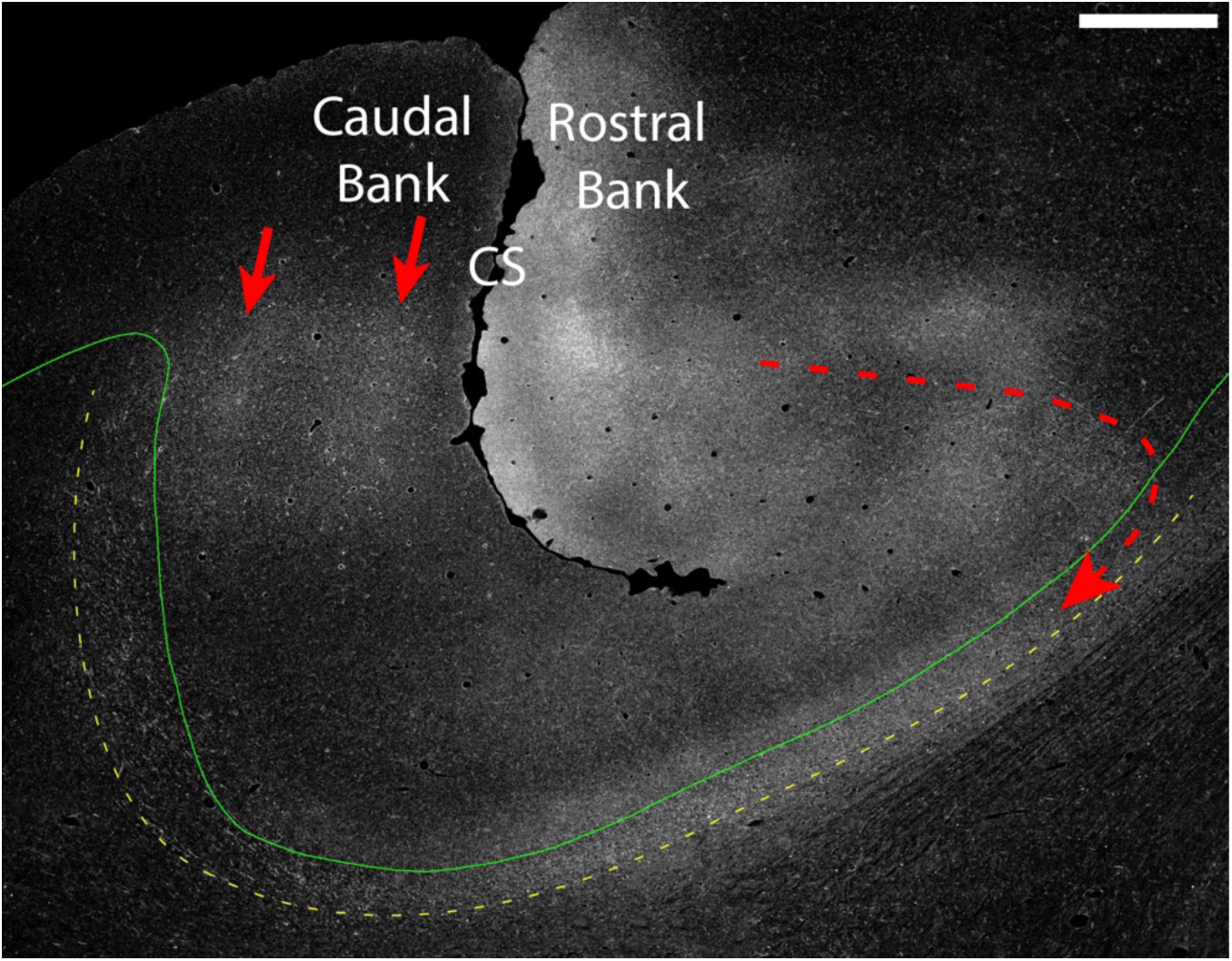
U-fiber confined to the superficial white matter band beneath the central sulcus. Higher-magnification darkfield image (case PGR-R, from Figure 6A-B): a U-fiber (dashed red line) courses beneath the central sulcus (CS), with terminations along the caudal bank (red arrows) and further caudally on the postcentral crown. The bundle remains within the superficial band (dashed yellow line) directly beneath the gray matter-white matter border (solid green line); the bundle is compact at the fundus and fans out within the postcentral white matter. The oblique plane of section makes the superficial band appear expanded along the caudal bank. Scale bar: 1 mm.

Only 50% (5/10) of U-fibers observed beneath the major sulci examined satisfied all three criteria and thus qualified as superficial (‘true’) U-fibers. These were not evenly distributed across the cortex; they accounted for 100% (2/2) of U-fibers observed beneath the principal sulcus (BMFG, BMFF), 100% (2/2) beneath the STS (BMJ, BML), and 50% (1/2) beneath the central sulcus (PGR-R) (Table 1). Across all U-fibers, symmetry was associated with a superficial position: the majority of symmetric bundles (83.3%; 5/6) coursed within the superficial band (Table 1). Meynert’s conception of symmetric U-fibers arranged along the length of a sulcus and occupying the white matter directly beneath the cortex (Figure 16A) therefore, captures only a subset of U-fibers.

**Figure 16:**
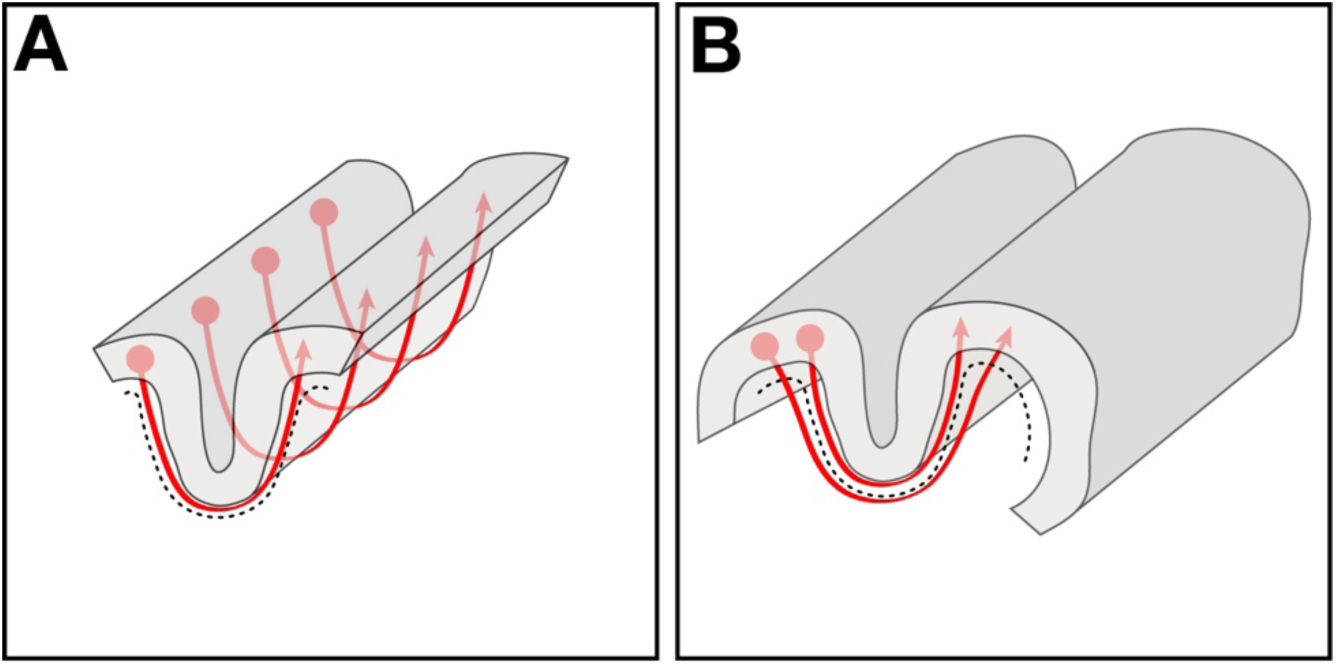
Schematics contrasting Meynert’s (1885) classic U-fiber principles. A. Classic symmetric U-fibers stacked along the length of a sulcus, interconnecting corresponding portions of adjacent gyri with trajectories orthogonal to the sulcal long axis and confined to the superficial white matter (dashed black line). B. Meynert’s rule: longer U-fibers interconnect more distant regions and course deeper in the white matter.

#### 2.5.2 U-fibers in Deeper Portions of the White Matter

The remaining 50% (5/10) of U-fibers beneath the major sulci examined traveled in deeper portions of the white matter. These bundles adopted U-shaped trajectories beneath sulci and interconnected adjacent gyri but coursed below the superficial white matter band. Consistent with Meynert’s description, these deeper U-fibers generally interconnected more distant cortical regions. However, their organization was not as simple as a longer symmetric U-fiber positioned beneath a shorter one (e.g., Figure 16B). Most deeper U-fibers originated some distance from where they crossed beneath the sulcus. Deeper U-fibers were most often observed along the mid-IPS, accounting for 60% (3/5) of all deeper U-fibers (Table 1).

In the three mid-IPS cases with SPL injections (BMD, BMP, BMT), labeled fibers first traveled rostrally and ventrally for approximately 4 mm within the SPL before wrapping around the IPS (Figure 17A, dashed red line) to terminate in a specific cortical area on the crown of the IPL (red arrow; alos see Figure 10C-E). Across their trajectories, the labeled fiber bundles followed the contour of the IPS while consistently avoiding the superficial band (Figure 17A-B, dashed yellow line) directly beneath the GM-WM border. In case BMP, the superficial margin of the fiber bundle maintained a consistent mean depth of 236 µm from the GM-WM border (SD 22.5 µm; range 218-280 µm) (see Figures 11D-F, H; 17A-B). At the rostral IPS, labeled fibers in case BMDH similarly avoided the superficial band (Figure 17C-D, dashed yellow line); the bundle’s superficial margin was at a mean depth of 201 µm (SD 18.5 µm; range 173-213 µm). In contrast to the mid-IPS cases, labeled fibers in case BMDH crossed the IPS at the level of the injection site rather than first traveling rostrally. The bundle then continued as a U-shaped bundle beneath the IPS, extending approximately 4 mm caudally beyond the main injection focus to terminate on the opposing gyrus.

**Figure 17:**
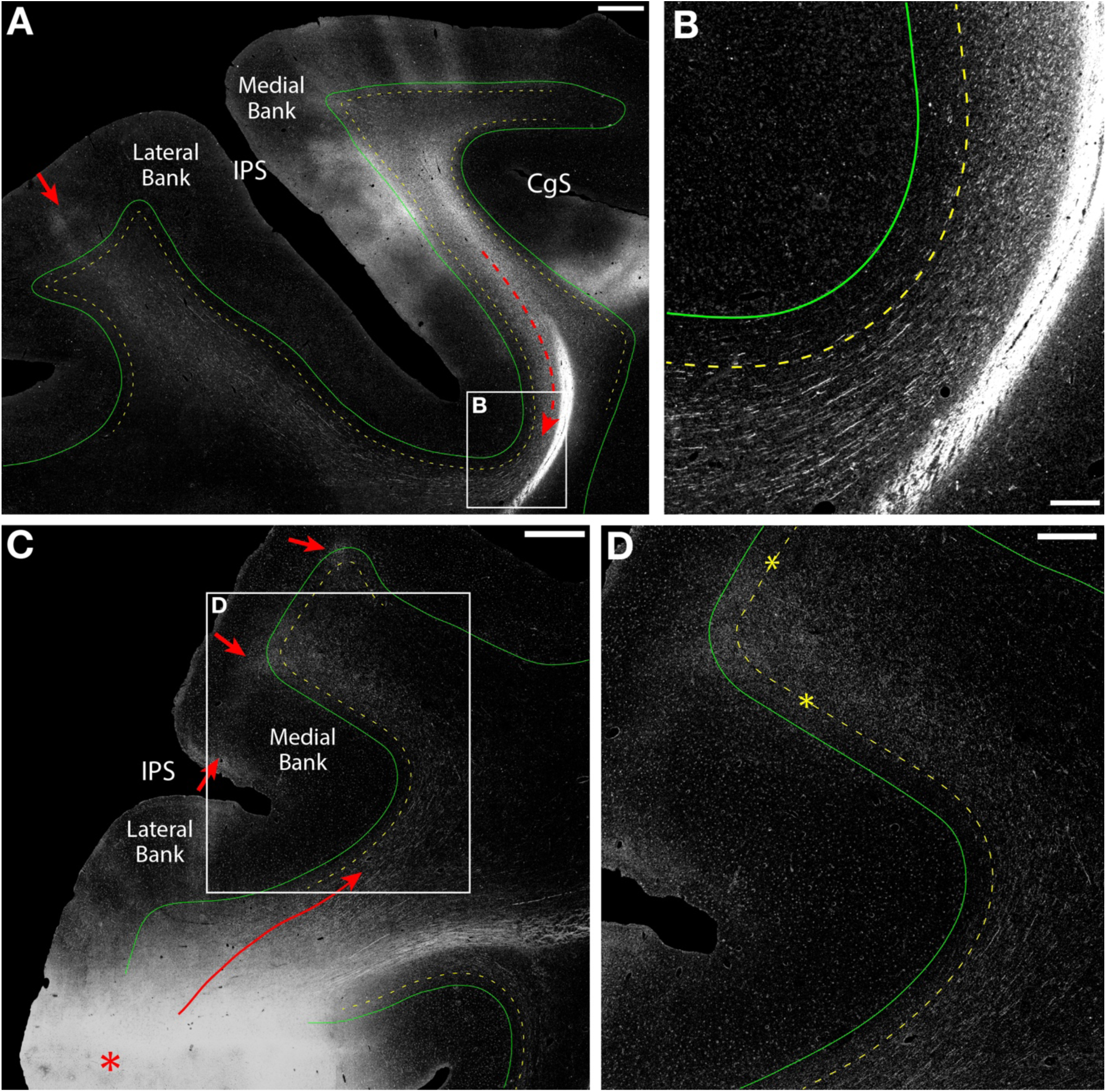
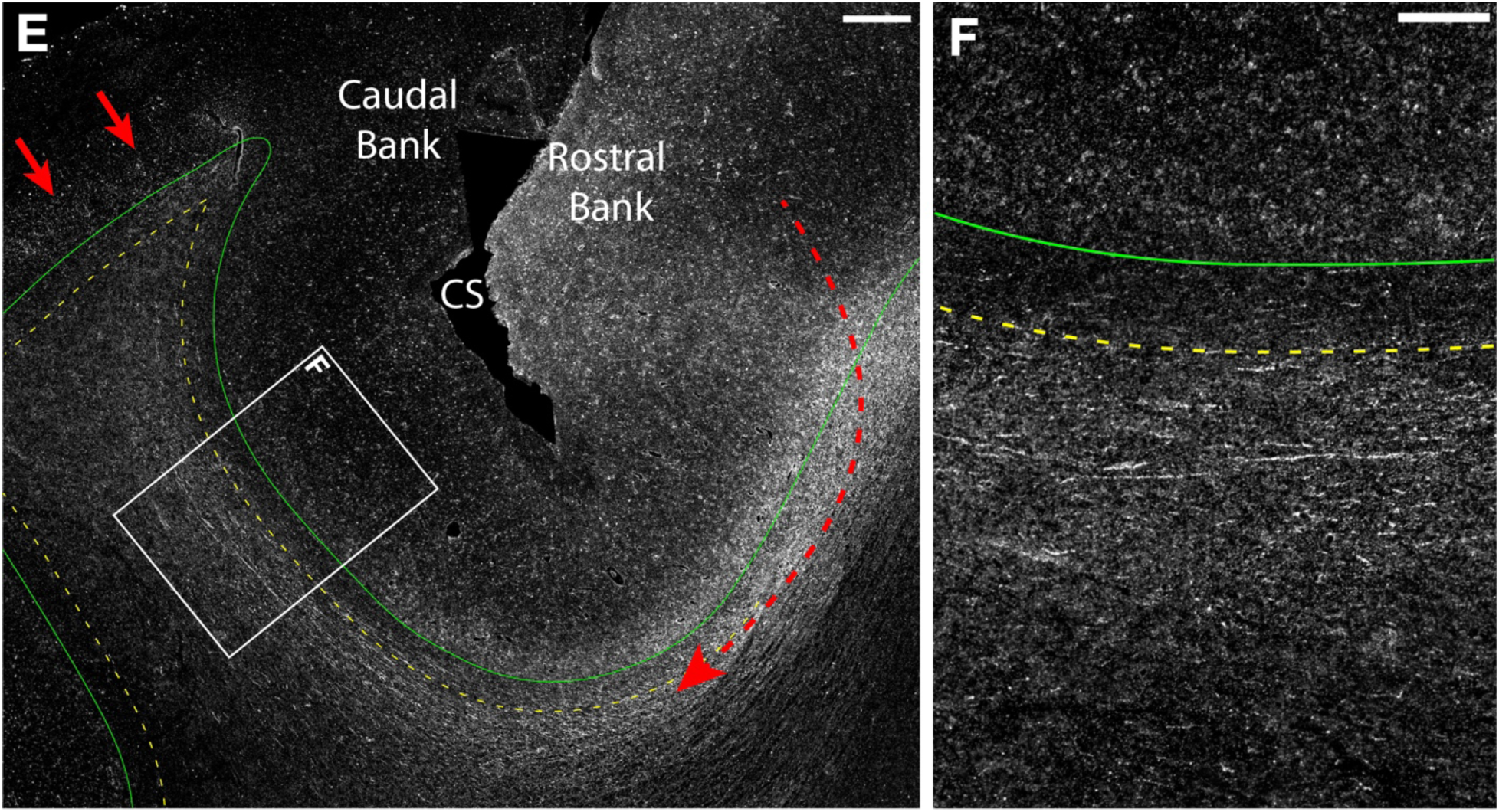
Deeper U-fibers that avoid the superficial white matter band. Coronal sections from three cases (from Figures 6, 7 and 8), imaged under darkfield. A. Low-magnification image, mid portion of the intraparietal sulcus (IPS): an asymmetric U-fiber (dashed red line) courses rostrally from a superior parietal lobule (SPL) injection, curves beneath the IPS, and terminates on the inferior parietal lobule (IPL) crest (red arrow), traveling in deeper portions of the white matter and avoiding the superficial band (dashed yellow line) beneath the gray matter-white matter (GM-WM) border (solid green line). B. Enlarged view of the boxed region in A: labeled fiber profiles course deep to the superficial band and leave it label-free. C. Low-magnification image, rostral IPS: an asymmetric U-fiber (solid red line), from an IPL injection (red asterisk), courses beneath the IPS in deeper portions of the white matter, providing terminations on the medial IPS bank and SPL crown (red arrows). D. Enlarged view of the boxed region in C: labeled fibers travel beneath the superficial band and then cross it to enter the cortex (dashed yellow line and asterisks). E. Low-magnification image, frontal/parietal lobes; a symmetric U-fiber (dashed red line) courses caudally from a precentral injection beneath the central sulcus (CS) to terminate on the postcentral gyrus (red arrows), traveling in deeper portions of the white matter and leaving the superficial white matter band free of label. F. Enlarged view of the boxed region in E: labeled fibers deep to the superficial band (dashed yellow line). A-B, case BMP; C-D, case BMDH; E-F, case BMEX-L. Scale bars: A, C = 1 mm; B, F = 250 µm; D-E = 500 µm. CgS, cingulate sulcus.

In the motor cortex (case BMEX-L), the remaining deeper U-fiber was located beneath the central sulcus. The injection in this case was positioned on the precentral gyrus farther from the sulcus than in the other motor cortex case (see Figure 3). Similar to the deeper U-fibers beneath the IPS, labeled fibers in BMEX-L (Figure 17E-F) consistently avoided the superficial band, with the bundle’s superficial margin at a mean depth of 245 µm from the GM-WM border (SD 27.0 µm; range 214-283 µm). Despite its deeper position, this U-fiber was classified as symmetric, as labeled fibers coursed caudally beneath the sulcus to interconnect the precentral gyrus with the corresponding portion of the opposing postcentral gyrus, similar to the superficial U-fiber in the other central sulcus case (PGR-R). The greater depth corresponded to the greater distance of the injection and terminal field from the sulcus, matching Meynert’s original observation that longer U-fibers travel deeper in the white matter (Figure 16B).

Similar to the superficial U-fibers, deeper U-fibers were not evenly distributed across the cortex, accounting for 100% (4/4) of bundles beneath the IPS (BMD, BMP, BMT, BMDH) and 50% (1/2) beneath the central sulcus (BMEX-L) (Table 1). In all five cases, deeper U-fibers were considerably longer than superficial U-fibers, indicating a relationship between U-fiber depth and bundle length. However, apart from the central sulcus, there was no evidence for the concentric ring-type organization described by Meynert, in which shorter superficial U-fibers are layered with progressively longer, deeper bundles beneath the same sulcus (e.g., Figure 16B).

In addition, all asymmetric U-fibers (4/4; 100%) were deeper U-fibers that avoided the superficial band (Table 1). In the majority of these asymmetric cases, depth did not correspond consistently to how far the interconnected gyral regions were from the sulcus. At the rostral IPS (BMDH), the injection was located farther from the sulcus but labeled fibers terminated on the opposing gyrus close to the sulcus (see Figures 3 and 17C), while at the mid-IPS (BMP, BMT), injections were close to the sulcus but labeled fibers terminated on the opposing gyrus farther from it (see Figures 3; 10C-E). The remaining mid-IPS case (BMD) followed the expected pattern, with a more medial injection producing terminations farther from the sulcus on the adjacent gyrus (see Figure 3). Together, these observations indicate that deep U-fibers do not consistently follow Meynert’s rule that deeper bundles interconnect more distant cortical regions (e.g., Figure 16B).

### 2.6 Other Deeper Fiber Bundles That Appeared U-Shaped Often Exhibited More Complex Shapes and Trajectories

During the analysis of all 28 cases, several instances were identified in which large, labeled fiber bundles occupied the depth of a sulcus in a U-shaped configuration. These bundles consistently occupied a deeper position within the white matter, similar to the deeper U-fibers described above. However, their trajectories, origins, and terminations indicated that these bundles did not interconnect adjacent gyral crowns and thus were not classified as U-fibers. Therefore, a U-shaped course beneath a sulcus does not by itself identify a U-fiber. These deeper fiber bundles with more complex shapes were identified in 17.9% (5/28) of cases analyzed (Table 1). Based on their origins and terminations, two reciprocal types of these fiber bundles were observed (Figure 18).

**Figure 18:**
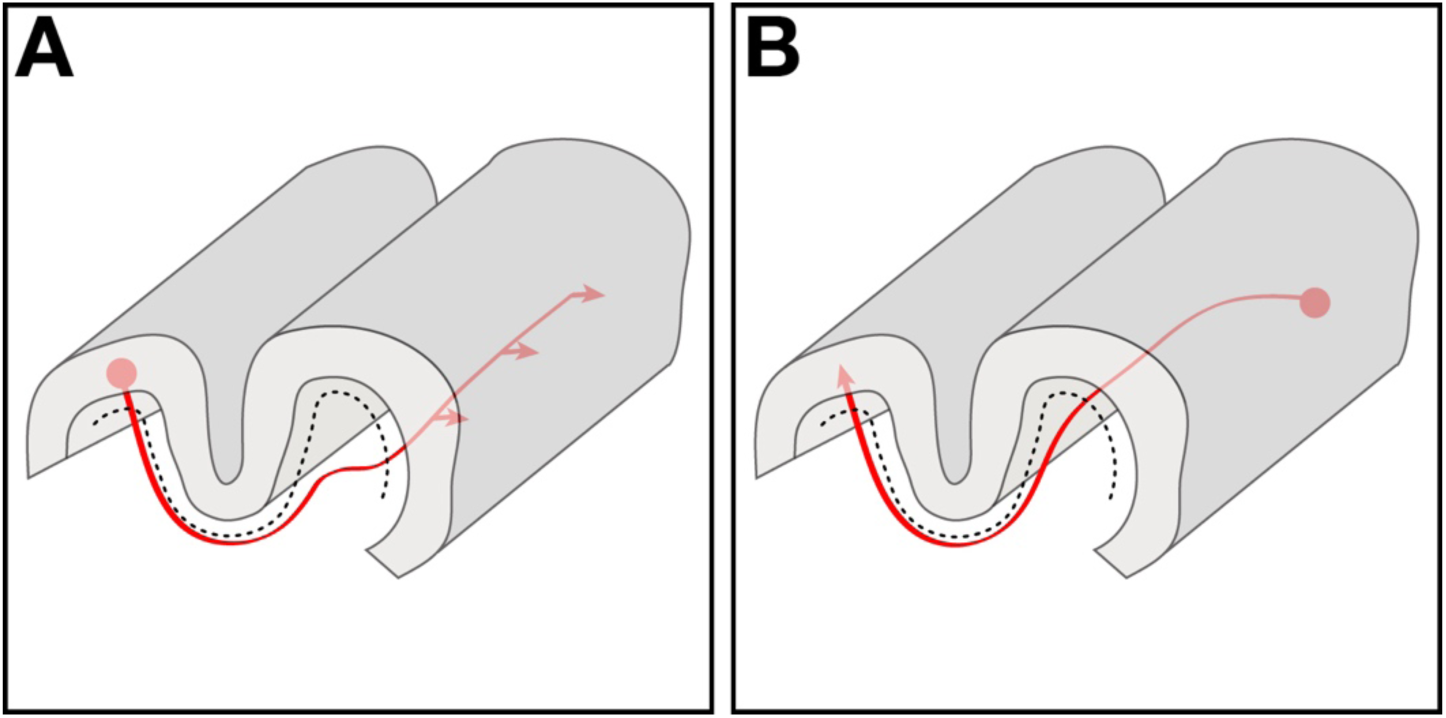
Schematics of two reciprocal deeper bundle types that appear U-shaped beneath a sulcus but do not interconnect adjacent gyral crowns and therefore are not classified as U-fibers. Both bundle types avoid the superficial band (dashed black line) and follow more complex trajectories than the deeper U-fibers described here. A. A bundle from a gyral-crown injection hooks beneath the fundus and ascends the opposing bank and turns medially through the gyral white matter to terminate on the medial surface of the hemisphere. B. The reciprocal case: a bundle from a medial hemisphere injection crosses the gyral white matter, then adopts a U-shaped trajectory beneath the fundus to reach the opposing gyral crown.

In the first instance (Figure 18A), a fiber bundle originating from a gyral crown injection, hooks beneath the sulcal fundus with a U-shaped trajectory, similar to a U-fiber, but rather than progressing to the adjacent gyrus, it turns medially to terminate on the medial surface of the hemisphere.

This is illustrated in case BML at the mid-IPS (Figure 19), in which an injection on the IPL crown (red asterisk) produced labeled fibers that descended through the core of the gyral white matter and hooked beneath the sulcal fundus. Labeled fibers traveled briefly along the opposing sulcal bank, then turned medially, coursed obliquely across the white matter core of the gyrus, and terminated on the medial hemisphere (i.e., the medial parietal lobe) (Figure 19, red arrows), with a terminal field spanning approximately 10 mm rostrocaudally (see Supplementary Case Atlas). Throughout the bundle’s course along the IPS, labeled fibers avoided the superficial band (Figure 19, dashed yellow line); the bundle’s superficial margin was at a mean depth of 260 µm from the GM-WM border (SD 30.0 µm; range 225-294 µm).

**Figure 19:**
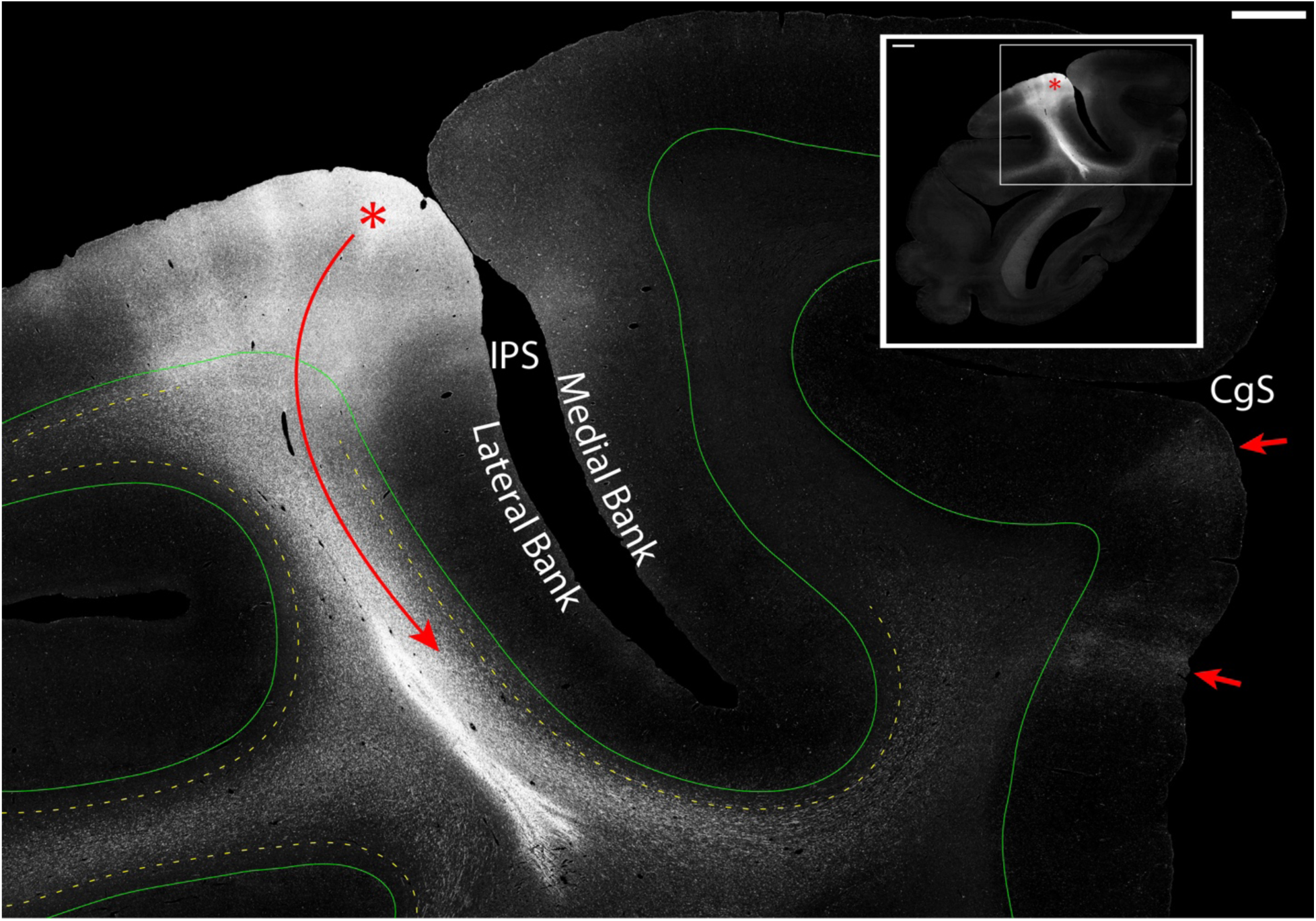
Deeper fiber bundles that appear U-shaped often follow complex trajectories. Coronal section, macaque parietal lobe, imaged under darkfield. Main panel: higher-magnification view of the inset region showing a U-shaped trajectory (solid red line); labeled fibers from the injection (red asterisk) course along the medial intraparietal sulcus (IPS) bank, hook beneath the fundus, and briefly follow the opposing bank like a U-fiber, but then turn medially through the gyral core to terminate on the medial hemispheric cortex (red arrows) ventral to the cingulate sulcus (CgS). The bundle travels in deeper portions of the white matter, deep to the superficial band (dashed yellow line) beneath the gray matter-white matter border (solid green line). Inset: low-magnification image following an injection in the inferior parietal lobule (red asterisk). Case BML. Scale bars: main panel = 1 mm; inset = 2 mm.

The second instance (Figure 18B) shows a mirror-image trajectory: fibers from an injection on the medial surface of the hemisphere crossed the gyral white matter, hooked beneath the sulcal fundus, and terminated on the opposite gyral crown. This is illustrated in two preoccipital cases (BMEQ, BMFA) in which injections on the medial parietal cortex (Figure 20B, E; red asterisk) produced labeling that crossed the white matter core of the SPL, curved beneath the IPS, and then followed the opposing sulcal bank to terminate on the gyrus (Figure 20C, F; red arrows) approximately 10 mm rostral to the injection (see Supplementary Case Atlas). Although bundles in both instances adopted a U-shaped trajectory beneath the IPS, they did not interconnect adjacent gyri and were therefore not classified as U-fibers. These deeper fiber bundles with complex U-shaped trajectories existed alongside shorter and more superficial L- and J-type bundles in the superficial white matter band (see Supplementary Case Atlas), consistent with Meynert’s rule.

**Figure 20:**
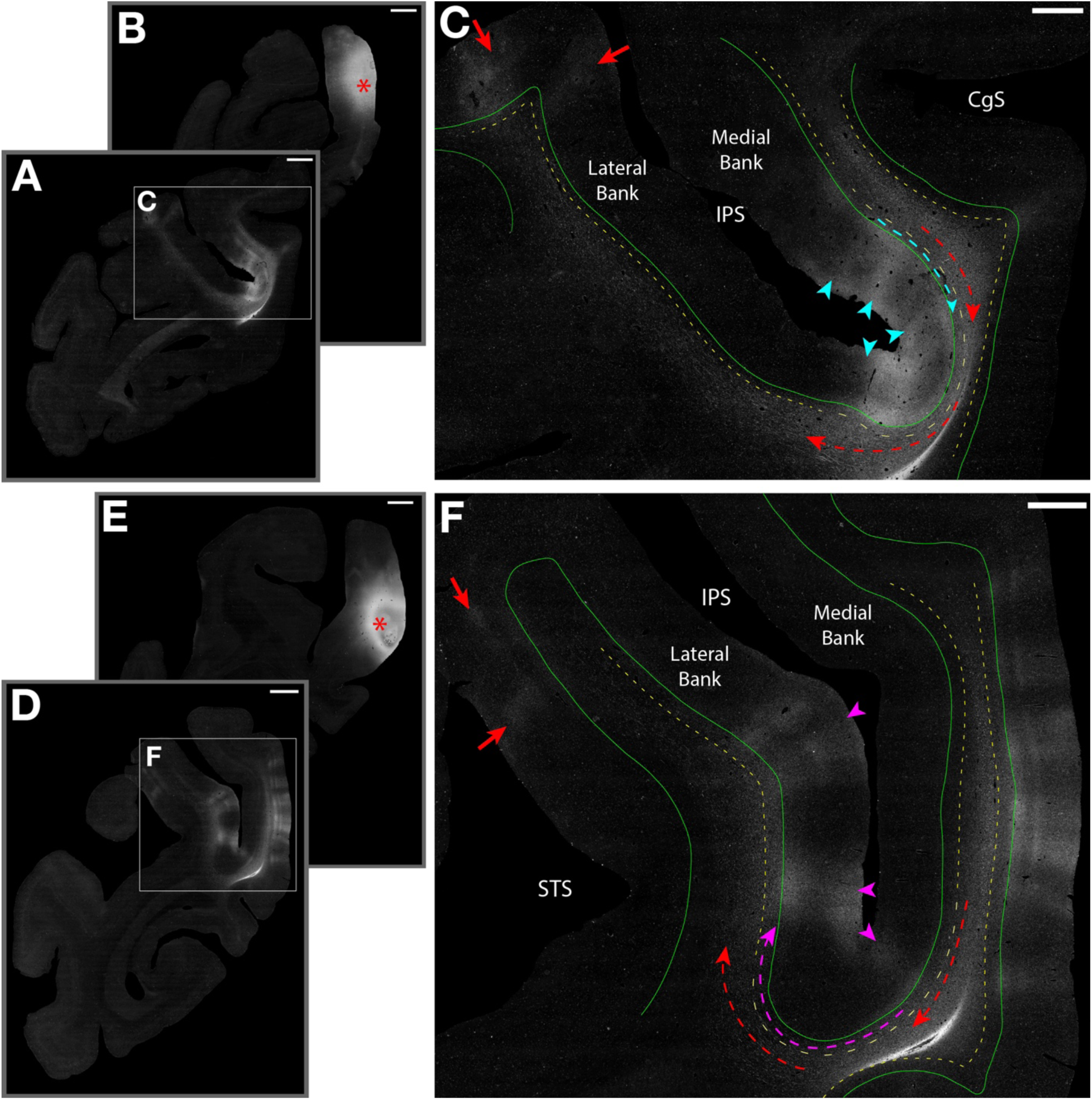
Deeper fiber bundles with U-shapes that do not interconnect adjacent gyral crowns. Darkfield autoradiographs from two preoccipital cases. A-B. Low-magnification images, rostral (A) to caudal (B), ∼10 mm apart; injection on the medial superior parietal lobule (SPL) (B, red asterisk). C. Higher-magnification view of A, showing a bundle with a U-shape (dashed red line) that courses rostrally within the SPL, curves around the intraparietal sulcus (IPS), and forms columnar terminations on the opposing gyral crown and lateral IPS bank (red arrows), remaining in deeper portions of the white matter and avoiding the superficial band (dashed yellow line), and an L-type bundle (dashed teal line) that courses within the superficial band beneath the gray matter-white matter (GM-WM) border (solid green line), terminating along the medial IPS bank and fundus (teal arrowheads). D-E. Low-magnification images, rostral (D) to caudal (E), ∼10 mm apart; injection on the medial SPL (E, red asterisk). F. Higher-magnification view of D: a U-shaped bundle (dashed red line) follows a similar course, curving around the IPS to terminate on the medial superior temporal sulcus (STS) bank (red arrows) and further rostrally on the IPL crown, while avoiding the superficial band; a J-type bundle (dashed magenta line) curves around the IPS fundus but travels within the superficial band, terminating near the fundus and on the lateral IPS bank (magenta arrowheads). A-C, case BMEQ; D-F, case BMFA (reflected for comparison). Superficial-band lines were manually delineated from labeling patterns. Scale bars: A-B, D-E = 2 mm; C, F = 500 µm. CgS, cingulate sulcus.

## 3. Discussion

### 3.1 Summary of findings

In this study, we examined the connectional neuroanatomy of short association fibers and U-fibers in 28 tract-tracing cases from the frontal, temporal, and parietal lobes and preoccipital regions of the rhesus macaque. Four major findings challenge the classic views of U-fibers and the SWM. **First**, although short association fiber bundles and/or local intragyral fibers were identified in every case, U-fibers, bundles that link opposing gyral crowns across a sulcus, were not ubiquitous. In our cases with injections placed on gyri adjacent to major sulci, labeled U-fibers were observed in 50% of cases (35.71% when restricted to the major sulci examined), and their frequency varied according to region, being more common in the prefrontal and motor cortex than in the parietal or temporal lobes. **Second**, U-fibers were not uniformly symmetrical; 60% linked corresponding portions of adjacent gyri with trajectories orthogonal to the long axis of the sulcus, whereas 40% connected opposing gyri at different levels with oblique trajectories. The distribution of symmetry was also sulcus-dependent; U-fibers beneath the central, principal, and superior temporal sulci were largely symmetric, whereas all IPS U-fibers were asymmetric. **Third**, we identified a discrete 200-300 µm superficial white matter band directly beneath cortical layer 6 that was present along every sulcus examined. This region was defined connectionally; short association fiber bundles traveled within it on their approach to their cortical terminations, while longer and deeper bundles consistently avoided it until they terminated. Importantly, U-fibers were found within this band and deeper to it. **Fourth,** U-shaped trajectories were not exclusive to gyrus-to-gyrus interconnections: 17.9% of cases contained deeper, complex U-shaped bundles whose trajectory adopted a U-shape but did not interconnect adjacent gyri. We also identified shorter L-type and J-type bundles that were more frequent at deeper sulci and typically were more superficial than U-fibers at these sulci. To our knowledge, this is the first study to systematically examine SWM organization by tracing the connectivity of the fiber bundles within and around it. Together, these findings reveal a more heterogeneous and regionally specific architecture than classic anatomical models and modern imaging studies have assumed, indicating the presence of U-fibers reflects the specific connectivity of cortical areas rather than the geometry of the sulcus per se, and that different cortical regions may have different white matter organizations.

### 3.2 Historical Models of Superficial White Matter and U-Fiber Organization

U-shaped fiber bundles interconnecting adjacent gyri were depicted and described in early anatomical studies (e.g., Luys, 1865; Arnold, 1883), with the most detailed initial observations made by Meynert (1885). Meynert described U-fibers at the depth of sulci as arranged in a symmetrical pattern, stacked along the length of the sulcus, an arrangement that he likened to a series of “wire rings” (Meynert,1885, p.38). He further proposed that fiber bundle length increased with distance from the overlying cortex (i.e., Meynert’s rule; Catani, 2025), such that U-fibers interconnecting opposing gyral regions close to the sulcus were located more superficially within the white matter. He viewed progressively deeper U-fibers as interconnecting increasingly distant gyral regions relative to the sulcus and possibly traveling beneath multiple sulci. Dejerine and Dejerine-Klumpke (1895) supported and expanded this framework. Building on earlier studies, including those by Sachs (1892), the Dejerines also suggested that U-fibers associated with different sulci should be referred to by distinct names, implying regional differences in their organization and connectivity, a notion consistent with the sulcal specificity observed in the present study. Later gross anatomical dissection studies supported these findings in humans (e.g., Ludwig & Klingler, 1956; Shinohara et al., 2020; Shah et al., 2025) and demonstrated analogous findings in macaque (e.g., Decramer et al., 2018) and capuchin monkeys (e.g., Borges et al., 2015).

The emergence of modern experimental tract tracing in the 1970s did little to challenge or revise the prevailing conception of U-fibers and the SWM. The main emphasis of these experimental studies was on long cortico-cortical association and subcortical projection pathways, leaving the study of U-fibers and the SWM largely uninvestigated. Only a limited number of published tracing studies acknowledged or discussed the anatomy and connectivity of local, short association connections, including those that travel within the superficial portions of the white matter (e.g., Jones et al., 1978; Pandya & Seltzer, 1982; Seltzer & Pandya, 1989; Burton et al., 1995; Petrides & Pandya, 2006, 2007; Yeterian & Pandya, 2010; Reveley et al., 2015). Other tracer studies incidentally illustrated axons traveling within the superficial portions of the white matter, but these findings were typically not discussed in detail (e.g., Friedman & Jones, 1981; Friedman et al., 1986; Cavada & Goldman-Rakic, 1989a, 1989b; Seltzer et al., 1996; Morris et al., 1999; Petrides & Pandya, 2006; Muñoz-López et al., 2015).

Only recently has interest in U-fibers increased. Based on theoretical calculations, Braitenberg and Schüz (1991) estimated that short association tracts within the SWM must comprise a much greater fraction of total cerebral white matter than long-range tracts. This finding highlighted the potential importance of local association fibers and U-fibers for cortical information processing and brought attention to the SWM as a critical component of cortical organization. The advent of diffusion MRI and tractography methods subsequently enabled the non-invasive investigation of fiber pathways in humans and, thus, the evaluation of U-fibers within the SWM. These studies consistently identified U-shaped bundles throughout the brain (e.g., Oishi et al., 2008; Zhang et al., 2014; Guevara et al., 2017; Román et al., 2017), reinforcing the view that U-fibers represent a common and broadly distributed component of cortical white matter organization. However, these studies lacked clear neuroanatomical ground truth against which to evaluate their results, making it difficult to determine whether the observed U-fibers reflected anatomically plausible pathways or constituted false-positive reconstructions. Thus, systematic anatomical evidence on U-fiber prevalence, distribution, and organization across different cortical regions has remained limited.

### 3.3 The Heterogeneous Nature of the U-fiber

One of the major findings of the current study was the relative paucity of classically defined U-fibers. U-fibers are among the most apparent features of the white matter beneath the cerebral cortex and are depicted in gross anatomical dissection studies in humans and non-human primates. Dejerine and Dejerine-Klumpke (1895) indicated that U-fibers were present beneath all sulci as a ubiquitous feature of white matter organization, and U-shaped bundles have been frequently reconstructed in numerous diffusion MRI studies in humans and gyrencephalic monkeys (e.g., Oishi et al., 2011; Zhang et al., 2014; Rojkova et al., 2016; Guevara et al., 2017, 2020; Catani et al., 2017; Román et al., 2017; Pron et al., 2021; Zhang et al., 2024; Chauvel et al., 2024). Thus, based on both classical anatomical descriptions and modern imaging evidence, we expected at the outset of the present study that U-fibers would be frequently observed and would traverse sulci in a uniform fashion. The present results, however, did not support this expectation.

To evaluate this question, we operationally defined U-fibers as fiber bundles that coursed from a gyral crown injection site around a sulcus with a U-shaped trajectory to terminate on the opposing gyral crown. We selected cases with injections on gyri adjacent to a major sulcus and found that U-fibers meeting this criterion were observed in only 35.7% of cases around the major targeted sulci. A review of the macaque connectional neuroanatomical literature found little evidence for a uniform or ubiquitous network of U-fibers beneath the cortex. Instead, the available literature and the findings of the present study suggest that U-fibers are region-specific. This finding is consistent with the idea that U-fiber distribution is related to the specificity of their connection, rather than forming a consistent architectural and geometric feature linking all adjacent gyral areas across a sulcus.

#### 3.3.1 U-fibers Beneath the Central Sulcus are Ordered and Symmetrical

Our finding that U-fibers were consistently identified beneath the central sulcus represents an important exception to this point and is supported in the literature, at both the single-axon and bundle levels. DeFelipe et al. (1986) reported labeled neurons in cortical area 4 (i.e., primary motor cortex) whose principal axons entered the white matter and could be followed caudally beneath the central sulcus toward cortical area 3b. Similarly, Yamashita and Arikuni (2001) reconstructed the axon and axonal branches of an individual pyramidal neuron in the putative forelimb representation of the motor cortex. This axon descended into the white matter and followed a sharply curved trajectory along the rostral bank, fundus, and caudal bank of the central sulcus. Along this course, the main axon gave off a branch along the rostral sulcal bank that entered and traveled through the cortical gray matter to terminate at the fundus (Yamashita & Arikuni, 2001), similar to the terminal field we observed in our central sulcus cases. Comparable results were reported by Künzle (1978), who demonstrated that tritiated amino acid injections into the precentral gyrus produced labeled terminals in specific regions on the postcentral gyrus. U-shaped bundles beneath the central sulcus have also been repeatedly identified in diffusion MRI studies in humans (e.g., Oishi et al., 2011; Catani et al., 2012; Zhang et al., 2014; Rojkova et al., 2016; Guevara et al., 2017, 2020; Román et al., 2017; Pron et al., 2021; Zhang et al., 2024; Chauvel et al., 2024), chimpanzees (e.g., Zhang et al., 2014; Chauvel et al., 2024), and macaque monkeys (e.g., Oishi et al., 2011; Zhang et al., 2014).

The consistent identification of U-fibers beneath the central sulcus, both in the present study and in neuroanatomical tract-tracing studies, relates to the organization of the primary motor cortex on the precentral gyrus and the primary somatosensory cortex on the postcentral gyrus. Each of these regions contains a somatotopic representation at the macroscale level (e.g., Penfield & Boldrey, 1937; Penfield & Rasmussen, 1950; Künzle, 1978; Jones et al., 1978; Germann et al., 2020), and these representations are arranged in a largely parallel manner across the central sulcus. As a result, connections corresponding to specific bodily representations - for example, the arm representation in motor cortex and the corresponding representation in somatosensory cortex - are expected to traverse the central sulcus in a U-shaped trajectory oriented perpendicular to the long axis of the sulcus. Such connections linking these two representations can be expected along much of the length of the central sulcus, giving the appearance of a generalized U-fiber layer, but only to the extent that the underlying functional representations are aligned. The regularity of U-fibers beneath the central sulcus, therefore, may have served as an exemplar that shaped subsequent interpretations of U-fibers as continuous sheets or widespread networks throughout the SWM more generally.

#### 3.3.2 Beyond the Central Sulcus: U-fibers Beneath Other Sulci

This assumption of a continuous U-fiber layer does not hold for all sulci. Unlike the central sulcus, where the cortical regions are arranged in parallel across the sulcus and the somatotopic representations are largely matched across it, regions such as the cortex surrounding the IPS contains a large number of functionally and structurally distinct cortical areas, both within the sulcus and on the adjacent gyral crowns (e.g., Seltzer & Pandya, 1980; Pandya & Seltzer, 1982; Cavada & Goldman-Rakic, 1989a; Blatt et al., 1990; Gamberini et al., 2020; Niu et al., 2020). The cortical areas forming the banks of the IPS and the adjacent gyri do not form uniform connections with one another; rather, each area connects selectively with a subset of neighboring or adjacent areas, many of which are located along the sulcal banks rather than on the opposing gyral crown (e.g., Pandya & Seltzer, 1982; Cavada & Goldman-Rakic, 1989a; Lewis & Van Essen, 2000; Rozzi et al., 2006). The distribution of local short association fiber bundles beneath the IPS must therefore reflect this pattern of selective connectivity. Our findings are consistent with this: U-fibers beneath the IPS were present in a minority of cases, while shorter fiber bundles terminating along the sulcal banks and fundus without reaching the opposing gyrus were more frequent. Of the U-fibers observed, the majority originated from SPL injections and crossed beneath the mid-IPS to terminate in the IPL. Furthermore, because the interconnected cortical areas are not evenly distributed or topographically matched across the sulcus, these U-fibers crossed at an oblique rather than perpendicular angle and were therefore classified as asymmetric. Together, these observations indicate that the organization of short association fiber bundles beneath the IPS reflects the selective connectivity of the cortical areas it separates rather than a uniform U-fiber architecture.

More broadly, our findings indicate that U-fiber connections are regionally specific. Where the representations on opposing gyri are aligned, as at the central sulcus, U-fibers are consistently present and largely symmetric, a finding in accord with classic descriptions (e.g., Penfield & Boldrey, 1937; Penfield & Rasmussen, 1950; Künzle, 1978). Where a sulcus crosses multiple cortical areas, as along the IPS, U-fibers are asymmetric because they interconnect regions that are not aligned across the sulcus. This pattern argues against the view of U-fibers as a continuous network beneath the cortex and supports a model in which their presence and trajectory reflect connectional specificity rather than sulcal geometry. This is particularly important as diffusion MRI studies have reported U-shaped streamlines around the IPS interconnecting the SPL and IPL in the macaque (e.g., Zhang et al., 2014; Catani et al., 2017) and connecting the supramarginal and angular gyri (corresponding to the IPL in macaque) in humans (e.g., Zhang et al., 2014; Burks et al., 2017; Catani et al., 2017; Chauvel et al., 2024) and chimpanzees (e.g., Zhang et al., 2014; Chauvel et al., 2024). Our data confirm the presence of these U-shaped short association fiber bundles around the IPS, but in a more restricted way than the dMRI streamlines suggest.

The anatomical characteristics of the sulcus may also contribute to the prevalence of U-fibers. The principal and central sulci were the least complex and shallowest sulci we analyzed, and both exhibited frequent and symmetric U-fibers. The IPS and STS are the most complex; both are deeper and operculated and span a longer physical distance. U-fibers were observed in rostral portions of the STS and IPS where the sulcus is shallower, and at the mid-portion of the IPS where it is deeper but not yet operculated. No U-fibers were observed beneath the STS or IPS at their operculated portions, and none were found around the lateral fissure, which is the deepest and most operculated sulcus of the brain (Table 1). However, in 40% of temporal cases, U-fibers were present around adjacent minor sulci, such as the inferior occipital and occipitotemporal sulci, rather than the STS itself, and these accounted for most of the U-fibers observed across the temporal cases (Table 1). The apparent association between shallow sulci and U-fibers may itself reflect connectivity: shallower sulci tend to border fewer distinct cortical areas, making direct connections between adjacent gyri more likely.

### 3.4 The Superficial White Matter

The division of the cerebral white matter into superficial and deep compartments is common in modern studies. The superficial layer (i.e., the SWM) is thought primarily to contain short cortico-cortical association fibers that interconnect adjacent and neighboring cortical regions (Van Dyken et al., 2024). By contrast, the deep white matter (DWM) contains ipsilateral and contralateral long cortico-cortical association fibers, as well as projection fibers (e.g., cortico-subcortical, cortico-brainstem, and cortico-spinal).

Despite its widespread use, no clear consensus currently exists on the specific depth at which the SWM transitions into the DWM. Many studies rely on fixed or arbitrary depth thresholds. In studies of the human cerebrum, the depth of the SWM has typically been set at approximately 1-3 mm from the GM-WM border (Table 4). Other approaches have used white matter neuron density to distinguish the superficial and deep compartments but rarely specify an explicit cutoff depth value. This may reflect, in part, the considerable variation in neuronal density between gyral and sulcal regions, which complicates the use of any single threshold (e.g., García-Marín et al., 2010; Mortazavi et al., 2017; Sedmak & Judaš, 2019). Still other studies have emphasized axonal orientation as a defining factor of the SWM-DWM border (e.g., Zikopoulos & Barbas, 2010; Schilling et al., 2018) or have identified the SWM as a subplate analog (Kostović et al., 2014). The variety of metrics employed reflects the difficulty of defining the SWM-DWM transition by any single measure. Indeed, a recent study that evaluated multiple metrics in the human brain argues that the SWM-DWM border is better understood as a continuum rather than a discrete anatomical line (Hwang et al., 2026).

**TABLE 4.** Summary of superficial white matter depths in different studies.

| Paper | Species | Approach/modality | Depth | Gyri/Sulci |
| --- | --- | --- | --- | --- |
| Von Monakow (1905) | Human | Dissection/Histological | 3 mm | - |
| Akbarian et al. (1993) | Human | Histology (neurons) | 800 μm (basis of delineation not clear) | Frontal lobe; gyral and sulcal |
| Schüz and Braitenberg (2002) | Human | Estimated thickness/cross-sectional area | 1.5 mm | Depth of sulci |
| García-Marín et al. (2010) | Human | White matter (WM) neurons | 0.175 mm (basis of delineation not clear) | Frontal, cingulate, temporal, and visual cortices; sulcal bank |
| Zikopoulos and Barbas (2010) | Human | Axonal orientation | 2 mm | Prefrontal cortex; gyral |
| Kostović et al. (2014) | Human | Histology, T2w MRI | Not specified; see Figs 8-11, 13-15 | Frontal, parietal, and occipital; gyri and sulci depicted |
| Reveley et al. (2015) | Macaque | Myelin stain, diffusion MRI | ≤1 mm (operationally defined); see Fig 4 (myelin-stained section ∼250 μm) | Frontal and parietal; gyri and sulci depicted |
| Wu et al. (2016) | Human | T1w and Magnetization Transfer MRI, (vertex-based surface statistics) | 1-5 mm below GM-WM boundary (operational sampling window; rationale = reduce partial volume) | Throughout the brain; gyral vs sulcal not specified |
| Mortazavi et al. (2017) | Macaque | WM neurons | 0.4 mm (basis of delineation not clear) | Frontal, temporal, and parietal cortices; gyral and sulcal |
| Zikopoulos et al. (2018) | Macaque and Human | Radial alignment of axons in SWM, sagittal axons in DWM | 0.5-2.5 mm | Prefrontal; gyrus vs sulcus not specified |
| Sedmak and Judaš (2019) | Human | WM neurons | 3 mm (based on von Monakow segment IV) | Frontal, cingulate, temporal, and occipital cortices; density differences between gyri and sulci discussed |
| Borra et al. (2020) | Macaque | WM neurons | 0.5 mm (basis of delineation not clear) | Frontal, cingulate, insular, and parietal cortices; gyrus vs sulcus not specified |
| Kirilina et al. (2020) | Human | T1w and PDw MRI (iron), Immunohistochemistry | 0.5 mm | Primarily temporal lobe; sulci |
| Yoshino et al. (2020) | Ferret | Histology, Dil tracer | Not specified; ~250–300 μm in Figs 1-2, 4 | Region not specified; sulcal only |
| Sedmak and Judaš (2021) | Human | WM neurons | 3 mm (based on von Monakow segment IV) | Density differences between gyri and sulci discussed |
| Zouridakis et al. (2023) | Human | WM neurons | 2 mm (based on Sedmak and Judaš 2021) | Frontal, temporal, and occipital regions; gyrus only |
| Schilling et al. (2023) | Human | Diffusion and T1w MRI | 2-3.5 mm | Throughout the brain; gyral and sulcal |
| Lee et al. (2023) | Human | MRI-based susceptibility source separation (iron vs myelin) | ~0.5-2.0 mm (operationally defined relative to GM-WM boundary and local cortical thickness) | Frontal regions; sulci only |
| Dannhoff et al. (2024) | Human | Klingler dissection, T2w MRI | 0.6-2.9 mm | Temporal lobe; sulcal |
| Kimura et al. (2025) | Macaque | Multimodal MRI (vasculature) | 0.32 mm (based on Kirilina et al., 2020) | Throughout the brain; gyral vs sulcal not specified |
| Tian et al. (2025) | Human | Not specified | 1.5 mm (rationale not explicit) | Not specified |
| Hwang et al. (2026) | Human | Histology, T1w and Magnetization Transfer MRI | 2 mm; evidence for gradient of features | Throughout the brain; gyral vs sulcal discussed |

The present study approaches this question from a connectional neuroanatomical perspective and provides strong evidence for a distinct superficial band approximately 200-300 µm beneath the cerebral cortex. This band was identified in two complementary ways: by the presence of labeled fibers that coursed within it when no labeled fibers were present beneath it, and by the absence of labeling in this layer when only fibers beneath this band were labeled. In both instances, labeled fibers only entered this band to terminate in the overlying cortex. We identified no cases in which anterogradely labeled fibers traveled within this layer and then exited into the white matter below. Thus, this 200-300 µm white matter band immediately beneath the cortex appears to represent a zone of axons ‘on approach’ to their cortical terminations.

It is therefore reasonable to ask whether this 200-300 µm superficial band corresponds to what has traditionally been termed the SWM. While this band clearly lies within the SWM and is consistent with depths reported in some previous studies (Table 4), our current data are insufficient to conclude that this band defines the full extent of the SWM. It is equally likely that the SWM encompasses this band together with additional fibers located subjacent to it. Consistent with this, U-fibers were present below this band. However, labeled fibers from cord fibers and long cortico-cortical association pathways, such as the superior longitudinal fasciculus (Petrides & Pandya, 2002; Schmahmann & Pandya, 2006) were also identified below this band. Therefore, the presence of mixed fiber pathways suggests that the region below the superficial band may not belong exclusively to either a local or long-range compartment, or that the boundary between superficial and deep white matter is potentially not uniform across the cortex. Thus, as discussed above and articulated by Hwang et al. (2026), the apparent width of the SWM depends on the metric used to define it. Although the results of our connectional approach identify a well-defined band characterized by highly local connectivity, this band does not preclude a more expansive and graded definition of the SWM. Further research will be required to determine how this band relates to other defining features of the SWM, including axonal orientation, MRI contrast, and regional variation in white matter neuronal density.

### 3.5 The Importance of a Precise Delineation of the GM-WM Border

In the present study, precise delineation of the GM-WM border was essential for detecting the distinct superficial white matter band immediately beneath the cortex (e.g., dashed yellow lines in Figures 14B, D and 17B, D, F). This was achieved by identifying the GM-WM border in brightfield images used to visualize the Nissl counterstain and transferring this border to the corresponding darkfield images of the same section, allowing the position of labeled fibers to be evaluated directly relative to the cortex. This band has not been explicitly identified in prior connectional neuroanatomical studies, likely due in part to a historical emphasis on long-range pathways and their organization within the white matter. In addition, many depictions of neuroanatomical results rely on schematic or template-based representations of white matter organization, which, while critical for identifying common patterns across cases, may also obscure fine-scale features that depend on accurate local measurements relative to the GM-WM border.

An additional factor that has complicated the interpretation of SWM organization and the identification of U-fibers is the extensive horizontal intracortical fiber network within cortical layer 6, consisting of axons and axon collaterals that travel parallel to the cortical surface for approximately 8 mm before terminating (Rockland & Drash, 1996; Pucak et al., 1996; Yamashita & Arikuni, 2001; Rockland & Knutson, 2001; Rockland, 2020). The intracortical axonal plexus can easily be mistaken for U-fibers when the GM-WM border is not precisely defined, as labeled axons cross this border to enter layer 6, travel within it, and terminate there. Conversely, axons within the layer 6 intracortical network may not strictly respect the GM-WM boundary and may intermittently extend into the subjacent white matter. Together, these interactions underscore the need for careful boundary definition when interpreting the axonal trajectories of short association fibers near the GM-WM interface.

#### 3.5.1 Superficial and Deeper U-fibers versus U-shaped bundles: U-shaped is different than U-fibers

The anatomical term ‘U-fiber’ refers to the fiber bundle’s characteristic U-shaped morphology and its superficial topographic location at the depth of a sulcus as observed in gross dissection of the brain (e.g., Arnold, 1883; Meynert, 1885; Dejerine & Dejerine-Klumpke, 1895). However, the precise depth of these U-fibers beneath the cortex is difficult to evaluate using anatomical dissection techniques, as these techniques are destructive in nature and are unable to preserve the GM-WM border. Without a clearly delineated GM-WM border as a fiducial reference, it is not possible to determine the position of a fiber bundle in the white matter. The precise delineation of the GM-WM border in the present study therefore enabled a depth-based evaluation of U-fiber position that is not possible using dissection methods and, to our knowledge has not been performed using tract tracing.

Precise delineation of the GM-WM border in the present study allowed us to directly examine a key feature that has guided many dMRI interpretations: the conception that U-fibers occupy the most superficial portions of the white matter (e.g., Guevara et al., 2020). However, in our material, U-fibers were morphologically diverse. Some traveled within the superficial band and interconnected adjacent gyri (‘true’ U-fibers). A separate population of deeper U-fibers traveled below the superficial band and interconnected adjacent gyri. A third population of longer U-shaped bundles also traveled below the superficial band but did not interconnect adjacent gyri. These observations are consistent with recent descriptions of longer cortico-cortical or thalamo-cortical bundles that have U-shapes in part of their trajectories (e.g., Xu et al., 2021; Schilling et al., 2025). Together, these findings support two distinctions: first, U-fibers are not synonymous with the most superficial portions of the white matter, and second, several types of fiber bundles take a U-shaped course, not only the U-fibers that interconnect adjacent gyri.

### 3.6 Association fibers Shorter than U-fibers

In addition to U-fibers, we identified shorter fiber bundles that did not extend from the gyral injection site to terminate on the adjacent gyrus. Rather than reaching the opposing gyral crown, these fibers terminated along the proximal sulcal bank or fundus (L-type) or hooked around the sulcal fundus to terminate on the opposing bank (J-type). These L- and J-type connections were not evenly distributed across the cortex, appearing more frequently at deeper and often opercular sulci such as the IPS and STS that intersect multiple distinct cortical areas. This suggests that at deeper sulci intersecting multiple distinct cortical areas, local gyral connectivity more commonly takes the form of connections terminating along the sulcus rather than connections with the adjacent gyrus. Together, these findings indicate that the SWM contains multiple classes of short association fiber bundles rather than a single canonical U-fiber system, a complexity not fully captured by classical descriptions of SWM organization.

Meynert’s rule proposes that fiber bundle length increases with distance from the overlying cortex. Consistent with this, these highly local fibers were positioned more superficially in the white matter, directly beneath the GM-WM border: 100% of L-type bundles (14/14) observed across all cases traveled within the white matter directly beneath the GM-WM border, while J-type bundles did so in 83.3% (5/6) of case. When both connection types were present beneath the same sulcus (e.g., case BMX; Figure 7C-D), the shorter L-type bundle occupied the white matter directly beneath the GM-WM border, while the J-type bundle coursed deeper and traveled farther around the sulcus. This superficial position and curved trajectory along the sulcus have potential implications for diffusion MRI tractography. J-type connections in particular follow a trajectory similar to U-fibers, curving beneath the fundus before terminating on the opposing sulcal bank rather than the opposing gyral crown. Tractography algorithms may therefore misinterpret such fibers, potentially contributing to false-positive U-fiber reconstructions. Conversely, the shorter L-type connections, which do not terminate beyond the fundus, may be susceptible to being underrepresented or missed entirely due to gyral bias (e.g., Schilling et al., 2018). Both L- and J-type bundles represent a potential bottleneck in which streamlines passing through the SWM may be difficult to disambiguate. Although the presence of L- and J-type fibers has been suggested in diffusion MRI-based investigations (e.g., Chauvel et al., 2024), to our knowledge the present study provides the first direct identification of these fibers using connectional neuroanatomical methods.

The L- and J-type connections identified here likely represent only a subset of short association fiber bundle types present within the superficial portions of the white matter, as our cases involved only gyral injections. For example, Blatt et al. (1990) reported terminations on the medial IPS bank following lateral bank injections, and their illustrations suggest labeled fibers coursed around the sulcal fundus in a curved trajectory, resembling J-type connections and U-fibers at the depth of the sulcus but originating from a sulcal bank rather than a gyral crown injection. Such fiber bundles would represent a distinct type not captured in the present dataset. Furthermore, evidence for reciprocal connectivity between the IPS banks and the IPL crown has also been reported using bidirectional tracer injections (e.g., Cavada & Goldman-Rakic, 1989a), suggesting that fibers projecting from the banks to the gyral crown may produce shapes resembling L- or J-type connections. Together, these observations suggest that additional types of short association bundles likely exist, and future studies focused on fiber bundles from sulcal bank injections will be necessary to fully characterize the diversity of these types.

### 3.7 Limitations

Because the tissue was processed following paraffin embedding, some degree of tissue shrinkage is expected (e.g., Kretschmann et al., 1982). This may influence the absolute depth measurements and should be considered when the 200-300 µm superficial band is compared to estimates derived from tissue processed differently in the existing literature (Table 4). Nonetheless, the consistency of this band across cases and sulci suggests that the relative architectural organization described here is unlikely to be an artifact of processing.

Beyond this methodological consideration, our conclusions are subject to four limitations. First, the analysis was restricted to a subset of major sulci, and other sulci were not systematically evaluated. Although labeling around other sulci was catalogued when present in our cases, future work should extend this approach more broadly across the cortex to include additional sulci. Second, our analysis focused primarily on U-fibers and other local short association fibers that originate from a gyrus and terminate along or across the adjacent sulcus and was not designed to evaluate intragyral fibers that terminate locally within the injected gyrus, which are more difficult to follow in autoradiographic material. Intragyral fibers are increasingly recognized as an important component of local cortical connectivity (Dannhoff et al., 2023), and future studies that apply similar connectional approaches to intragyral fiber systems will be necessary to more fully characterize SWM organization.

Third, these findings are based on material from the macaque monkey. Differences in cortical expansion, gyrification, and relative white matter volume may limit direct translation of these findings to human SWM architecture. Both the gray matter and white matter are expanded in humans relative to other primate species (Rilling & Insel, 1999; Donahue et al., 2018), and this expansion could be accompanied by a disproportionate increase in short association fibers relative to the macaque monkey brain. Furthermore, given the greater overall white matter volume and cortical thickness in humans, the absolute depth of the 200-300 µm superficial white matter band identified here, defined based on labeled fibers that either traveled within the band or deeper to it, may differ from that observed in the macaque. Experimental tract-tracing methods cannot be performed in humans, so confirmation of these features in human SWM will require methods that can resolve them non-invasively.

Finally, our analysis was limited to cases with cortical injections on gyral crowns, which allowed us to relate the frequency of U-fibers to individual sulci. However, this approach also constrained the fiber types that could be identified and did not account for fibers originating from sulcal banks or fundus. This limitation reflects, in part, the fact that tract-tracing studies in the connectional literature have predominantly targeted gyri rather than sulcal banks or the fundus (e.g., Jones et al., 1978; Friedman et al., 1986; Seltzer & Pandya, 1989; Blatt et al., 1990; Darian-Smith & Darian-Smith, 1993; Felleman et al., 1997; Cipolloni & Pandya, 1999; Lewis & Van Essen, 2000; Nakamura et al., 2001; Mohedano-Moriano et al., 2005; Reveley et al., 2015). This represents a significant gap in our understanding of white matter organization, as the fiber pathways that originate from sulcal banks and the fundus may differ in their trajectories and relationship to the superficial white matter band identified in the present study.

### 3.8 Implications for Modern Interpretations of the U-fiber System

To our knowledge, a detailed connectional analysis of the local association fibers within the SWM has not yet been performed, despite extensive investigations of the SWM in the human and macaque brain using dissection and imaging methods (e.g., Oishi et al., 2008; Zikopoulos & Barbas, 2010; Reveley et al., 2015; Román et al., 2017; Zikopoulos et al., 2018; Guevara et al., 2020; Shinohara et al., 2020; Kirilina et al., 2020; Schilling et al., 2023a; Xue et al., 2023; Schilling et al., 2023b; Dannhoff et al., 2024; Nie et al., 2024; Van Dyken et al., 2024; Zhang et al., 2024; Schilling et al., 2025; Wang et al., 2025; Hwang et al., 2026). The current study indicates that the SWM is more complex and contains more fiber bundle classes than previously appreciated.

This study has implications on understanding how the more limited U-fiber distribution observed here relates to the widespread detection of U-shaped bundles in gross anatomical dissection and diffusion MRI-based studies. Both techniques have a limited ability to fully resolve crossing or interdigitating fibers within the SWM (Reveley et al., 2015), so anatomically distinct pathways may appear to have a continuous U-shaped architecture, particularly when short and long fibers are in close proximity under a sulcus. The superficial position and curved trajectories of L- and J-type connections and potentially bank-to-bank connections (e.g., Figures 8, 11 of Blatt et al., 1990) may further contribute to this appearance, as such fibers could be incorrectly reconstructed as U-shaped streamlines in tractography given the documented gyral bias of tractography (e.g., Schilling et al., 2018). Our current findings suggest that at least some of the apparent ubiquity of U-fibers reported in prior work may reflect methodological limitations. In addition, the present results strongly support the conclusion that U-fibers are present where specific connections exist, rather than forming a diffuse or uniform component of the SWM, and sulcal white matter.

The present findings have several direct implications for diffusion MRI tractography. First, the distinction between superficial and deeper U-fibers identified here has important consequences for streamline termination and seeding strategies. Superficial (‘true’) U-fibers, which course within the 200-300 µm band directly beneath the GM-WM border, and deeper U-fibers, which avoid this band until approaching their cortical terminations, represent distinct subpopulations of gyrus-to-gyrus connections in terms of their depth within the white matter. This complexity is further compounded by the presence of fiber bundles that appear U-shaped beneath sulci but did not interconnect opposing gyri, which traveled at a similar depth from the GM-WM border as the deeper U-fibers. Tractography algorithms that treat the SWM as a homogeneous zone may not adequately reflect this complexity and lead to errors in reconstructing bundles passing in or through the SWM.

Second, we observed association fiber bundles that had a U-shaped trajectory beneath sulci but did not connect opposing gyri. Such U-shaped trajectories are not exclusive to cortico-cortical association fibers, as depictions of thalamocortical fibers indicate that as they approach their cortical terminations, the distal portions of these fibers and bundles adopt U-shaped trajectories beneath sulci (e.g., Friedman & Jones, 1981; Xu et al., 2021). These observations suggest that U-shaped morphology alone is insufficient to identify a fiber as a short association U-fiber.

Finally, U-fibers are not evenly distributed across sulci, but are present only when specific connections between adjacent gyri exist. Tractography studies that report U-shaped streamlines beneath all major sulci should be interpreted with caution, as such streamlines may reflect methodological limitations or false-positive reconstructions rather than true anatomical connections.

#### 3.8.1 Clinical Relevance

Short association fibers in the SWM, including those that form U-fibers, are among the last axons to undergo myelination and play a role in brain plasticity, development, and aging (e.g., Wu et al., 2014). However, this protracted myelination may also increase their vulnerability to damage during brain development (Wu et al., 2014). Although the architecture and connectivity of the SWM remain poorly characterized, local association fibers and the SWM have been shown to be sensitive to certain disorders or trauma (Guevara et al., 2020; Van Dyken et al., 2024). In autism spectrum disorder (ASD), an increase in thinly myelinated SWM axons beneath the anterior cingulate cortex (Zikopoulos & Barbas, 2010) and changes in axonal myelination below the orbitofrontal cortex have been reported (Zikopoulos & Barbas, 2013). In chronic traumatic encephalopathy (CTE), the SWM at the depths of sulcal folds is affected more severely and earlier than other regions of the brain (e.g., McKee et al., 2009; Holleran et al., 2017). Recent studies have also demonstrated progressive sulcal neuronal loss within cortical layers 2/3 that accumulates with repeated head trauma exposure (Butler et al., 2025). Because layer 2/3 neurons are critical in intercortical processing, neuronal loss in the sulcal cortex may preferentially affect the shorter L- and J-type connections that terminate along sulcal banks.

Diffusion MRI tractography is widely used to evaluate SWM integrity in neurological and psychiatric conditions. Diffusion MRI studies have revealed lower fractional anisotropy and connectivity deficits in SWM bundles that correlate with symptom severity in ASD across frontal, temporal, and parietal regions (d’Albis et al., 2018). Increases in diffusivity that correlate with cognitive decline have been reported across the SWM in Alzheimer’s disease (AD), with particularly pronounced changes in the parahippocampal regions and the temporal and frontal lobes (Phillips et al., 2016a). In Huntington’s disease, increases in diffusivity across the SWM have been observed even in presymptomatic individuals and expand to include the whole brain in symptomatic patients, with changes correlating with genotype and disease burden, suggesting this pattern tracks disease progression from its earliest stages (Phillips et al., 2016b).

These findings highlight the broad clinical relevance of the SWM across neurological and psychiatric conditions (see Van Dyken et al. (2024) and Guevara et al. (2020) for reviews). However, the ability to detect, localize, and interpret SWM changes using diffusion MRI depends on the accuracy of the underlying anatomical model of the SWM. Our results indicate that the effect of these disorders on the white matter is likely more complex than presently appreciated. Accurate characterization of SWM pathology requires methods that can distinguish between the short association fiber systems identified here. Many diffusion MRI tractography studies currently treat the SWM as a single undifferentiated system of U-fibers, and this simplification may obscure clinically meaningful differences. Higher-resolution dMRI scans may produce more refined tractography and help better resolve the SWM (Zhang et al., 2024). However, increases in spatial resolution increase the number of voxels with complex fiber architectures (Schilling et al., 2017), which dMRI tractography struggles to resolve correctly. Understanding how fibers within the SWM are organized is therefore essential not only for characterizing short association connectivity itself, but also for accurately tracing long-range pathways, as the SWM has been shown to impede the reconstruction of streamlines crossing into and out of the cortex in more than half of the cortical hemisphere in the macaque (Reveley et al., 2015), particularly along sulci (Schilling et al., 2018). Advances in dMRI tractography approaches, such as seeding within the cortical gray matter, therefore depend on enhancing our understanding of the neuroanatomical organization of U-fibers and the SWM.

## 4. Conclusions

The classical model of the superficial white matter, articulated by Meynert (1885) and the Dejerines (Dejerine & Dejerine-Klumpke, 1895, 1901) from gross dissection and histology, describes a uniform sheet of symmetric U-fibers beneath all sulci interconnecting adjacent gyri. Using experimentally labeled fibers whose depth was assessed relative to a precisely delineated GM-WM border, we found that this model does not hold. U-fibers are neither ubiquitous nor uniformly organized; their presence, trajectory, and symmetry varied across cortical regions, and they coexisted with shorter L- and J-type bundles that terminate along the sulcal banks and fundus rather than on the adjacent gyrus. Furthermore, we found that U-shaped bundles beneath sulci are not limited to short association fibers but may also occur in long cortico-cortical association tracts whose trajectories are deflected by the presence of sulci.

At the same time, our connectional approach revealed a laminar arrangement within the superficial portions of the white matter: a consistent 200-300 µm band immediately beneath cortical layer 6 that contained short association fiber bundles. This band was present beneath all sulci examined, and long cortico-cortical association fibers avoided it. U-fibers that interconnected opposing gyri across a sulcus did not always course within this superficial white matter band but could also be observed directly beneath this band. Together with the regional specificity of U-fibers and the presence of multiple classes of short association fiber bundles, this indicates that the white matter beneath the cortex is not a uniform sheet of U-fibers, but a complex system that reflects patterns of short cortico-cortical association connectivity. Essential next steps include extending this connectional analysis to additional cortical regions to better define the regional specificity of short association connections, characterizing the connections that arise from the sulcal banks and fundus, and examining their counterparts in the human brain using non-invasive approaches such as diffusion MRI. The present study provides the first systematic connectional account of the complexity of U-fibers and the SWM in a gyrencephalic primate and offers a foundation for this work.

## 5. Materials and Methods

### 5.1 Methods Overview

The present study analyzed anterograde autoradiographic tract-tracing material from a collection of approximately 500 rhesus macaque tract-tracing cases generated in the laboratories of Dr. Deepak Pandya and Dr. Douglas Rosene, the Pandya-Rosene Neuroanatomical Archive. In these cases, tritiated amino acids were injected into a defined cortical area, and brains were subsequently sectioned in the coronal plane, with sections collected into multiple regularly spaced interleaved (interrupted) series. Sections were processed for autoradiography (Cowan et al., 1972) and counterstained for Nissl substance, allowing labeled fibers and terminal fields to be interpreted relative to cortical cytoarchitecture within the same section. Using this material, SWM connectivity, U-fiber distribution, and related short association fiber bundles were examined in 28 cases around four major sulci: the principal, central, intraparietal, and superior temporal sulci.

### 5.2 Pandya-Rosene Neuroanatomical Archive

The Pandya-Rosene Neuroanatomical Archive contains experimental tract-tracing cases in the rhesus macaque monkey that employed standard anterograde autoradiography techniques for tract tracing. In these cases, a radiolabeled cocktail of tritiated amino acids (^3^H-leucine, ^3^H-proline, ^3^H-lysine, or mixtures of these) was injected into a specific cortical or subcortical area, after which labeled proteins were transported anterogradely along axons. In general, injections consisted of one or two juxtaposed injections per cortical area with a total volume of ∼0.3-1.0 μl containing ∼30-80 μCi of tritiated amino acids in saline. Following a survival interval of ∼5-10 days, animals were deeply anesthetized and perfused with saline followed by a 10% formalin solution. Brains were blocked in situ in the coronal plane, embedded in paraffin, and sectioned in the coronal plane at a thickness of 10 μm. Sections were then processed for autoradiography using a modification of the method described by Cowan et al. (1972), with exposure times of 3 to 6 months, yielding silver grain labeling that visualized labeled axons and their terminal branches. Sections were then counterstained for Nissl substance with thionine to identify cortical architecture. The full methodology has been described in detail in numerous studies (e.g., Petrides & Pandya, 1988, 2007; Demeter et al., 1990; Pandya & Rosene, 1993; Schmahmann et al., 2004; Yeterian & Pandya, 2010).

For our analysis, we selected cases with injections placed on gyri adjacent to the principal, central, intraparietal, and superior temporal sulci. All cases underwent an initial quality assessment and were excluded if they met any of the following criteria: 1) tissue sections exhibited frequent major cutting artifacts that limited our ability to follow labeled fiber trajectories across sections, 2) the Nissl stain was too light to consistently delineate the GM-WM border, or 3) injection placement or tracer spread resulted in either i) labeling in the subjacent white matter or ii) labeling that could not be reliably attributed to one source, such as when the injection foci involved both gyral crowns adjacent to a sulcus or when nonspecific spread produced a halo on the opposite gyral crown or sulcal bank. Following this process, 28 cases spanning the frontal, parietal, and temporal lobes met the inclusion criteria and were analyzed (Table 5). All 28 cases contained the expected labeled fiber components following a successful cortical injection, including cord fibers dividing into commissural and subcortical bundles, cortico-striatal fibers, and long association fiber pathways such as the superior longitudinal fasciculus (Schmahmann & Pandya, 2006). The principal organizational features observed across the dataset, including short association bundle types, symmetry patterns, and depth relationships relative to the GM-WM border, recurred consistently across all 28 cases and are illustrated in the 14 representative cases (Figure 3) described through the current study. All 28 cases contributed to the frequency analyses and summary tables reported here (e.g., Tables 1-2).

**TABLE 5.** Overview of cases and their respective injection sites analyzed from the Pandya - Rosene Neuroanatomical Archive.

| **Case:** | **Lobe/Cortex:** | **Injection Region/Area:** |
| --- | --- | --- |
| BMFF | Prefrontal | Caudal Prefrontal Cortex, Ventral Area 9/46 |
| BMFG | Prefrontal | Caudal Prefrontal Cortex, Dorsal Area 9/46 |
| BMEX-L | Frontal | Precentral Gyrus, M1, Trunk Representation |
| PGR-R | Frontal | Precentral Gyrus, M1, Hand Representation |
| BMJ | Temporal | Rostral Superior Temporal Gyrus, Area Pro/Ts1 |
| BMAW | Temporal | Upper Superior Temporal Sulcus, Area TPO |
| BMBA | Temporal | Rostral Superior Temporal Sulcus, Areas TE<sub>m</sub> and TE<sub>a</sub> |
| BMDV | Temporal | Rostral Inferior Temporal Gyrus, Rostral Area TE1 |
| BMDW | Temporal | Lateral Bank of the Occipitotemporal Sulcus, Areas TF and TE3 |
| BMDD | Temporal | Mid Inferior Lateral Fissure, Areas ProK and KA |
| BMEV | Temporal | Mid Superior Temporal Gyrus, Area paAlt |
| BMFC | Temporal | Mid Inferotemporal Region, Areas TE2 and TE3 |
| PBT | Temporal | Mid Inferior Temporal Gyrus, Areas TE1 and TE2 |
| PDJ | Temporal | Mid Inferotemporal Region, Area TE3 |
| BMD | Parietal | Superior Parietal Lobule, Areas PG<sub>m</sub> and PE<sub>c</sub> |
| BML | Parietal | Inferior Parietal Lobule, Areas PG and Opt |
| BMP | Parietal | Superior Parietal Lobule, Area PE<sub>c</sub> |
| BMT | Parietal | Intraparietal Sulcus, Area PE<sub>c</sub> |
| BMX | Parietal | Rostral Inferior Parietal Lobe, Areas PF and PFG |
| BMAM | Parietal | Ventral Parietal Lobe, Area POG |
| BMAU | Parietal | Caudal Ventral Inferior Parietal Lobe, Area PGOp |
| BMDH | Parietal | Dorsal Opercular Surface of the Lateral Sulcus, Areas PFop and SII |
| MRG | Parietal | Rostral Inferior Parietal Lobe, Area PF |
| BMEQ | Preoccipital | Parieto-Occipital Medial Sulcus, Areas PO and PG<sub>m</sub> |
| BMER | Preoccipital | Preoccipital Dorsal, Area V4d |
| BMFA | Preoccipital | Parieto-Occipital Medial Sulcus, Area 19 |
| BMFD | Preoccipital | Caudal Superior Temporal Sulcus, Areas DP and Upper V4d |
| BMFS | Preoccipital | Annectant Gyrus, Area V3 |

### 5.3 Imaging and Image Processing

Tract-tracing cases from the Pandya-Rosene Neuroanatomical Archive were scanned on a custom Huron TissueScope™ LE120 slide scanner (Huron Digital Pathology, Waterloo, ON, Canada) under brightfield and darkfield illumination using a 10× objective at a native resolution of 0.45 µm/pixel. The brightfield and darkfield imaging passes were acquired sequentially over the same whole-tissue section with identical field dimensions, yielding inherently co-registered images without the need for additional registration.

For all darkfield images, we applied a fixed per-channel histogram adjustment in the native HuronViewer™ to optimize the image and minimize background signal. The first peak in each channel was suppressed (minimum set to ∼20), the red channel’s upper intensity threshold was lowered to ∼160-170, the green and blue channel upper intensity thresholds were lowered to ∼250, and the green channel’s gamma was increased from 1.0 to ∼1.2. After these adjustments, full-color darkfield and brightfield images were downsampled and exported. Each darkfield image was then converted to 8-bit in Fiji/ImageJ software (v2.120/1.54p) (Schindelin et al., 2012) to produce a single grayscale image consolidating the RGB intensity information. Darkfield images were subsequently opened in Adobe Photoshop 2025 (v26.11.0) (Adobe Inc., San Jose, CA, USA). Image adjustments were applied as needed to facilitate visualization of label.

### 5.4 Gray Matter-White Matter Boundary

Each section contained autoradiographic labeling, visualized under darkfield illumination, as well as cytoarchitectural detail visualized under brightfield illumination. To examine the SWM and the depth of fibers relative to the cortex, we manually delineated the GM-WM border on the brightfield image and then transferred this border to the darkfield image (Figure 2) using Adobe Illustrator 2025 (v29.8.2) (Adobe Inc., San Jose, CA, USA). The GM-WM border was delineated on the brightfield image in regions relevant to where labeling was present in the matched darkfield image. In some cases, the line was not drawn directly over the injection site because dense silver grains obscured the cytoarchitecture and the GM-WM border could not be confidently delineated. Border placement was agreed upon by two authors (T.J.C., R.J.R.). Brightfield and darkfield images were imported as separate layers into a single Adobe Illustrator (v29.8.2) document and aligned by overlaying the brightfield image on the corresponding darkfield image. Once the border was delineated on the brightfield image, it was not adjusted further. Because both images were acquired over the same whole tissue section with identical pixel dimensions, the images were aligned without additional warping or registration.

### 5.5 Quantification of Fiber Bundle Depth

We quantified downsampled images (1.8 µm/pixel) to determine the spatial relationship of the white matter bundles with the GM-WM border. For the bundles that were directly beneath the GM-WM border, we measured the depth between the GM-WM border and the deep margin of the bundle (see Figure 11A-C, G). For bundles that were deeper and separated from the GM-WM border by an unlabeled region, we measured the distance between the GM-WM border and the superficial margin of the deeper labeled bundle (see Figure 11D-F, H). Measurement placements were agreed upon by two authors (T.J.C., R.J.R.). Depth measurements were performed in 11/28 cases selected to span the major sulci analyzed and to include both fiber bundle labeling patterns, while minimizing redundancy across cases with similar organizational patterns. Across the 11 cases measured, we sampled three to seven regions of interest (ROIs) per bundle depending on the depth of the sulcus and the length of the fiber bundle. ROIs were placed at the sulcal fundus and on adjacent banks, as appropriate for the bundle being measured, while avoiding any tissue/staining artifacts. We also obtained depth values directly from darkfield intensity profiles sampled orthogonal to the GM-WM border using Fiji. For each case, rectangular ROIs were drawn orthogonal to the local contour of the GM-WM border on the same sections as above (see Figure 11B, E; white boxes). The delineated GM-WM border was retained as a white line so it could be identified on the intensity plot. ROI length extended from the gray matter, through the GM-WM border, and across labeled fibers. Mean gray values were extracted across ROI depth, and depth measurements were made from the GM-WM border. Mean gray values were also converted to squared-intensity profiles to confirm the selected local maximum.

### 5.6 Statistical Analysis

Quantitative analyses of fiber bundle depth were performed on ROI-based measurements derived from darkfield intensity profiles (see Methods: Quantification of Fiber Bundle Depth). Each rectangular ROI contributed a single depth value; within each case, ROIs were treated as repeated samples of that case.

We compared two fiber bundle depth patterns from the GM-WM border: one at the deep margin of labeled superficial bundles (‘no-gap’), and one at the superficial margin of the labeled deeper bundles (‘gap’). Normality within each group was assessed with the Shapiro-Wilk test (No-gap: p = 0.659; Gap: p = 0.360). For the statistical comparison between the mean depths of these two groups, we used an independent-samples Welch’s t-test following a positive Levene’s test for unequal variances (F(1,8) = 5.81, p = 0.019).

For the sulcal-level comparison, ROIs were grouped by sulcus (principal, central, rostral intraparietal, mid intraparietal, rostral superior temporal, caudal superior temporal, and inferior occipital) and we conducted a one-way ANOVA with sulcus as the factor. Shapiro-Wilk tests indicated approximately normal distributions by sulcus (only mid intraparietal showed a modest deviation; p = 0.024). Given the small sample sizes and the robustness of ANOVAs to mild non-normality, we proceeded with a one-way ANOVA. Homogeneity of variance was evaluated with Levene’s test; as variances were unequal across sulci (F(6, 51) = 2.69, p = 0.024), post hoc pairwise comparisons used the Games-Howell test. Descriptive statistics for fiber bundle depths are reported as mean ± standard deviation (SD) and range. Statistical significance was set at α = 0.05 for all test. Analyses were performed in IBM SPSS Statistics, version 29.0.2.0 (IBM Corp., Armonk, N.Y., USA).

## Supporting information

Supplementary Case Atlas

## Acknowledgments/Funding

R01NS125307 (RJR, NM, DLR)

R01MH125860 (LJO, YR, NM)

## References

Akbarian, S., Bunney, W. E., Jr, Potkin, S. G., Wigal, S. B., Hagman, J. O., Sandman, C. A., & Jones, E. G. (1993). Altered distribution of nicotinamide-adenine dinucleotide phosphate-diaphorase cells in frontal lobe of schizophrenics implies disturbances of cortical development. Archives of General Psychiatry, 50(3), 169–177. 10.1001/archpsyc.1993.01820150007001

Amaral, D. G., & Price, J. L. (1984). Amygdalo-cortical projections in the monkey (Macaca fascicularis). The Journal of Comparative Neurology, 230(4), 465–496. 10.1002/cne.902300402

Arnold, F. (1883). Tabulae Anatomicae Fasciculus I. Continens icones cerebri et medullae spinalis. Orell Füssli Verlag.

Assem, M., Shashidhara, S., Glasser, M. F., & Duncan, J. (2024). Basis of executive functions in fine-grained architecture of cortical and subcortical human brain networks. Cerebral Cortex (New York, N.Y.: 1991), 34(2), bhad537. 10.1093/cercor/bhad537

Barbas, H., & Pandya, D. N. (1989). Architecture and intrinsic connections of the prefrontal cortex in the rhesus monkey. The Journal of Comparative Neurology, 286(3), 353–375. 10.1002/cne.902860306

Blatt, G. J., Andersen, R. A., & Stoner, G. R. (1990). Visual receptive field organization and cortico-cortical connections of the lateral intraparietal area (area LIP) in the macaque. The Journal of Comparative Neurology, 299(4), 421–445. 10.1002/cne.902990404

Blatt, Gene J., Pandya, D. N., & Rosene, D. L. (2003). Parcellation of cortical afferents to three distinct sectors in the parahippocampal gyrus of the rhesus monkey: An anatomical and neurophysiological study. The Journal of Comparative Neurology, 466(2), 161–179. 10.1002/cne.10866

Borges, K. C. M., Nishijo, H., Aversi-Ferreira, T. A., Ferreira, J. R., & Caixeta, L. F. (2015). Anatomical study of intrahemispheric association fibers in the brains of Capuchin monkeys (Sapajus sp.). BioMed Research International, 2015, 648128. 10.1155/2015/648128

Borra, E., Luppino, G., Gerbella, M., Rozzi, S., & Rockland, K. S. (2020). Projections to the putamen from neurons located in the white matter and the claustrum in the macaque. The Journal of Comparative Neurology, 528(3), 453–467. 10.1002/cne.24768

Braitenberg, V., & Schüz, A. (1991). Anatomy of the cortex: Statistics and geometry. Springer.

Brodmann, K. (1909). Vergleichende Lokalisationslehre der Gro hirnrinde in ihren Prinzipien dargestellt auf Grund des Zellenbaues. Barth.

Budde, M. D., & Annese, J. (2013). Quantification of anisotropy and fiber orientation in human brain histological sections. Frontiers in Integrative Neuroscience, 7, 3. 10.3389/fnint.2013.00003

Bullmore, E., & Sporns, O. (2012). The economy of brain network organization. Nature Reviews. Neuroscience, 13(5), 336–349. 10.1038/nrn3214

Burks, J. D., Boettcher, L. B., Conner, A. K., Glenn, C. A., Bonney, P. A., Baker, C. M., Briggs, R. G., Pittman, N. A., O’Donoghue, D. L., Wu, D. H., & Sughrue, M. E. (2017). White matter connections of the inferior parietal lobule: A study of surgical anatomy. Brain and Behavior, 7(4), e00640. 10.1002/brb3.640

Burton, H., Fabri, M., & Alloway, K. (1995). Cortical areas within the lateral sulcus connected to cutaneous representations in areas 3b and 1: a revised interpretation of the second somatosensory area in macaque monkeys. The Journal of Comparative Neurology, 355(4), 539–562. 10.1002/cne.903550405

Butler, M. L. M. D., Pervaiz, N., Breen, K., Calderazzo, S., Ypsilantis, P., Wang, Y., Breda, J. C., Mazzilli, S., Nicks, R., Spurlock, E., Hefti, M. M., Fiock, K. L., Huber, B. R., Alvarez, V. E., Stein, T. D., Campbell, J. D., McKee, A. C., & Cherry, J. D. (2025). Repeated head trauma causes neuron loss and inflammation in young athletes. Nature, 647, 228–237. 10.1038/s41586-025-09534-6

Catani, M. (2025). The brain and its pathways. In F. Dell’Acqua, M. Descoteaux, & A. Leemans (Eds.), Handbook of Diffusion MR Tractography (pp. 3–13). Elsevier. 10.1016/b978-0-12-818894-1.00034-3

Catani, M., Dell’acqua, F., Vergani, F., Malik, F., Hodge, H., Roy, P., Valabregue, R., & Thiebaut de Schotten, M. (2012). Short frontal lobe connections of the human brain. Cortex; a Journal Devoted to the Study of the Nervous System and Behavior, 48(2), 273–291. 10.1016/j.cortex.2011.12.001

Catani, M., Robertsson, N., Beyh, A., Huynh, V., de Santiago Requejo, F., Howells, H., Barrett, R. L. C., Aiello, M., Cavaliere, C., Dyrby, T. B., Krug, K., Ptito, M., D’Arceuil, H., Forkel, S. J., & Dell’Acqua, F. (2017). Short parietal lobe connections of the human and monkey brain. Cortex; a Journal Devoted to the Study of the Nervous System and Behavior, 97, 339–357. 10.1016/j.cortex.2017.10.022

Cavada, C., & Goldman-Rakic, P. S. (1989a). Posterior parietal cortex in rhesus monkey: I. Parcellation of areas based on distinctive limbic and sensory corticocortical connections. The Journal of Comparative Neurology, 287(4), 393–421. 10.1002/cne.902870402

Cavada, C., & Goldman-Rakic, P. S. (1989b). Posterior parietal cortex in rhesus monkey: II. Evidence for segregated corticocortical networks linking sensory and limbic areas with the frontal lobe. The Journal of Comparative Neurology, 287(4), 422–445. 10.1002/cne.902870403

Chauvel, M., Pascucci, M., Uszynski, I., Herlin, B., Mangin, J.-F., Hopkins, W. D., & Poupon, C. (2024). Comparative analysis of the chimpanzee and human brain superficial structural connectivities. Brain Structure & Function, 229(8), 1943–1977. 10.1007/s00429-024-02823-2

Cipolloni, P. B., & Pandya, D. N. (1989). Connectional analysis of the ipsilateral and contralateral afferent neurons of the superior temporal region in the rhesus monkey. The Journal of Comparative Neurology, 281(4), 567–585. 10.1002/cne.902810407

Cipolloni, P. B., & Pandya, D. N. (1999). Cortical connections of the frontoparietal opercular areas in the rhesus monkey. The Journal of Comparative Neurology, 403(4), 431–458. 10.1002/(sici)1096-9861(19990125)403:4%3C431::aid-cne2%3E3.0.co;2-1

Corbetta, M. (2012). Functional connectivity and neurological recovery. Developmental Psychobiology, 54(3), 239–253. 10.1002/dev.20507

Cowan, W. M., Gottlieb, D. I., Hendrickson, A. E., Price, J. L., & Woolsey, T. A. (1972). The autoradiographic demonstration of axonal connections in the central nervous system. Brain Research, 37(1), 21–51. 10.1016/0006-8993(72)90344-7

d’Albis, M.-A., Guevara, P., Guevara, M., Laidi, C., Boisgontier, J., Sarrazin, S., Duclap, D., Delorme, R., Bolognani, F., Czech, C., Bouquet, C., Ly-Le Moal, M., Holiga, S., Amestoy, A., Scheid, I., Gaman, A., Leboyer, M., Poupon, C., Mangin, J.-F., & Houenou, J. (2018). Local structural connectivity is associated with social cognition in autism spectrum disorder. Brain: A Journal of Neurology, 141(12), 3472–3481. 10.1093/brain/awy275

Dannhoff, G., Morichon, A., Smirnov, M., Barantin, L., Destrieux, C., & Maldonado, I. L. (2024). Direct inside-out observation of superficial white matter fasciculi in the human brain. Brain Connectivity, 14(2), 107–121. 10.1089/brain.2023.0050

Dannhoff, G., Poudel, P. P., Bhattarai, C., Kalthur, S. G., & Maldonado, I. L. (2023). Depicting the anatomy of the gyral white matter: ubi sumus? quo vadimus? Brain Communications, 5(5), fcad265. 10.1093/braincomms/fcad265

Darian-Smith, C., & Darian-Smith, I. (1993). Thalamic projections to areas 3a, 3b, and 4 in the sensorimotor cortex of the mature and infant macaque monkey. The Journal of Comparative Neurology, 335(2), 173–199. 10.1002/cne.903350204

de Pasquale, F., Corbetta, M., Betti, V., & Della Penna, S. (2018). Cortical cores in network dynamics. NeuroImage, 180(Pt B), 370–382. 10.1016/j.neuroimage.2017.09.063

Decramer, T., Swinnen, S., van Loon, J., Janssen, P., & Theys, T. (2018). White matter tract anatomy in the rhesus monkey: a fiber dissection study. Brain Structure & Function, 223(8), 3681–3688. 10.1007/s00429-018-1718-x

DeFelipe, J., Conley, M., & Jones, E. G. (1986). Long-range focal collateralization of axons arising from corticocortical cells in monkey sensory-motor cortex. The Journal of Neuroscience: The Official Journal of the Society for Neuroscience, 6(12), 3749–3766. 10.1523/jneurosci.06-12-03749.1986

Dejerine, J., & Dejerine-Klumpke, A. (1895). Anatomie des centres nerveux (Tome I). Paris: Rueff.

Dejerine, J., & Dejerine-Klumpke, A. (1901). Anatomie des centres nerveux (Tome II). Paris: Rueff.

Demeter, S., Rosene, D. L., & Van Hoesen, G. W. (1985). Interhemispheric pathways of the hippocampal formation, presubiculum, and entorhinal and posterior parahippocampal cortices in the rhesus monkey: The structure and organization of the hippocampal commissures. The Journal of Comparative Neurology, 233(1), 30–47. 10.1002/cne.902330104

Demeter, S., Rosene, D. L., & Van Hoesen, G. W. (1990). Fields of origin and pathways of the interhemispheric commissures in the temporal lobe of macaques. The Journal of Comparative Neurology, 302(1), 29–53. 10.1002/cne.903020104

Dermon, C. R., & Barbas, H. (1994). Contralateral thalamic projections predominantly reach transitional cortices in the rhesus monkey. The Journal of Comparative Neurology, 344(4), 508–531. 10.1002/cne.903440403

Dombrowski, S. M., Hilgetag, C. C., & Barbas, H. (2001). Quantitative architecture distinguishes prefrontal cortical systems in the rhesus monkey. Cerebral Cortex (New York, N.Y.: 1991), 11(10), 975–988. 10.1093/cercor/11.10.975

Donahue, C. J., Glasser, M. F., Preuss, T. M., Rilling, J. K., & Van Essen, D. C. (2018). Quantitative assessment of prefrontal cortex in humans relative to nonhuman primates. Proceedings of the National Academy of Sciences of the United States of America, 115(22), E5183–E5192. 10.1073/pnas.1721653115

Felleman, D. J., Burkhalter, A., & Van Essen, D. C. (1997). Cortical connections of areas V3 and VP of macaque monkey extrastriate visual cortex. The Journal of Comparative Neurology, 379(1), 21–47. 10.1002/(sici)1096-9861(19970303)379:1%253C21::aid-cne3%253E3.0.co;2-k

Frankle, W. G., Laruelle, M., & Haber, S. N. (2006). Prefrontal cortical projections to the midbrain in primates: evidence for a sparse connection. Neuropsychopharmacology: Official Publication of the American College of Neuropsychopharmacology, 31(8), 1627–1636. 10.1038/sj.npp.1300990

Friedman, D. P., & Jones, E. G. (1981). Thalamic input to areas 3a and 2 in monkeys. Journal of Neurophysiology, 45(1), 59–85. 10.1152/jn.1981.45.1.59

Friedman, D. P., Murray, E. A., O’Neill, J. B., & Mishkin, M. (1986). Cortical connections of the somatosensory fields of the lateral sulcus of macaques: evidence for a corticolimbic pathway for touch. The Journal of Comparative Neurology, 252(3), 323–347. 10.1002/cne.902520304

Gamberini, M., Passarelli, L., Fattori, P., & Galletti, C. (2020). Structural connectivity and functional properties of the macaque superior parietal lobule. Brain Structure & Function, 225(4), 1349–1367. 10.1007/s00429-019-01976-9

García-Marín, V., Blazquez-Llorca, L., Rodriguez, J. R., Gonzalez-Soriano, J., & DeFelipe, J. (2010). Differential distribution of neurons in the gyral white matter of the human cerebral cortex. The Journal of Comparative Neurology, 518(23), 4740–4759. 10.1002/cne.22485

Germann, J., Chakravarty, M. M., Collins, D. L., & Petrides, M. (2020). Tight coupling between morphological features of the central sulcus and somatomotor body representations: A combined anatomical and functional MRI study. Cerebral Cortex (New York, N.Y.: 1991), 30(3), 1843–1854. 10.1093/cercor/bhz208

Glasser, M. F., Smith, S. M., Marcus, D. S., Andersson, J. L. R., Auerbach, E. J., Behrens, T. E. J., Coalson, T. S., Harms, M. P., Jenkinson, M., Moeller, S., Robinson, E. C., Sotiropoulos, S. N., Xu, J., Yacoub, E., Ugurbil, K., & Van Essen, D. C. (2016). The Human Connectome Project’s neuroimaging approach. Nature Neuroscience, 19(9), 1175–1187. 10.1038/nn.4361

Guevara, M., Guevara, P., Román, C., & Mangin, J. F. (2020). Superficial white matter: A review on the dMRI analysis methods and applications. NeuroImage, 212(2020), 116673. 10.1016/j.neuroimage.2020.116673

Guevara, M., Román, C., Houenou, J., Duclap, D., Poupon, C., Mangin, J. F., & Guevara, P. (2017). Reproducibility of superficial white matter tracts using diffusion-weighted imaging tractography. NeuroImage, 147, 703–725. 10.1016/j.neuroimage.2016.11.066

Haber, S. N., Fudge, J. L., & McFarland, N. R. (2000). Striatonigrostriatal pathways in primates form an ascending spiral from the shell to the dorsolateral striatum. The Journal of Neuroscience: The Official Journal of the Society for Neuroscience, 20(6), 2369–2382. 10.1523/jneurosci.20-06-02369.2000

Hau, J., Sarubbo, S., Houde, J. C., Corsini, F., Girard, G., Deledalle, C., Crivello, F., Zago, L., Mellet, E., Jobard, G., Joliot, M., Mazoyer, B., Tzourio-Mazoyer, N., Descoteaux, M., & Petit, L. (2017). Revisiting the human uncinate fasciculus, its subcomponents and asymmetries with stem-based tractography and microdissection validation. Brain Structure & Function, 222(4), 1645–1662. 10.1007/s00429-016-1298-6

Hilgetag, C. C., & Barbas, H. (2006). Role of mechanical factors in the morphology of the primate cerebral cortex. PLoS Computational Biology, 2(3), e22. 10.1371/journal.pcbi.0020022

Holleran, L., Kim, J. H., Gangolli, M., Stein, T., Alvarez, V., McKee, A., & Brody, D. L. (2017). Axonal disruption in white matter underlying cortical sulcus tau pathology in chronic traumatic encephalopathy. Acta Neuropathologica, 133(3), 367–380. 10.1007/s00401-017-1686-x

Hwang, Y., Rodriguez-Cruces, R., DeKraker, J., Cabalo, D. G., Leppert, I. R., Thevakumaran, R., Tardif, C. L., Rudko, D. A., Paquola, C., Bazin, P.-L., Bernasconi, A., Bernasconi, N., Concha, L., Evans, A. C., & Bernhardt, B. C. (2026). Microstructural profiles of the human superficial white matter and their associations to cortical geometry and connectivity. PLoS Biology, 24(1), e3003629. 10.1371/journal.pbio.3003629

Jones, E. G., Coulter, J. D., & Hendry, S. H. (1978). Intracortical connectivity of architectonic fields in the somatic sensory, motor and parietal cortex of monkeys. The Journal of Comparative Neurology, 181(2), 291–347. 10.1002/cne.901810206

Kimura, I., Hayashi, T., & Autio, J. A. (2025). Linking neuron-axon-synapse architecture to white matter vasculature using high-resolution multimodal MRI in primate brain. Imaging Neuroscience (Cambridge, Mass.), 3, IMAG.a.77. 10.1162/IMAG.a.77

Kirilina, E., Helbling, S., Morawski, M., Pine, K., Reimann, K., Jankuhn, S., Dinse, J., Deistung, A., Reichenbach, J. R., Trampel, R., Geyer, S., Müller, L., Jakubowski, N., Arendt, T., Bazin, P.-L., & Weiskopf, N. (2020). Superficial white matter imaging: Contrast mechanisms and whole-brain in vivo mapping. Science Advances, 6(41), eaaz9281. 10.1126/sciadv.aaz9281

Klingler, J. (1935). Erleichterung der makrokopischen Präparation des Gehirns durch den Gefrierprozess. Schweiz. Arch. Neurol. Psychiatr., 36, 247–256.

Kostović, I., Jovanov-Milošević, N., Radoš, M., Sedmak, G., Benjak, V., Kostović-Srzentić, M., Vasung, L., Čuljat, M., Radoš, M., Hüppi, P., & Judaš, M. (2014). Perinatal and early postnatal reorganization of the subplate and related cellular compartments in the human cerebral wall as revealed by histological and MRI approaches. Brain Structure & Function, 219(1), 231–253. 10.1007/s00429-012-0496-0

Kretschmann, H. J., Tafesse, U., & Herrmann, A. (1982). Different volume changes of cerebral cortex and white matter during histological preparation. Microscopica Acta, 86(1), 13–24. https://pubmed.ncbi.nlm.nih.gov/7048029/

Künzle, H. (1978). Cortico-cortical efferents of primary motor and somatosensory regions of the cerebral cortex in Macaca fascicularis. Neuroscience, 3(1), 25–39. 10.1016/0306-4522(78)90151-3

Lee, S., Shin, H.-G., Kim, M., & Lee, J. (2023). Depth-wise profiles of iron and myelin in the cortex and white matter using χ-separation: A preliminary study. NeuroImage, 273(120058), 120058. 10.1016/j.neuroimage.2023.120058

Lewis, J. W., & Van Essen, D. C. (2000). Corticocortical connections of visual, sensorimotor, and multimodal processing areas in the parietal lobe of the macaque monkey. The Journal of Comparative Neurology, 428(1), 112–137. 10.1002/1096-9861(20001204)428:1%253C112::aid-cne8%253E3.0.co;2-9

Ludwig, E., & Klingler, J. (1956). Atlas cerebri humani. Little, Brown.

Luys, J. B. (1865). Recherches sur le système nerveux cérébro-spinal, sa structure, ses fonctions et ses maladies. Paris, J.B. Baillière et fils.

McKee, A. C., Cantu, R. C., Nowinski, C. J., Hedley-Whyte, E. T., Gavett, B. E., Budson, A. E., Santini, V. E., Lee, H.-S., Kubilus, C. A., & Stern, R. A. (2009). Chronic traumatic encephalopathy in athletes: progressive tauopathy after repetitive head injury. Journal of Neuropathology and Experimental Neurology, 68(7), 709–735. 10.1097/NEN.0b013e3181a9d503

Mesulam, M. M., & Mufson, E. J. (1984). Neural inputs into the nucleus basalis of the substantia innominata (Ch4) in the rhesus monkey. Brain: A Journal of Neurology, 107(1), 253–274. 10.1093/brain/107.1.253

Meynert, T. (1885). Psychiatry; A clinical treatise on diseases of the fore-brain based upon a study of its structure, functions, and nutrition. G P Putnam’s Sons.

Mohedano-Moriano, A., Martinez-Marcos, A., Muñoz, M., Arroyo-Jimenez, M. M., Marcos, P., Artacho-Pérula, E., Blaizot, X., & Insausti, R. (2005). Reciprocal connections between olfactory structures and the cortex of the rostral superior temporal sulcus in the Macaca fascicularis monkey. The European Journal of Neuroscience, 22(10), 2503–2518. 10.1111/j.1460-9568.2005.04443.x

Morecraft, R. J., Stilwell-Morecraft, K. S., Cipolloni, P. B., Ge, J., McNeal, D. W., & Pandya, D. N. (2012). Cytoarchitecture and cortical connections of the anterior cingulate and adjacent somatomotor fields in the rhesus monkey. Brain Research Bulletin, 87(4–5), 457–497. 10.1016/j.brainresbull.2011.12.005

Morecraft, Robert J., Ge, J., Stilwell-Morecraft, K. S., Rotella, D. L., Pizzimenti, M. A., & Darling, W. G. (2019). Terminal organization of the corticospinal projection from the lateral premotor cortex to the cervical enlargement (C5-T1) in rhesus monkey. The Journal of Comparative Neurology, 527(16), 2761–2789. 10.1002/cne.24706

Morris, R., Pandya, D. N., & Petrides, M. (1999). Fiber system linking the mid-dorsolateral frontal cortex with the retrosplenial/presubicular region in the rhesus monkey. The Journal of Comparative Neurology, 407(2), 183–192. 10.1002/(sici)1096-9861(19990503)407:2%253C183::aid-cne3%253E3.0.co;2-n

Morris, R., Petrides, M., & Pandya, D. N. (1999). Architecture and connections of retrosplenial area 30 in the rhesus monkey (Macaca mulatta). The European Journal of Neuroscience, 11(7), 2506–2518. 10.1046/j.1460-9568.1999.00672.x

Mortazavi, F., Romano, S. E., Rosene, D. L., & Rockland, K. S. (2017). A survey of white matter neurons at the gyral crowns and sulcal depths in the rhesus monkey. Frontiers in Neuroanatomy, 11, 69. 10.3389/fnana.2017.00069

Mufson, E. J., Mesulam, M. M., & Pandya, D. N. (1981). Insular interconnections with the amygdala in the rhesus monkey. Neuroscience, 6(7), 1231–1248. 10.1016/0306-4522(81)90184-6

Mufson, E. J., & Pandya, D. N. (1984). Some observations on the course and composition of the cingulum bundle in the rhesus monkey. The Journal of Comparative Neurology, 225(1), 31–43. 10.1002/cne.902250105

Muñoz-López, M., Insausti, R., Mohedano-Moriano, A., Mishkin, M., & Saunders, R. C. (2015). Anatomical pathways for auditory memory II: information from rostral superior temporal gyrus to dorsolateral temporal pole and medial temporal cortex. Frontiers in Neuroscience, 9, 158. 10.3389/fnins.2015.00158

Nakamura, H., Kuroda, T., Wakita, M., Kusunoki, M., Kato, A., Mikami, A., Sakata, H., & Itoh, K. (2001). From three-dimensional space vision to prehensile hand movements: the lateral intraparietal area links the area V3A and the anterior intraparietal area in macaques. The Journal of Neuroscience: The Official Journal of the Society for Neuroscience, 21(20), 8174–8187. 10.1523/jneurosci.21-20-08174.2001

Nie, X., Ruan, J., Otaduy, M. C. G., Grinberg, L. T., Ringman, J., & Shi, Y. (2024). Surface-based probabilistic fiber tracking in superficial white matter. IEEE Transactions on Medical Imaging, 43(3), 1113–1124. 10.1109/TMI.2023.3329451

Niu, M., Impieri, D., Rapan, L., Funck, T., Palomero-Gallagher, N., & Zilles, K. (2020). Receptor-driven, multimodal mapping of cortical areas in the macaque monkey intraparietal sulcus. eLife, 9, e55979. 10.7554/eLife.55979

Oishi, K., Huang, H., Yoshioka, T., Ying, S. H., Zee, D. S., Zilles, K., Amunts, K., Woods, R., Toga, A. W., Pike, G. B., Rosa-Neto, P., Evans, A. C., van Zijl, P. C. M., Mazziotta, J. C., & Mori, S. (2011). Superficially located white matter structures commonly seen in the human and the macaque brain with diffusion tensor imaging. Brain Connectivity, 1(1), 37–47. 10.1089/brain.2011.0005

Oishi, K., Zilles, K., Amunts, K., Faria, A., Jiang, H., Li, X., Akhter, K., Hua, K., Woods, R., Toga, A. W., Pike, G. B., Rosa-Neto, P., Evans, A., Zhang, J., Huang, H., Miller, M. I., van Zijl, P. C. M., Mazziotta, J., & Mori, S. (2008). Human brain white matter atlas: identification and assignment of common anatomical structures in superficial white matter. NeuroImage, 43(3), 447–457. 10.1016/j.neuroimage.2008.07.009

Pandya, D. N., & Rosene, D. L. (1993). Laminar termination patterns of thalamic, callosal, and association afferents in the primary auditory area of the rhesus monkey. Experimental Neurology, 119(2), 220–234. 10.1006/exnr.1993.1024

Pandya, D. N., Rosene, D. L., & Doolittle, A. M. (1994). Corticothalamic connections of auditory-related areas of the temporal lobe in the rhesus monkey. The Journal of Comparative Neurology, 345(3), 447–471. 10.1002/cne.903450311

Pandya, D. N., & Seltzer, B. (1982). Intrinsic connections and architectonics of posterior parietal cortex in the rhesus monkey. The Journal of Comparative Neurology, 204(2), 196–210. 10.1002/cne.902040208

Pandya, D. N., & Rosene, D. L. (1985). Some observations on trajectories and topography of commissural fibers. In Epilepsy and the Corpus Callosum (pp. 21–39). Springer US. 10.1007/978-1-4613-2419-5_2

Penfield, W., & Rasmussen, T. (1950). The cerebral cortex of man; a clinical study of localization of function. Macmillan.

Penfield, W., & Boldrey, E. (1937). Somatic motor and sensory representation in the cerebral cortex of man as studied by electrical stimulation. Brain: A Journal of Neurology, 60(4), 389–443. 10.1093/brain/60.4.389

Petrides, M., & Pandya, D. N. (1984). Projections to the frontal cortex from the posterior parietal region in the rhesus monkey. The Journal of Comparative Neurology, 228(1), 105–116. 10.1002/cne.902280110

Petrides, M., & Pandya, D. N. (2006). Efferent association pathways originating in the caudal prefrontal cortex in the macaque monkey. The Journal of Comparative Neurology, 498(2), 227–251. 10.1002/cne.21048

Petrides, M., & Pandya, D. N. (1988). Association fiber pathways to the frontal cortex from the superior temporal region in the rhesus monkey. The Journal of Comparative Neurology, 273(1), 52–66. 10.1002/cne.902730106

Petrides, M., & Pandya, D. N. (2002). Association pathways of the prefrontal cortex and functional observations. In D. T. Stuss & R. T. Knight (Eds.), Principles of Frontal Lobe Function (pp. 31–50). Oxford University Press. 10.1093/acprof:oso/9780195134971.003.0003

Petrides, M., & Pandya, D. N. (2007). Efferent association pathways from the rostral prefrontal cortex in the macaque monkey. The Journal of Neuroscience: The Official Journal of the Society for Neuroscience, 27(43), 11573–11586. 10.1523/JNEUROSCI.2419-07.2007

Phillips, O. R., Joshi, S. H., Piras, F., Orfei, M. D., Iorio, M., Narr, K. L., Shattuck, D. W., Caltagirone, C., Spalletta, G., & Di Paola, M. (2016a). The superficial white matter in Alzheimer’s disease: Superficial White Matter in Alzheimer’s Disease. Human Brain Mapping, 37(4), 1321–1334. 10.1002/hbm.23105

Phillips, O. R., Joshi, S. H., Squitieri, F., Sanchez-Castaneda, C., Narr, K., Shattuck, D. W., Caltagirone, C., Sabatini, U., & Di Paola, M. (2016b). Major superficial white matter abnormalities in Huntington’s disease. Frontiers in Neuroscience, 10, 197. 10.3389/fnins.2016.00197

Pron, A., Deruelle, C., & Coulon, O. (2021). U-shape short-range extrinsic connectivity organisation around the human central sulcus. Brain Structure & Function, 226(1), 179–193. 10.1007/s00429-020-02177-5

Pucak, M. L., Levitt, J. B., Lund, J. S., & Lewis, D. A. (1996). Patterns of intrinsic and associational circuitry in monkey prefrontal cortex. The Journal of Comparative Neurology, 376(4), 614–630. 10.1002/(SICI)1096-9861(19961223)376:4%253C614::AID-CNE9%253E3.0.CO;2-4

Ramón y Cajal, S. (1995). Histology of the Nervous System of Man and Vertebrates (1909) (S. L. W. Swanson N, Trans.). Oxford University Press. https://play.google.com/store/books/details?id=VLlqAAAAMAAJ

Reveley, C., Seth, A. K., Pierpaoli, C., Silva, A. C., Yu, D., Saunders, R. C., Leopold, D. A., & Ye, F. Q. (2015). Superficial white matter fiber systems impede detection of long-range cortical connections in diffusion MR tractography. Proceedings of the National Academy of Sciences of the United States of America, 112(21), E2820–2828. 10.1073/pnas.1418198112

Rilling, J. K., & Insel, T. R. (1999). The primate neocortex in comparative perspective using magnetic resonance imaging. Journal of Human Evolution, 37(2), 191–223. 10.1006/jhev.1999.0313

Rockland, K. S., & Knutson, T. (2001). Axon collaterals of Meynert cells diverge over large portions of area V1 in the macaque monkey. The Journal of Comparative Neurology, 441(2), 134–147. 10.1002/cne.1402

Rockland, K. S., & Pandya, D. N. (1981). Cortical connections of the occipital lobe in the rhesus monkey: interconnections between areas 17, 18, 19 and the superior temporal sulcus. Brain Research, 212(2), 249–270. 10.1016/0006-8993(81)90461-3

Rockland, K. S. (2020). What we can learn from the complex architecture of single axons. Brain Structure & Function, 225(4), 1327–1347. 10.1007/s00429-019-02023-3

Rockland, K. S., & Drash, G. W. (1996). Collateralized divergent feedback connections that target multiple cortical areas. The Journal of Comparative Neurology, 373(4), 529–548. 10.1002/(SICI)1096-9861(19960930)373:4%253C529::AID-CNE5%253E3.0.CO;2-3

Rockland, K. S., & Rushmore, R. J. (2025). Cortical white matter: no longer a silent partner. Frontiers in Neuroanatomy, 19, 1726067. 10.3389/fnana.2025.1726067

Rojkova, K., Volle, E., Urbanski, M., Humbert, F., Dell’Acqua, F., & Thiebaut de Schotten, M. (2016). Atlasing the frontal lobe connections and their variability due to age and education: a spherical deconvolution tractography study. Brain Structure & Function, 221(3), 1751–1766. 10.1007/s00429-015-1001-3

Román, C., Guevara, M., Valenzuela, R., Figueroa, M., Houenou, J., Duclap, D., Poupon, C., Mangin, J.-F., & Guevara, P. (2017). Clustering of whole-brain white matter short association bundles using HARDI data. Frontiers in Neuroinformatics, 11, 73. 10.3389/fninf.2017.00073

Rozzi, S., Calzavara, R., Belmalih, A., Borra, E., Gregoriou, G. G., Matelli, M., & Luppino, G. (2006). Cortical connections of the inferior parietal cortical convexity of the macaque monkey. Cerebral Cortex (New York, N.Y.: 1991), 16(10), 1389–1417. 10.1093/cercor/bhj076

Sachs, H. (1892). Das Hemisphärenmark des menschlichen Grosshirns. Georg Thieme Verlag KG.

Sarubbo, S., De Benedictis, A., Maldonado, I. L., Basso, G., & Duffau, H. (2013). Frontal terminations for the inferior fronto-occipital fascicle: anatomical dissection, DTI study and functional considerations on a multi-component bundle. Brain Structure & Function, 218(1), 21–37. 10.1007/s00429-011-0372-3

Saunders, R. C., & Rosene, D. L. (1988). A comparison of the efferents of the amygdala and the hippocampal formation in the rhesus monkey: I. Convergence in the entorhinal, prorhinal, and perirhinal cortices: ENTORHINAL AFFERENTS FROM AMYGDALA AND HIPPOCAMPUS. The Journal of Comparative Neurology, 271(2), 153–184. 10.1002/cne.902710202

Schilling, K. G., Archer, D., Yeh, F.-C., Rheault, F., Cai, L. Y., Shafer, A., Resnick, S. M., Hohman, T., Jefferson, A., Anderson, A. W., Kang, H., & Landman, B. A. (2023a). Short superficial white matter and aging: a longitudinal multi-site study of 1293 subjects and 2711 sessions. Aging Brain, 3, 100067. 10.1016/j.nbas.2023.100067

Schilling, K. G., Archer, D., Rheault, F., Lyu, I., Huo, Y., Cai, L. Y., Bunge, S. A., Weiner, K. S., Gore, J. C., Anderson, A. W., & Landman, B. A. (2023b). Superficial white matter across development, young adulthood, and aging: volume, thickness, and relationship with cortical features. Brain Structure & Function, 228(3–4), 1019–1031. 10.1007/s00429-023-02642-x

Schilling, K., Gao, Y., Janve, V., Stepniewska, I., Landman, B. A., & Anderson, A. W. (2017). Can increased spatial resolution solve the crossing fiber problem for diffusion MRI? NMR in Biomedicine, 30(12), e3787. 10.1002/nbm.3787

Schilling, K., Gao, Y., Janve, V., Stepniewska, I., Landman, B. A., & Anderson, A. W. (2018). Confirmation of a gyral bias in diffusion MRI fiber tractography. Human Brain Mapping, 39(3), 1449–1466. 10.1002/hbm.23936

Schilling, K., Zhang, F., Román, C., O’Donnell, L. J., & Guevara, P. (2025). Short association fiber tractography: key insights and surprising facts. Brain Structure & Function, 230(6), 97. 10.1007/s00429-025-02966-w

Schindelin, J., Arganda-Carreras, I., Frise, E., Kaynig, V., Longair, M., Pietzsch, T., Preibisch, S., Rueden, C., Saalfeld, S., Schmid, B., Tinevez, J.-Y., White, D. J., Hartenstein, V., Eliceiri, K., Tomancak, P., & Cardona, A. (2012). Fiji: an open-source platform for biological-image analysis. Nature Methods, 9(7), 676–682. 10.1038/nmeth.2019

Schmahmann, J. D., & Pandya, D. N. (1997). The cerebrocerebellar system. International Review of Neurobiology, 41, 31–60. 10.1016/s0074-7742(08)60346-3

Schmahmann, Jeremy D., & Pandya, D. (2006). Fiber pathways of the brain. Oxford University Press. 10.1093/acprof:oso/9780195104233.001.0001

Schmahmann, Jeremy D., Pandya, D. N., Wang, R., Dai, G., D’Arceuil, H. E., de Crespigny, A. J., & Wedeen, V. J. (2007). Association fibre pathways of the brain: parallel observations from diffusion spectrum imaging and autoradiography. Brain: A Journal of Neurology, 130(Pt 3), 630–653. 10.1093/brain/awl359

Schmahmann, Jeremy D., Rosene, D. L., & Pandya, D. N. (2004). Motor projections to the basis pontis in rhesus monkey. The Journal of Comparative Neurology, 478(3), 248–268. 10.1002/cne.20286

Schüz, A., & Braitenberg, V. (2002). The Human Cortical White Matter: Quantitative Aspects of Cortico-Cortical Long-Range Connectivity. In A. Schüz & R. Miller (Eds.), Cortical Areas: Unity and Diversity (pp. 377–385). Taylor & Francis.

Sedmak, G., & Judaš, M. (2019). The total number of white matter interstitial neurons in the human brain. Journal of Anatomy, 235(3), 626–636. 10.1111/joa.13018

Sedmak, G., & Judaš, M. (2021). White matter interstitial neurons in the adult human brain: 3% of cortical neurons in quest for recognition. Cells (Basel, Switzerland), 10(1), 190. 10.3390/cells10010190

Seltzer, B., Cola, M. G., Gutierrez, C., Massee, M., Weldon, C., & Cusick, C. G. (1996). Overlapping and nonoverlapping cortical projections to cortex of the superior temporal sulcus in the rhesus monkey: double anterograde tracer studies. The Journal of Comparative Neurology, 370(2), 173–190. 10.1002/(SICI)1096-9861(19960624)370:2%253C173::AID-CNE4%253E3.0.CO;2-%2523

Seltzer, B., & Pandya, D. N. (1980). Converging visual and somatic sensory cortical input to the intraparietal sulcus of the rhesus monkey. Brain Research, 192(2), 339–351. 10.1016/0006-8993(80)90888-4

Seltzer, B., & Pandya, D. N. (1989). Intrinsic connections and architectonics of the superior temporal sulcus in the rhesus monkey. The Journal of Comparative Neurology, 290(4), 451–471. 10.1002/cne.902900402

Shah, A., Jhawar, S., Goel, A., & Goel, A. (2026). Anatomical issues related to the transcortical versus the trans-sulcal approach to intra-axial brain tumors: A fiber dissection study. Journal of Clinical Neuroscience: Official Journal of the Neurosurgical Society of Australasia, 148(111969), 111969. 10.1016/j.jocn.2026.111969

Shah, A., Jhawar, S. S., & Goel, A. (2025). The connections of the short arcuate fibers of the frontal lobe: An anatomical study. Neurology India, 73(1), 70–76. 10.4103/neurol-india.Neurol-India-D-23-00227

Shinohara, H., Liu, X., Nakajima, R., Kinoshita, M., Ozaki, N., Hori, O., & Nakada, M. (2020). Pyramid-shape crossings and intercrossing fibers are key elements for construction of the neural network in the superficial white matter of the human cerebrum. Cerebral Cortex (New York, N.Y.: 1991), 30(10), 5218–5228. 10.1093/cercor/bhaa080

Siegel, J. S., Shulman, G. L., & Corbetta, M. (2022). Mapping correlated neurological deficits after stroke to distributed brain networks. Brain Structure & Function, 227(9), 3173–3187. 10.1007/s00429-022-02525-7

Stefanacci, L., & Amaral, D. G. (2002). Some observations on cortical inputs to the macaque monkey amygdala: an anterograde tracing study. The Journal of Comparative Neurology, 451(4), 301–323. 10.1002/cne.10339

Tian, Q., Ngamsombat, C., Lee, H.-H., Berger, D. R., Wu, Y., Fan, Q., Bilgic, B., Li, Z., Novikov, D. S., Fieremans, E., Rosen, B. R., Lichtman, J. W., & Huang, S. Y. (2025). Quantifying axonal features of human superficial white matter from three-dimensional multibeam serial electron microscopy data assisted by deep learning. NeuroImage, 313, 121212. 10.1016/j.neuroimage.2025.121212

Van Dyken, P. C., Khan, A. R., & Palaniyappan, L. (2024). Imaging of the superficial white matter in health and disease. Imaging Neuroscience, 2, 1–35. 10.1162/imag_a_00221

Vogt, O. (1910). Die myeloarchitektonische Felderung des menschlichen Stirnhirns. Journal of Psychology and Neurology, 15, 221–238.

Vogt, O. (1911). La nouvelle division myeloarchitecturale de l’ecorce cerebrale et ses rapports avec la physiologie et la psychologie. Journal of Psychology and Neurology, 17, 1–15.

Von Monakow, C. (1905). Gehirnpathologie. Alfred Hölder.

Wang, S., Zhang, F., Zeng, Q., Hong, H., Zhang, Y., Xie, L., Lin, M., Jiaerken, Y., Yu, X., Zhang, R., Luo, X., Li, K., Xu, X., Hassanzadeh-Behbahani, S., Lin, B., Rushmore, J., Wang, C., Rathi, Y., Makris, N.,…Alzheimer’s Disease Neuroimaging Initiative. (2025). Association of superficial white matter microstructure with cortical pathology deposition across early stages of the AD continuum. Neurology, 105(2), e213666. 10.1212/WNL.0000000000213666

Wu, M., Kumar, A., & Yang, S. (2016). Development and aging of superficial white matter myelin from young adulthood to old age: Mapping by vertex-based surface statistics (VBSS): Wu et al. Human Brain Mapping, 37(5), 1759–1769. 10.1002/hbm.23134

Wu, M., Lu, L. H., Lowes, A., Yang, S., Passarotti, A. M., Zhou, X. J., & Pavuluri, M. N. (2014). Development of superficial white matter and its structural interplay with cortical gray matter in children and adolescents: Development of SWM in Children and Adolescents. Human Brain Mapping, 35(6), 2806–2816. 10.1002/hbm.22368

Xu, F., Shen, Y., Ding, L., Yang, C.-Y., Tan, H., Wang, H., Zhu, Q., Xu, R., Wu, F., Xiao, Y., Xu, C., Li, Q., Su, P., Zhang, L. I., Dong, H.-W., Desimone, R., Xu, F., Hu, X., Lau, P.-M., & Bi, G.-Q. (2021). High-throughput mapping of a whole rhesus monkey brain at micrometer resolution. Nature Biotechnology, 39(12), 1521–1528. 10.1038/s41587-021-00986-5

Xue, T., Zhang, F., Zhang, C., Chen, Y., Song, Y., Golby, A. J., Makris, N., Rathi, Y., Cai, W., & O’Donnell, L. J. (2023). Superficial white matter analysis: An efficient point-cloud-based deep learning framework with supervised contrastive learning for consistent tractography parcellation across populations and dMRI acquisitions. Medical Image Analysis, 85(102759), 102759. 10.1016/j.media.2023.102759

Yamashita, A., & Arikuni, T. (2001). Axon trajectories in local circuits of the primary motor cortex in the macaque monkey (Macaca fuscata). Neuroscience Research, 39(2), 233–245. 10.1016/s0168-0102(00)00220-0

Ye, A. Q., Zhan, L., Conrin, S., GadElKarim, J., Zhang, A., Yang, S., Feusner, J. D., Kumar, A., Ajilore, O., & Leow, A. (2015). Measuring embeddedness: Hierarchical scale-dependent information exchange efficiency of the human brain connectome: Hierarchical Efficiency of the Brain. Human Brain Mapping, 36(9), 3653–3665. 10.1002/hbm.22869

Yendiki, A., Aggarwal, M., Axer, M., Howard, A. F. D., van Walsum, A.-M. van C., & Haber, S. N. (2022). Post mortem mapping of connectional anatomy for the validation of diffusion MRI. NeuroImage, 256(119146), 119146. 10.1016/j.neuroimage.2022.119146

Yeterian, E. H., & Pandya, D. N. (1985). Corticothalamic connections of the posterior parietal cortex in the rhesus monkey. The Journal of Comparative Neurology, 237(3), 408–426. 10.1002/cne.902370309

Yeterian, E. H., & Pandya, D. N. (1997). Corticothalamic connections of extrastriate visual areas in rhesus monkeys. The Journal of Comparative Neurology, 378(4), 562–585. 10.1002/(SICI)1096-9861(19970224)378:4%253C562::AID-CNE10%253E3.0.CO;2-L

Yeterian, E. H., & Pandya, D. N. (2010). Fiber pathways and cortical connections of preoccipital areas in rhesus monkeys. The Journal of Comparative Neurology, 518(18), 3725–3751. 10.1002/cne.22420

Yoshino, M., Saito, K., Kawasaki, K., Horiike, T., Shinmyo, Y., & Kawasaki, H. (2020). The origin and development of subcortical U-fibers in gyrencephalic ferrets. Molecular Brain, 13(1), 37. 10.1186/s13041-020-00575-8

Zemmoura, I., Blanchard, E., Raynal, P.-I., Rousselot-Denis, C., Destrieux, C., & Velut, S. (2016). How Klingler’s dissection permits exploration of brain structural connectivity? An electron microscopy study of human white matter. Brain Structure & Function, 221(5), 2477–2486. 10.1007/s00429-015-1050-7

Zemmoura, I., Serres, B., Andersson, F., Barantin, L., Tauber, C., Filipiak, I., Cottier, J.-P., Venturini, G., & Destrieux, C. (2014). FIBRASCAN: a novel method for 3D white matter tract reconstruction in MR space from cadaveric dissection. NeuroImage, 103, 106–118. 10.1016/j.neuroimage.2014.09.016

Zhang, F., Chen, Y., Ning, L., Rushmore, J., Liu, Q., Du, M., Hassanzadeh-Behbahani, S., Legarreta, J. H., Yeterian, E., Makris, N., Rathi, Y., & O’Donnell, L. J. (2024). Assessment of the depiction of superficial white matter using ultra-high-resolution diffusion MRI. Human Brain Mapping, 45(14), e70041. 10.1002/hbm.70041

Zhang, T., Chen, H., Guo, L., Li, K., Li, L., Zhang, S., Shen, D., Hu, X., & Liu, T. (2014). Characterization of U-shape streamline fibers: Methods and applications. Medical Image Analysis, 18(5), 795–807. 10.1016/j.media.2014.04.005

Zikopoulos, B., & Barbas, H. (2007). Parallel driving and modulatory pathways link the prefrontal cortex and thalamus. PloS One, 2(9), e848. 10.1371/journal.pone.0000848

Zikopoulos, B., & Barbas, H. (2010). Changes in prefrontal axons may disrupt the network in autism. The Journal of Neuroscience: The Official Journal of the Society for Neuroscience, 30(44), 14595–14609. 10.1523/JNEUROSCI.2257-10.2010

Zikopoulos, B., & Barbas, H. (2013). Altered neural connectivity in excitatory and inhibitory cortical circuits in autism. Frontiers in Human Neuroscience, 7, 609. 10.3389/fnhum.2013.00609

Zikopoulos, B., García-Cabezas, M. Á., & Barbas, H. (2018). Parallel trends in cortical gray and white matter architecture and connections in primates allow fine study of pathways in humans and reveal network disruptions in autism. PLoS Biology, 16(2), e2004559. 10.1371/journal.pbio.2004559

Zilles, K., & Palomero-Gallagher, N. (2017). Multiple transmitter receptors in regions and layers of the human cerebral cortex. Frontiers in Neuroanatomy, 11, 78. 10.3389/fnana.2017.00078

Zouridakis, A., Ayala, I., Minogue, G., Kawles, A., Keszycki, R., Macomber, A., Bigio, E. H., Geula, C., Mesulam, M.-M., & Gefen, T. (2023). Shades of gray in human white matter. The Journal of Comparative Neurology, 531(18), 2109–2120. 10.1002/cne.25512

