## Supplementary Case Atlas for "Connectional neuroanatomy of U-fibers in the rhesus monkey brain"

### Table of Contents


1. Overview ... 2

2. Prefrontal Cortex ... 3

3. Motor Cortex ... 8

4. Temporal Lobe ... 13

5. Parietal Lobe ... 15

   5.1 Rostral Intraparietal Sulcus ... 15

   5.2 Mid Intraparietal Sulcus ... 20

6. Preoccipital Region ... 27

7. Abbreviations ... 31

8. References ... 31

### 1. Overview

This Supplementary Case Atlas provides detailed descriptions of the 14 representative cases presented throughout the paper, illustrating the core organizational features of short association fiber bundles and the superficial white matter observed across the cases analyzed. Cases were selected from the Pandya-Rosene Neuroanatomical Archive in which radiolabeled tritiated amino acids were injected on gyri adjacent to four major sulci (the principal, central, intraparietal, and superior temporal sulci) each of which has clear anatomical correspondences to sulci in the human cerebral cortex. Following an initial quality assessment (see Methods), 28 cases were analyzed.

For each case, the organization of long-range efferent fiber components is briefly described. These components include: 1) cord fibers dividing into commissural fibers that travel to the opposite hemisphere through either the corpus callosum or anterior commissure, and a subcortical bundle that divides further into thalamic and pontine components; 2) long cortico-cortical association fibers, identified by their characteristic stippled appearance of transverse fiber profiles and their location in the deep white matter within named fiber pathways; and 3) cortico-striatal projections coursing through the subcallosal fasciculus of Muratoff, or Muratoff's bundle, and/or the external capsule (Schmahmann & Pandya, 2006). The presence of these well-characterized efferent fiber components confirmed that tracer was deposited across all cortical layers in each case. The trajectories and terminations of these long-range systems in these cases, or in cases with similar injections, have been described in detail elsewhere, including their thalamic connectivity (e.g., Yeterian & Pandya, 1985, 1991, 1997; Siwek & Pandya, 1991; Schmahmann & Pandya, 2006), pontine projections (e.g., Schmahmann & Pandya, 1989, 1991, 1992, 1993, 1997, 2006; Schmahmann et al., 2004), cortico-striatal projections (e.g., Yeterian & Pandya, 1991, 1993, 1995, 1998; Schmahmann & Pandya, 2006), and long association fiber pathways, including the superior longitudinal fasciculus (SLF), cingulum bundle, uncinate fasciculus, arcuate fasciculus, middle longitudinal fasciculus (MdLF), inferior longitudinal fasciculus (ILF), and fronto-occipital fasciculus (FOF) (Mufson & Pandya, 1984; Petrides & Pandya, 1984, 1988, 2006, 2009; Seltzer & Pandya, 1989; Schmahmann & Pandya, 2006; Yeterian & Pandya, 2010). The primary focus of each case description is therefore on the local short association fiber bundles, specifically their trajectories, termination patterns, and organizational relationships within the superficial white matter.

### 2. Prefrontal Cortex

In the prefrontal cortex, the sulcal white matter was analyzed along the rostrocaudal extent of the principal sulcus in two cases (BMFG, BMFF). The principal sulcus is the primary sulcus of the dorsolateral prefrontal cortex. It begins proximal the frontal pole, extends caudally to divide the dorsolateral prefrontal cortex into dorsal and ventral subdivisions, and ends rostral to the concavity of the arcuate sulcus.

#### *2.1 Injection Sites and Long-Range Fiber Systems*

Case BMFG had an injection on the dorsal gyrus of the principal sulcus, centered on cortical area 9/46d and encroaching on area 8Ad (Supplementary Figures 1A-B, 2A-B; red asterisks), whereas case BMFF had an injection on the ventral gyrus centered on cortical area 9/46v (Supplementary Figure 1C-D, red asterisks). In both cases, the injection produced a cord of densely labeled fibers (Supplementary Figure 1B, D; solid black or white lines) that traveled medially through the deep white matter before separating into two principal components. The more dorsal bundle continued medially, entering the corpus callosum to reach the opposite hemisphere, whereas the more ventral fiber bundle traveled to the anterior limb of the internal capsule before further separating into fibers projecting to the brainstem and thalamus. Cortico-striatal fibers were also observed in two distinct groups: medially directed fibers that entered Muratoff's bundle and coursed caudally to terminate in the caudate nucleus and ventrally directed fibers that entered the external and/or extreme capsule to reach the putamen and claustrum. Both cases also gave rise to separate long association fiber systems (e.g., SLF I-III, cingulum bundle, uncinate fasciculus, FOF). These systems were identifiable based on their deep position in the white matter and their characteristic stippled appearance of transverse fiber profiles.

#### *2.2 Short Association Fibers along the Principal Sulcus*

Label corresponding to local short association fiber bundles was evident beneath the principal sulcus in both cases, forming prominent U-shaped bundles (Supplementary Figure 1B, D; solid red lines). In case BMFG, the injection was centered on the dorsal gyrus (Supplementary Figure 1A-B, red asterisks) and produced a densely labeled U-shaped bundle beneath the principal sulcus (Supplementary Figure 1B, solid red line), with columnar terminations in cortical area 9/46v on the adjacent gyrus and ventral bank of the sulcus (red arrows). Further caudally, additional terminations were observed on the opposing gyrus in cortical area 8Av and along the ventral sulcal bank in cortical area 9/46v (Supplementary Figure 2B, red arrowheads). The bundle consisted of densely labeled longitudinal fiber profiles that coursed from the injection site within the white matter directly beneath the gray matter-white matter (GM-WM) border along the dorsal bank and fundus. This pattern indicates that fibers traveled roughly within the plane of section and orthogonal to the long axis of the sulcus (see Figure 14B). After curving beneath the sulcus, fiber profiles along the ventral bank transitioned to a mix of longitudinal and oblique profiles, indicating that the bundle fanned out in the rostrocaudal direction before terminating. A similar organization was observed in case BMFF, with the injection centered on the ventral gyrus (Supplementary Figure 1C-D, red asterisks). Labeled fibers formed a U-shaped bundle (Supplementary Figure 1D, solid red line), emerging from the injection as tightly arrayed in-plane profiles before becoming relatively dispersed after curving around the sulcal fundus and terminating in a columnar manner in cortical area 9/46d on the adjacent gyrus (red arrow). In both cases, label was also observed in cortical layer 6, continuous with the underlying white matter label (Supplementary Figure 1B, D; white arrowheads). At the caudal extent of the injection and U-shaped bundle in case BMFG, labeling within cortical layer 6 (Supplementary Figure 2B, white arrowheads) occupied a larger portion of the layer as it continued around the sulcus, suggesting a progressive shift of label from the white matter into the lower cortical layers.

In both cases, the U-shaped bundles interconnected opposing gyri beneath the principal sulcus and were classified as U-fibers. Columnar terminations on the adjacent gyrus were located at approximately the same rostrocaudal level as the injection in both cases and labeled fiber profiles maintained a relatively consistent orientation orthogonal to the long axis of the sulcus throughout their trajectory, supporting a symmetric classification (see Results: U-Fiber Trajectories do not Always Reflect Sulcal Geometry). Both U-fibers occupied the white matter directly beneath the GM-WM border (Supplementary Figures 1B, D, 2B; solid green line), with depth measurements confirming that each bundle coursed within the 200-300 μm superficial white matter band (Supplementary Figure 1B, D; dashed yellow line): in BMFG, the superficial white matter band measured a mean of 240 μm (SD 18.3 μm; range 220-263 μm), and in BMFF, it measured a mean of 239 μm (SD 24.0 μm; range 207-264 μm). Together, these measurements indicate that both bundles occupied the white matter at equivalent depths and were arranged symmetrically around the sulcus, suggesting that reciprocal fibers from the two gyri likely intermingle within this superficial band.

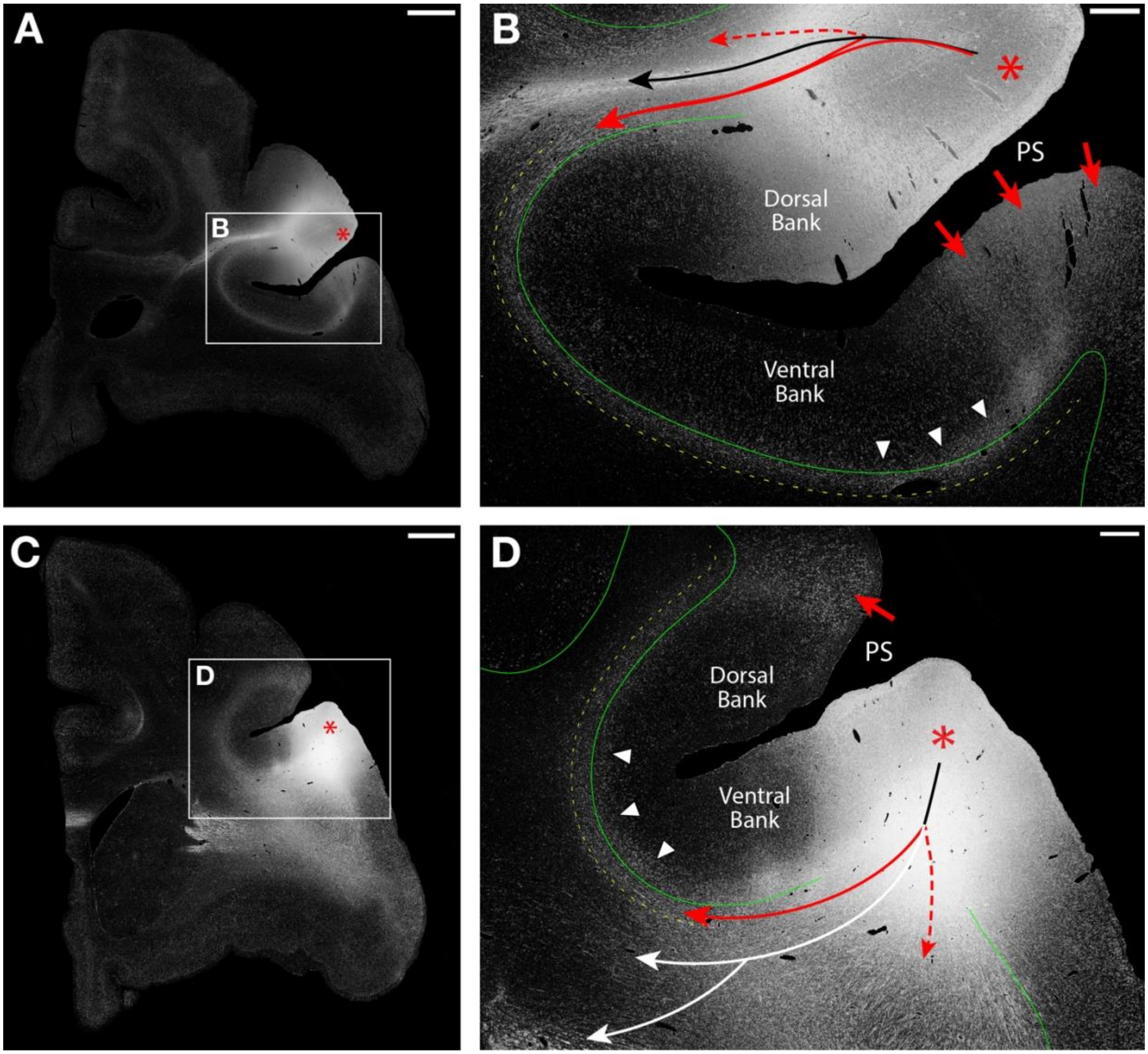


**Supplementary Figure 1:** Autoradiographs of short association fibers in the coronal plane from two macaque monkey prefrontal cases processed for autoradiography and imaged under darkfield illumination. A. Low-magnification darkfield image following an injection in dorsal area 9/46 (red asterisk). B. Higher-magnification view of boxed region in A highlighting the different labeled fiber bundles: a U-shaped bundle (solid red line) coursing around the principal sulcus (PS), cord fibers within the gyral core (solid black line), and an additional superficial bundle coursing around the superior limb of the arcuate sulcus (dashed red line). The U-shaped bundle around the PS travels within the white matter immediately deep to the gray matter-white matter (GM-WM) border (solid green line). Labeling extends across the GM-WM border into the cortex (white arrowheads) before forming columnar terminations along the ventral bank of the PS and gyrus (red arrows). Throughout its trajectory, this U-fiber remains within the superficial white matter band (dashed yellow line). C. Low-magnification view of an injection in ventral area 9/46 (red asterisk). D. Higher-magnification view of the boxed region in C showing a U-shaped bundle (solid red line) coursing around the PS, cord fibers (solid white lines), and a ventrally directed superficial fiber bundle that curve caudally around the inferior limb of the arcuate sulcus (dashed red line). The U-shaped fiber bundle around the PS has extensive labeling extending into the cortex (white arrowheads) as it courses in the white matter directly below the GM-WM border (solid green line) before terminating in a columnar manner on the adjacent gyrus (red arrow). This U-fiber remains restricted to the superficial white matter band (dashed yellow line) throughout its trajectory around PS. Panels A-B are from case BMFG; Panels C-D are from case BMFF. Scale bars: A, C = 2 mm; B, D = 500 μm.

*2.3 Short Association Fibers along the Arcuate Sulcus*

In both prefrontal cases, additional labeled fiber bundles were observed beneath the arcuate sulcus. In case BMFG, labeled fibers from the dorsal injection followed a U-shaped trajectory (Supplementary Figure 1B, dashed red line) beneath the superior limb of the arcuate sulcus. At this level, the sulcus was relatively shallow and oblique to the plane of section; therefore, fibers appeared less compact and more diffuse than those observed beneath the principal sulcus. Labeled fibers coursed directly beneath the GM-WM border as predominantly longitudinal profiles, traveling medially from the injection site along the ventral bank of the arcuate sulcus with an orientation roughly within the plane of section and orthogonal to the local long axis of the sulcus. Further caudally, the fiber bundle crossed the sulcal fundus (Supplementary Figure 2B, dashed red line) and ascended the dorsal bank. Fiber profiles then progressively shifted from oblique to transverse, indicating that the bundle fanned out before forming columnar terminations in cortical area 6 on the sulcal bank and gyral crown (Supplementary Figure 2B, red arrows), and further caudally in cortical area 6DR on the gyral crown. Because this U-shaped bundle interconnected adjacent gyri, it was classified as a U-fiber. These labeled fibers maintained an orientation orthogonal to the long axis of the sulcus and terminated on the adjacent gyrus at approximately the same rostrocaudal level as the injection, supporting a symmetric classification analogous to the principal sulcus U-fiber described above. Throughout its trajectory, the bundle maintained a position within the superficial white matter band directly beneath the GM-WM border. Although quantitative depth measurements were not obtained for this bundle, overlay of the 300 μm superficial white matter band boundary (Supplementary Figure 2B, dashed yellow line) confirmed that labeled fibers were largely confined to this band.

In case BMFF, labeled fibers from the ventral injection were observed beneath the inferior limb of the arcuate sulcus. Unlike the U-fibers described above, the trajectory of this bundle was difficult to evaluate with certainty because of the oblique orientation of the inferior limb relative to the coronal plane and the bundle's predominantly caudal trajectory. It was therefore not included in the primary analysis. Briefly, labeled fibers emanated from the injection site as predominantly longitudinal profiles (Supplementary Figure 1D, dashed red line), fanning out ventrally within the gyral white matter core rostral to the inferior limb before shifting to profiles indicating a rostrocaudal orientation. Further caudally, as the inferior limb appeared, a densely labeled band formed directly beneath the GM-WM border along the caudal sulcal bank, suggesting a U-shaped trajectory. These labeled fibers were associated with columnar terminations in ventral area 6,

ProM, and area 44 on the opposing gyrus, consistent with a U-fiber, and additional terminations along the sulcal banks and fundus in cortical areas 8Av and 45.

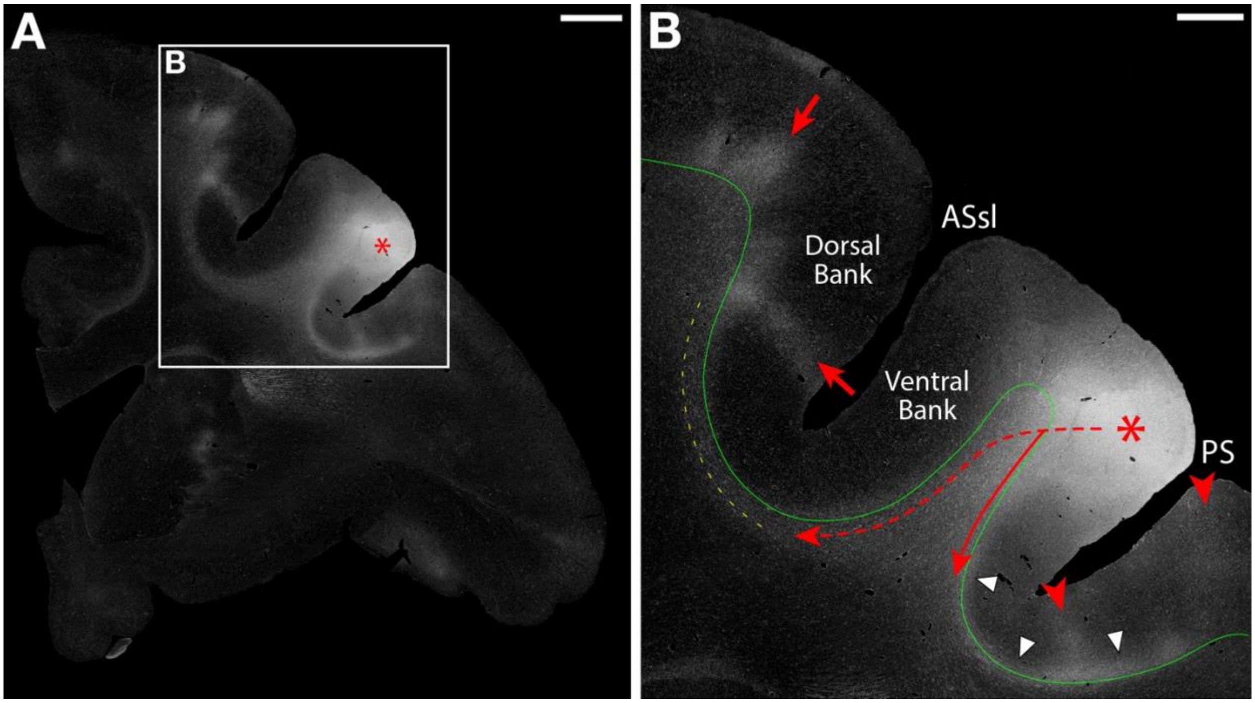


**Supplementary Figure 2:** Autoradiographs of short association fibers in the prefrontal cortex of the macaque monkey. Coronal section from one case processed for autoradiography and imaged under darkfield illumination. A. Low-magnification darkfield image following an injection in dorsal area 9/46 encroaching on cortical area 8Ad (red asterisk). B. Higher-magnification view of the boxed region in A showing two U-shaped bundles. The first (dashed red line) courses beneath the superior limb of the arcuate sulcus (ASsl), providing columnar terminations on the dorsal sulcal bank and gyral crown (red arrows). Therefore, this bundle was classified as a U-fiber. The second (solid red line) courses around the principal sulcus (PS), with columnar terminations on the sulcal bank and gyral crown (red arrowheads). Both bundles travel within the white matter directly beneath the gray matter-white matter (GM-WM) border (solid green line) and within the superficial white matter band (dashed yellow line). Label extending across the GM-WM border into the cortex along the PS is also present (white arrowheads). Case BMFG. Scale bars: A = 2 mm; B = 1 mm.

#### *2.4 Intragyral Fibers*

Intragyral fibers were observed in both cases as labeled transverse profiles, indicating a rostrocaudal trajectory within the white matter beneath the injected gyrus. In case BMFG, intragyral fibers terminated rostral to the injection in the dorsal portions of cortical areas 46 and 9, and caudally in cortical area 8Ad. An additional group of intragyral fibers coursed medially from the injection site, occupying a large portion of the superior frontal gyrus white matter as predominantly longitudinal profiles before forming columnar terminations in cortical area 8B on the medial surface of the hemisphere. Further caudally, these fibers were observed as more diffuse label with mixed fiber profiles, providing additional terminations in cortical area 8B. In case BMFF, intragyral fibers coursed rostrally from the injection to terminate in the ventral portions of cortical area 46, and caudally to terminate in cortical area 8Av.

#### *2.5 Summary*

The bundles identified beneath both the principal sulcus and the arcuate sulcus, the latter analyzed only in BMFG, matched the classical conception of U-fibers. That is, they were oriented orthogonally to the long axis of the sulcus, terminated on the adjacent gyrus at approximately the same rostrocaudal level as the injection, and traveled within the superficial white matter band directly beneath the GM-WM border. In both prefrontal cases, label was also observed in cortical layer 6, continuous with the underlying white matter label, underscoring the importance of precise GM-WM border delineation in distinguishing white matter fiber bundles from intracortical labeling.

### 3. Motor Cortex

The white matter was analyzed along the rostrocaudal extent of the central sulcus in two motor cortex cases (PGR-R, BMEX-L). The central sulcus, a key border between the frontal and parietal lobes, has been a common focus for studying U-fibers and local connections in macaque and human brains (e.g., Jones et al., 1978; Künzle, 1978; DeFelipe et al., 1986; Yamashita & Arikuni, 2001; Catani et al., 2012). It extends obliquely from dorsal to ventral on the lateral aspect of the hemisphere, separating the primary motor cortex of the precentral gyrus rostrally (cortical area 4) from the primary somatosensory cortex of the postcentral gyrus caudally (cortical areas 3, 1, 2).

#### *3.1 Injection Sites and Long-Range Fiber Systems*

Injections were placed in the precentral gyrus, centered on the hand representation of cortical area 4 in case PGR-R (Supplementary Figure 3A-B, red asterisks), confirmed by electrophysiological stimulation, and on the trunk representation of cortical area 4 in case BMEX-L (Supplementary Figure 3C-D, red asterisks). In both cases, the injection produced a thick cord of densely labeled fibers that descended medially in the deep white matter and separated into a commissural bundle (Supplementary Figure 3B, solid black line) directed to the opposite hemisphere via the corpus callosum and a subcortical bundle (solid white line) that descended ventrally through the corona radiata to enter the posterior limb of the internal capsule before separating into thalamic and brainstem projections. Cortico-striatal fibers were also observed descending ventrally from the injection site toward the external and extreme capsules to reach the putamen and claustrum in both cases. In case BMEX-L, a small number of fibers also entered Muratoff’s bundle to reach the caudate nucleus. Both cases gave rise to long association fibers (e.g., SLF I-II), identifiable by their deep white matter position and characteristic stippled appearance of transverse fiber profiles.

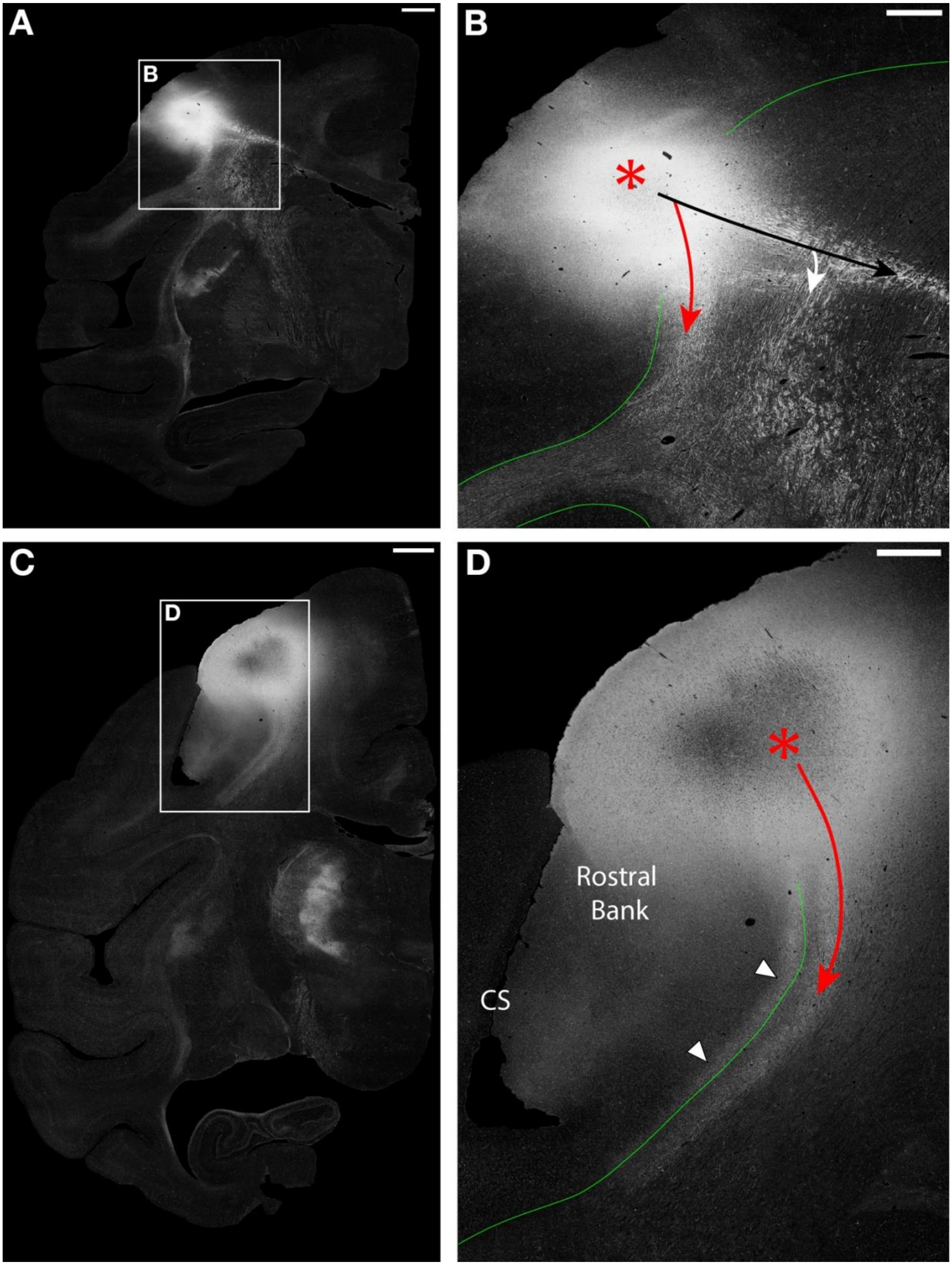
A
B
B
C
D
D
Rostral
Bank
CS

**Supplementary Figure 3**: Autoradiographs of short association fibers in the peri-Rolandic (motor/somatosensory) cortex of the macaque monkey. Coronal sections from two motor cortex cases processed for autoradiography and imaged under darkfield illumination. A. Low-magnification darkfield image following an injection in the hand representation of the primary motor cortex (M1) on the precentral gyrus (red asterisk). B. Higher-magnification view of the boxed region in A highlighting multiple bundles: a U-shaped bundle (solid red line) courses caudally around the central sulcus (CS) within the white matter directly beneath the gray matter-white matter (GM-WM) border (solid green line), alongside cord fibers divided into commissural (solid black line) and subcortical components (solid white line). C. Low-magnification darkfield image following an injection in the M1 trunk representation on the precentral gyrus (red asterisk), ~2 mm caudal to the main injection foci. D. Higher-magnification view of the boxed region in C showing two distinct intensity bands beneath the CS: a superficial band traveling predominantly within cortical layer 6 along the rostral sulcal bank and fundus (white arrowheads), and a U-shaped bundle (solid red line) coursing caudally around the CS in deeper portions of the white matter, leaving a region devoid of label between the bundle and the GM-WM border (solid green line). Panels A-B are from case PGR-R; Panels C-D are from case BMEX-L (reflected for comparison with PGR-R). Scale bars: A, C = 2 mm; B, D = 1 mm.

#### *3.2. Short Association Fibers along the Central Sulcus*

In both cases, labeled bundles were evident in the white matter beneath the central sulcus, originating as concentrated label beneath the injection site (Supplementary Figure 3B, D; solid red lines). Labeled profiles were initially longitudinal before transitioning to transverse at the sulcal fundus, indicating a rostrocaudal orientation. Following fibers across rostral and caudal sections confirmed a U-shaped trajectory throughout (Supplementary Figure 4B, D; dashed red lines).

In case PGR-R, the fiber bundle curved beneath the sulcal fundus, with label in the lower cortical layers at the fundus (Supplementary Figure 4B, white arrowheads), continuous with label in the underlying white matter. Fiber profiles transitioned from transverse to oblique and then longitudinal as the bundle fanned out within the postcentral gyral white matter, forming columnar terminations along the caudal bank in area 3b (Supplementary Figure 4B, red arrows) and more caudally on the postcentral gyral crown in cortical areas 1 and 2, supporting classification as a U-fiber. In case BMEX-L, labeling beneath the central sulcus resolved into two distinct intensity bands. The more superficial band coursed predominantly within cortical layer 6, extending only slightly into the subjacent white matter, with label in the lower cortical layers at the fundus and sulcal banks (Supplementary Figures 3D, 4D; white arrowheads), indicating labeling consistent with the intrinsic intracortical fiber network within layer 6. The deeper band occupied deeper portions of the white matter, maintaining a consistent distance from the GM-WM border and producing a region devoid of label between it and the cortex. Following a trajectory similar to the U-fiber in PGR-R, fiber profiles transitioned to oblique and then longitudinal after curving beneath the fundus before terminating in the lower cortical layers of areas 1 and 2 at the gyral crown (Supplementary Figure 4D, red arrows). Because the deeper bundle in BMEX-L interconnected opposing gyri with a U-shaped trajectory, it was classified as a U-fiber.

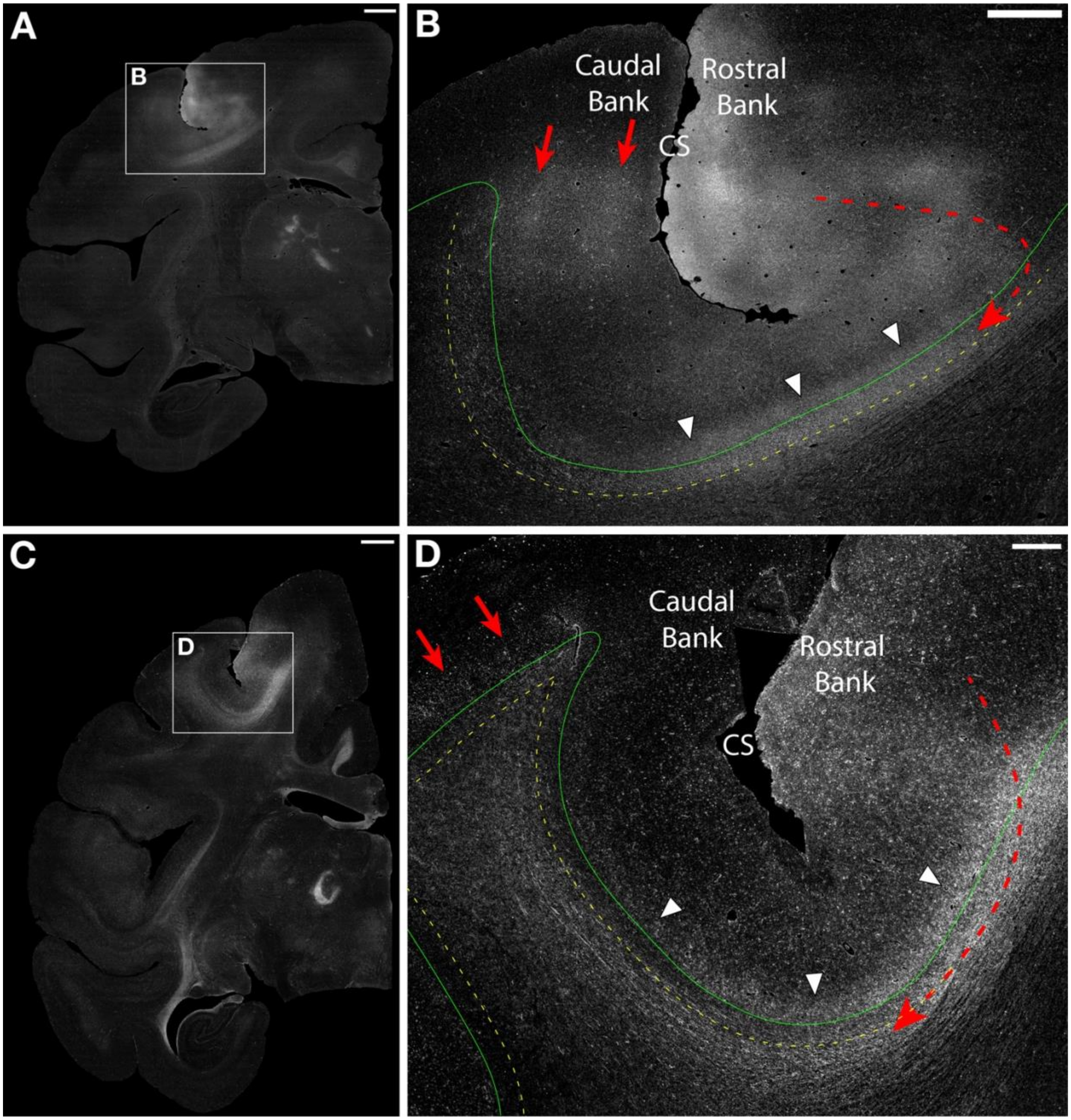


**Supplementary Figure 4:** Autoradiographs of short association fibers in the peri-Rolandic (motor/somatosensory) cortex of the macaque monkey. A. Low-magnification darkfield image ~4 mm caudal to an injection in the M1 hand representation on the precentral gyrus. B. Higher-magnification view of the boxed region in A showing a U-shaped bundle (dashed red line) coursing caudally around the central sulcus (CS) within the superficial white matter band (dashed yellow line) directly beneath the GM-WM border (solid green line). Label in the lower cortical layers at the fundus (white arrowheads) is continuous with the underlying white matter label. Labeled fibers fan out within the postcentral gyral white matter before providing columnar terminations along the caudal bank of CS (red arrows) and further caudally on the gyral crown, supporting its classification as a U-fiber. C. Low-magnification darkfield image ~6 mm caudal to an injection in the M1 trunk representation on the precentral gyrus. D. Higher-magnification view of the boxed region in C showing two distinct intensity bands beneath the CS: a superficial band traveling predominantly within cortical layer 6 along the sulcal banks and fundus (white arrowheads), and a deeper U-shaped bundle (dashed red line) coursing caudally around the CS and terminating in the lower cortical layers of areas 1 and 2 on the postcentral gyrus (red arrows). This U-fiber travels below the superficial white matter band (dashed yellow line), creating a region devoid of label beneath the GM-WM border (solid green line), most evident along the caudal bank of CS. Panels A-B are from case PGR-R; Panels C-D are from case BMEX-L (flipped for direct comparison with PGR-R). Scale bars: A, C = 2 mm; B = 1 mm; D = 500 μm.

Despite similar overall organization, the two U-fibers differed in their depth relative to the GM-WM border and terminal field position on the postcentral gyrus. In PGR-R, the injection was positioned close to the sulcus, and terminations were located on the corresponding portion of the postcentral gyrus close to the central sulcus. Reflecting this shorter trajectory around the sulcus, labeled fibers in PGR-R remained directly beneath the GM-WM border throughout, fully occupying the superficial white matter band (Supplementary Figure 4B, dashed yellow line); the deep margin of the bundle at the sulcal fundus measured 283 μm from the GM-WM border (SD 13.7 μm; range 270-297 μm). The apparent increase in band depth along the caudal bank reflects the oblique plane of section rather than a true increase in bundle depth. In BMEX-L, the injection was positioned farther from the sulcus, and the terminal field was located more caudally on the postcentral gyrus, closer to the intraparietal sulcus (IPS), reflecting the greater injection-to-sulcus distance. Labeled fibers in BMEX-L traveled below the superficial white matter band throughout their trajectory (Supplementary Figure 4D, dashed yellow line), with the superficial margin measured at a mean depth of 245 μm from the GM-WM border (SD 27.0 μm; range 214-283 μm). This increased depth accords with Meynert's observation that longer U-fibers interconnecting regions on gyri more distant from the sulcus travel deeper in the white matter. Despite this difference in depth, labeled fibers in both cases maintained a consistent orientation orthogonal to the long axis of the sulcus, with terminations on the corresponding portion of the adjacent gyrus, supporting a symmetric U-fiber classification.

#### *3.3 Intragyral Fibers*

In addition to short association fiber bundles coursing beneath and along the central sulcus, local intragyral fibers were observed in both cases. These fibers traveled within the precentral gyrus to terminate medially in cortical area 4 and more rostrally in area 6. In coronal sections, these intragyral fibers appeared as oblique and transverse profiles within the white matter beneath the gyrus as well as within cortical layer 6.

#### *3.4 Summary*

The U-fibers identified beneath the central sulcus in both motor cortex cases matched the classical conception of symmetric U-fibers. That is, they maintained an orientation orthogonal to the long axis of the sulcus and terminated on the corresponding portion of the opposing postcentral gyrus. However, unlike the prefrontal cases, the two U-fibers differed in their depth within the white matter: the U-fiber in PGR-R traveled within the superficial white matter band directly beneath the GM-WM border, whereas the U-fiber in BMEX-L coursed below this band, in accordance with Meynert's rule that longer U-fibers interconnecting regions on gyri more distant from the sulcus travel at a greater depth from the cortex.

### 4. Temporal Lobe

The white matter was analyzed along the rostrocaudal extent of the superior temporal sulcus (STS) in 10 cases. The STS is a prominent operculated sulcus that begins rostrally proximal to the temporal pole and runs dorsocaudally along the lateral surface of the temporal lobe. In more posterior locations, it ascends dorsally to intersect with the IPS, the lunate sulcus, and the parieto-occipital medial sulcus. U-shaped bundles linking the superior temporal gyrus and inferior temporal gyrus around the STS were observed in only one case, BMJ (see Table 1).

#### *4.1 Injection Sites and Long-Range Fiber Systems*

In case BMJ, the injection was placed in the rostral superior temporal gyrus, centered on cortical areas Pro and TS1 with a slight extension into area TS2 (Supplementary Figure 5A-B, red asterisks). The injection produced a densely labeled caudally directed cord that descended ventrally through the deep white matter (Supplementary Figure 5B, solid black line), giving rise to several subcortically directed fiber systems. A large medial bundle entered the anterior commissure to reach the opposite hemisphere, whereas a smaller medial bundle followed a similar course but passed beneath the anterior commissure to terminate ipsilaterally in the amygdala and basal forebrain. The remaining subcortical bundle traveled caudally between the tail of the caudate and the putamen, where it divided into fiber bundles projecting to the thalamus and brainstem. Cortico-striatal fibers, separate from the cord fibers, were also observed traveling dorsally to reach the claustrum and then entering the external capsule to reach the caudate nucleus and putamen. Another group of cortico-striatal fibers also coursed caudally to reach the tail of the caudate and the ventral putamen. Label was also observed in several long association fiber pathways (Supplementary Figure 5B, solid white lines), such as the MdLF and uncinate fasciculus. These long association fiber bundles were readily differentiated from other non-cord fibers based on their location in the deep white matter and/or their characteristic stippled organization formed by predominantly transverse fiber profiles.

*4.2 Short Association Fiber Bundles along the Superior Temporal Sulcus*

The remaining label in case BMJ comprised short association fibers observed in the white matter beneath the STS, forming a U-shaped bundle (Supplementary Figure 5B, solid red line). The fiber bundle consisted of densely labeled fibers observed predominantly as oblique and transverse profiles that coursed in the white matter between the cord and the GM-WM border along the superior bank of the STS. As the bundle continued ventrally around the fundus and along the inferior bank of the STS, labeled fibers fanned out with a roughly rostrocaudal orientation before terminating in the upper cortical layers of areas TEa and TEm on the sulcal bank and inferior temporal gyrus (Supplementary Figure 5B, red arrows). At the depth of the STS, the deep margin of the fiber bundle was located at a mean depth of 250 μm from the GM-WM border (SD 20.3 μm; range 232-279 μm). These measurements indicated that this fiber bundle occupied the superficial white matter band. Given its U-shaped trajectory and traceable continuity from the injection to terminations on the corresponding portion of the adjacent inferior temporal gyrus, we classified this bundle as a symmetric U-fiber that matched the classical definition.

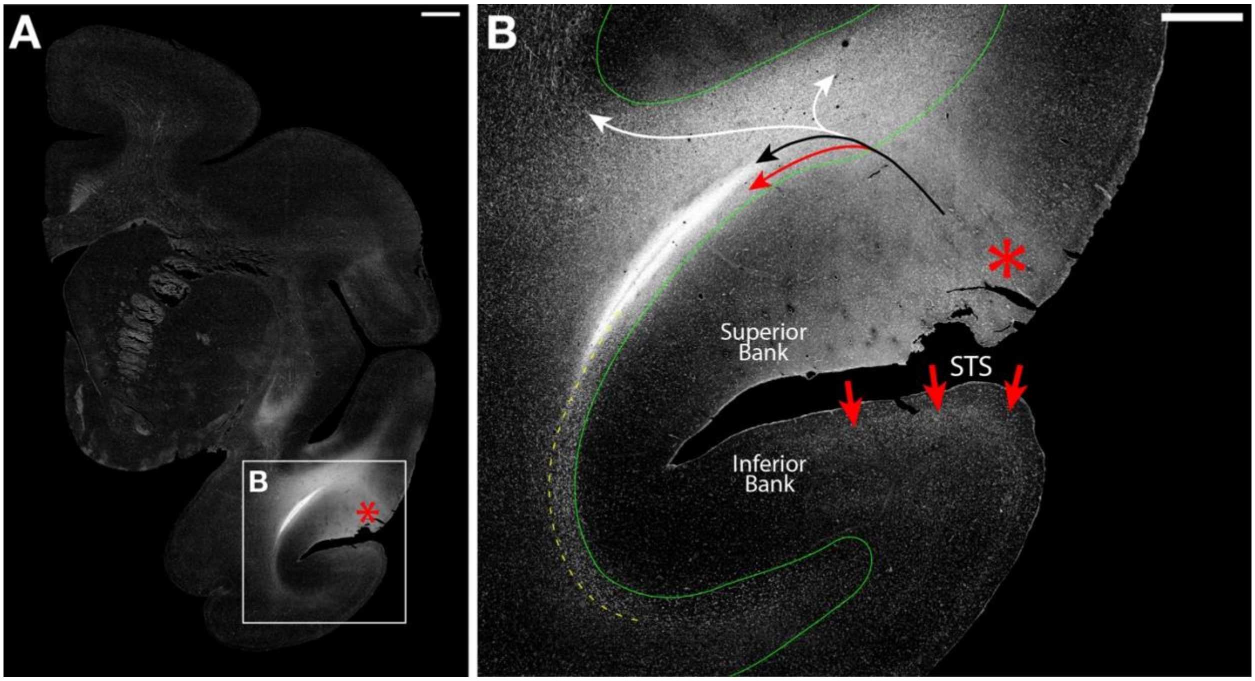


**Supplementary Figure 5:** Autoradiographs of short association fibers in the rostral temporal lobe of the macaque monkey. Coronal section from one case processed for autoradiography and imaged under darkfield illumination to reveal labeled fibers. A. Low-magnification darkfield microscopic image following an injection in the rostral superior temporal gyrus centered on cortical areas Pro and TS1 and extending into TS2 (red asterisk). B. High-magnification view of the boxed region in A showing multiple labeled fiber bundles, including a U-shaped bundle (solid red line) coursing beneath the superior temporal sulcus (STS), cord fibers within the temporal white matter (solid black line), and several long association fiber pathways (solid white lines). The U-shaped bundle travels within the white matter directly below the gray matter-white matter border (solid green line) within the superficial white matter band (dashed yellow line), as it courses ventrally around STS, before forming terminations along the inferior bank of STS and crest of inferior temporal gyrus (red arrows), consistent with a U-fiber. Images from case BMJ. Scale bars: A = 2 mm; B = 1 mm.

*4.3 Intragyral Fibers*

Labeling corresponding to local intragyral fibers were also observed coursing from the injection site across the white matter of the superior temporal gyrus, terminating in the upper cortical layers of area TS2 on the gyral crown and in area Pro along the Sylvian (lateral) fissure.

*4.4 Summary*

The U-fiber identified beneath the rostral STS in case BMJ matched the classical conception of U-fibers proposed by Meynert. That is, it was oriented orthogonally to the long axis of the sulcus, terminated on the corresponding portion of the adjacent inferior temporal gyrus at approximately the same rostrocaudal level as the injection, and traveled within the superficial white matter band directly beneath the GM-WM border. This bundle therefore satisfied all three criteria for classification as a superficial ('true') symmetric U-fiber.

### 5. Parietal Lobe

The sulcal white matter was analyzed at several points along the rostrocaudal extent of the IPS in 14 cases. The IPS begins anteriorly caudal to the postcentral region and extends dorsally and caudally to join the STS, lunate sulcus, and parieto-occipital medial sulcus. Rostrally, the IPS is relatively shallow, whereas more caudally it deepens and branches, with features characteristic of an operculated sulcus. This morphological change allowed us to examine U-shaped bundles both in a larger, more complex sulcus and in its shallower rostral extension.

#### 5.1 Rostral Intraparietal Sulcus

*5.1.1 Injection Site and Long-Range Fibers*

At the rostral IPS, three cases were examined with injections on the gyrus forming the lateral bank of the sulcus. Case BMDH had an injection on the dorsal opercular surface of the lateral sulcus centered on cortical areas PFop and SII (Supplementary Figure 6A-B, red asterisk), whereas cases BMX and MRG had injections centered on cortical area PF, with case BMX encroaching on area PFG (Supplementary Figure 6C-D, red asterisk). In all three cases, the injection produced a central cord of densely labeled fibers (Supplementary Figure 6B, D; upper solid white lines) that coursed medially through the deep white matter of the inferior parietal lobule (IPL) before splitting into two bundles. The more dorsal bundle continued medially through the corona radiata toward the corpus callosum, whereas the more ventral bundle entered the internal capsule, where it separated into fiber bundles projecting to the thalamus and brainstem. A separate cortico-striatal bundle was observed ventral to the cord (Supplementary Figure 6B, D; lower solid white lines), traveling around the superior bank of the lateral fissure before entering the external capsule to project to the putamen in all cases, with additional projections through the external and extreme capsules to the claustrum in cases BMX and MRG. Labeled fibers corresponding to long association fiber pathways (e.g., SLF I, III, cingulum bundle), were also present outside the cord and coursed rostrally and caudally from the injection sites. These fibers were identified based on their deep position in the white matter, their position relative to the large bundles described above, and their stippled appearance of transverse fiber profiles.

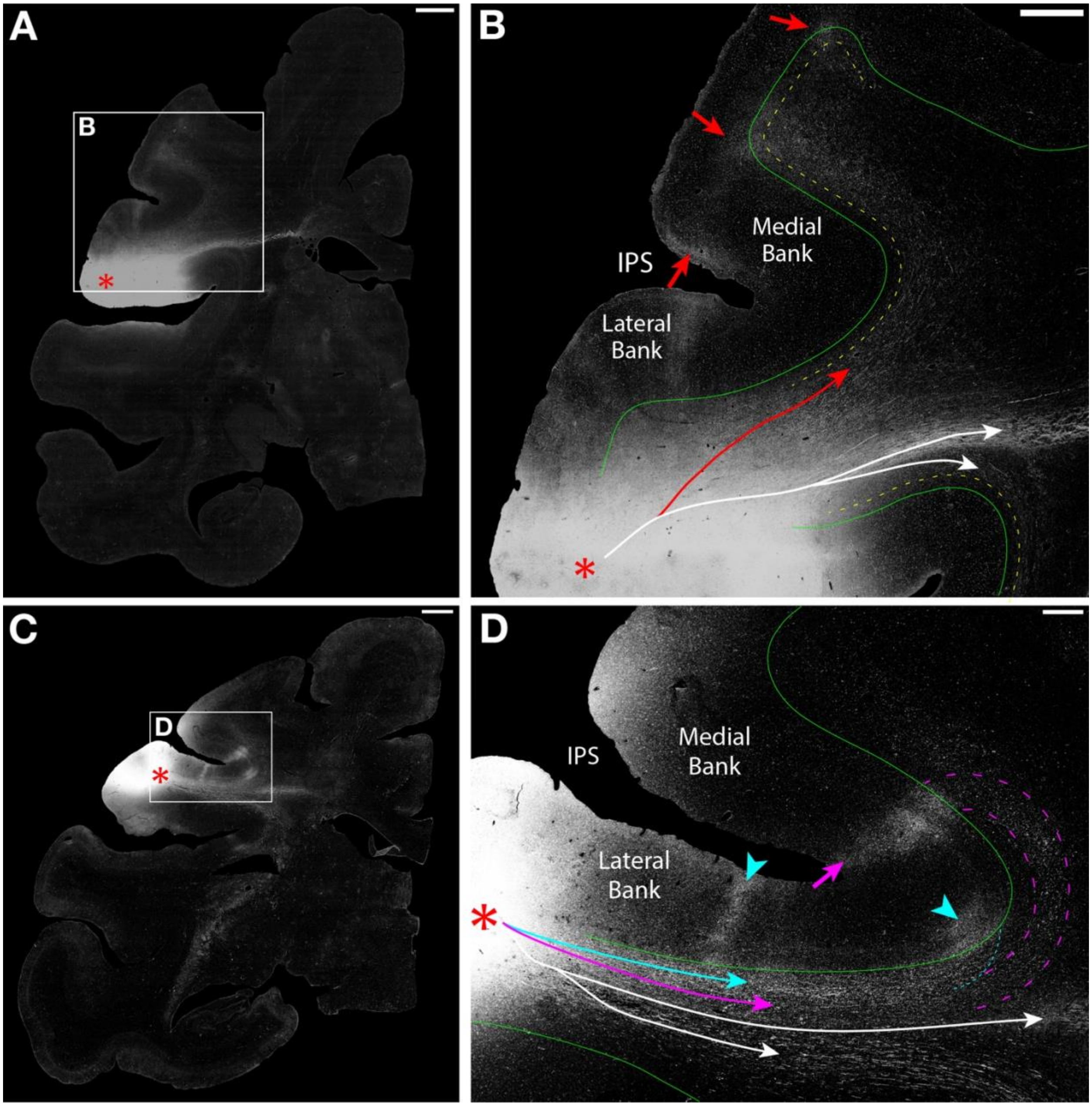

**Supplementary Figure 6:** Autoradiographs of short association fibers in the rostral parietal lobe of the macaque monkey. This figure shows coronal sections from two macaque inferior parietal lobule (IPL) cases processed for autoradiography and imaged under darkfield illumination to reveal labeled fibers. A. Low-magnification darkfield image following an injection on the dorsal opercular surface of the lateral sulcus in cortical areas PFop and SII (red asterisk). B. Higher-magnification view of the boxed region in A highlighting multiple labeled fiber bundles, including a U-shaped bundle (solid red line) coursing dorsally around the intraparietal sulcus (IPS), cord fibers within the deep white matter projecting medially (upper solid white line), and ventrally positioned cortico-striatal fibers (lower solid white line). The U-shaped bundle around the IPS travels in deeper portions of the white matter and largely avoids the superficial white matter band (dashed yellow line) beneath the gray matter-white matter (GM-WM) border (solid green line). After crossing the sulcal fundus, labeling extends into the cortex along the medial bank of IPS and within areas 1 and 2 on the adjacent gyrus (red arrows), supporting classification as a U-fiber. C. Low-magnification darkfield image following an injection in the rostral IPL in cortical areas PF and PFG (red asterisk). D. Higher-magnification view of the boxed region in C showing an L-type bundle (solid teal line), a J-type bundle (solid magenta line; outlined by dashed magenta lines), cord fibers within deep white matter (upper solid white line), and ventrally positioned cortico-striatal fibers (lower solid white line). The L-type bundle courses from the injection within the white matter immediately deep to the GM-WM border (solid green line), with terminations observed along the lateral bank and fundus of the IPS (teal arrowheads). In contrast, the J-type bundle travels medially along the lateral bank of the IPS in deeper portions of the white matter and hooks dorsally around the sulcus to enter the superior parietal lobule, with terminations along the medial bank of the IPS (magenta arrow). Panels A-B are from case BMDH; Panels C-D are from case BMX. Scale bars: A, C = 2 mm; B, D = 500 μm.

#### *5.1.2 Short Association Fiber Bundles along the Intraparietal Sulcus*

Labeling was evident beneath the GM-WM interface of the rostral IPS in three cases (BMDH, BMX, MRG), forming distinct local short association fiber bundles. However, their organization and relationships differed based on their injection sites. In case BMDH, the injection was centered on cortical areas PFop and SII on the more lateral aspect of the gyrus (Supplementary Figure 6A-B, red asterisk). Labeled fibers formed a U-shaped bundle (Supplementary Figure 6B, solid red line) coursing beneath the IPS into the superior parietal lobule (SPL), terminating in the upper cortical layers on the medial IPS bank and in the upper and lower layers of cortical areas 1 and 2 on the adjoining gyrus (red arrows), supporting classification as a U-fiber. Throughout its trajectory, this bundle avoided the superficial white matter band (Supplementary Figure 6B, dashed yellow line), with the superficial margin of the bundle located at a mean depth of 201 μm from the GM-WM border (SD 18.5 μm; range 173-213 μm), entering the band only beneath its cortical terminations. In adjacent caudal sections, this U-shaped bundle was traceable beneath the IPS approximately 4 mm caudal to the main injection foci, providing terminations in cortical area 2 on the opposing gyrus, with additional terminations in cortical area POa on the medial IPS bank, IPd at the fundus, and PEa on the lateral IPS bank.

In case BMX, the injection was centered on cortical area PF, encroaching on area PFG (Supplementary Figure 6C-D, red asterisks). Two distinct fiber bundles were identified along the lateral IPS bank. The first bundle (Supplementary Figure 6D, solid teal line) coursed directly beneath the GM-WM border, with overlying terminations observed on the lateral IPS bank in cortical area POa and at the fundus in cortical area IPd (teal arrowheads). Labeled fibers in this more superficial bundle did not progress past its termination at the fundus (Supplementary Figure 6D, dashed teal line), with the deep margin of the bundle located at a mean depth of 270 μm from the GM-WM border (SD 17.0 μm; range 250-291 μm), confirming its position within the superficial white matter band. This bundle was classified as an L-type connection based on its relatively straight trajectory from the injection site along the proximal sulcal bank and terminations that did not progress to the opposing sulcal bank or gyrus. The second bundle (Supplementary Figure 6D, solid magenta line) traveled in deeper portions of the white matter; once the first bundle ended (dashed teal line), a gap devoid of label was evident between the GM-WM border and this deeper bundle (dashed magenta lines), with the superficial margin located at a mean depth of 265 μm from the GM-WM border (SD 50.5 μm; range 207-301 μm). This bundle continued around the sulcus into the SPL to terminate on the medial IPS bank in cortical area IPd (magenta arrow), supporting classification as a J-type connection. A similar L-type connection was observed in case MRG, without an accompanying J-type connection.

The distinction between the two bundle types in case BMX is further evident in sections through the caudal extent of the injection site, where the IPS was slightly deeper (Supplementary Figure 7A-B). At this level, the L-type bundle persists with terminations along the lateral IPS bank in cortical area POa (Supplementary Figure 7B, teal arrows), whereas the J-type bundle is absent. This case illustrates the depth-connectivity relationship described by Meynert: the more superficial L-type bundle terminated after a shorter course along the proximal bank and fundus, whereas the deeper J-type bundle traveled farther around the sulcus to reach the medial bank.

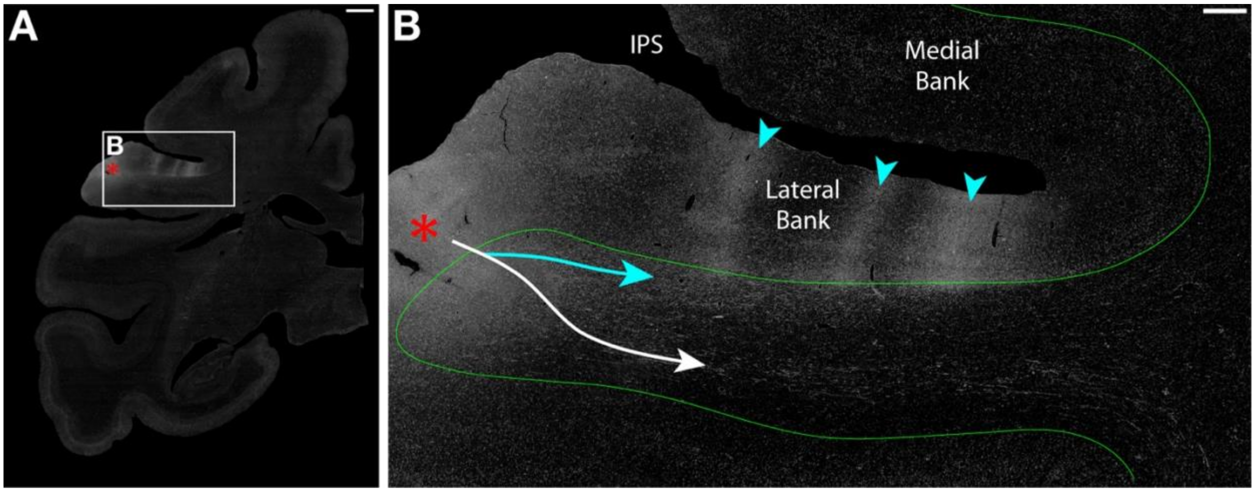


**Supplementary Figure 7:** Autoradiographs of short association fibers in the rostral parietal lobe of the macaque monkey. Coronal section from one case processed for autoradiography and imaged under darkfield illumination. A. Low-magnification darkfield image at the caudal extent of the injection site in the rostral inferior parietal lobule in cortical areas PF and PFG (red asterisk), where the depth of the IPS is slightly greater than at the level shown in Supplementary Figure 6C-D. B. Higher-magnification view of the boxed region in A showing multiple labeled bundles: ventrally positioned cortico-striatal fibers (solid white line) and an L-type bundle (solid teal line). Similar to the more rostral section, the L-type bundle courses directly beneath the gray matter-white matter border (solid green line) with a series of columnar terminations along the lateral IPS bank (teal arrowheads). At this level, the J-type bundle observed more rostrally (Supplementary Figure 6D, solid magenta line) is absent, no labeled fibers curve around the fundus and no terminations are present on the medial IPS bank. Images from case BMX. Scale bars: A = 2 mm; B = 500 µm.

It is important to note that in both cases, fibers coursing around the depth of the IPS did not remain in the coronal plane, including the U-fiber in BMDH and the J-type connection in BMX. Fibers along the lateral IPS bank were observed primarily as longitudinal profiles, indicating a trajectory roughly within the plane of section and perpendicular to the local long axis of the sulcus. After coursing beneath the fundus, profiles shifted to transverse, indicating a rostrocaudal trajectory out of the plane of section. This interpretation was supported by the diverging fan-shaped labeling observed in adjacent rostral and caudal sections. On this basis, the U-fiber in case BMDH and the J-type connection in case BMX were both classified as asymmetric, as labeled fibers did not maintain a consistent orientation orthogonal to the local long axis of the sulcus throughout their trajectory. The asymmetric classification in case BMDH is further supported by the observation that labeled fibers formed a U-shaped bundle beneath the IPS approximately 4 mm caudal to the main injection foci. By contrast, the L-type connection in case BMX was classified as symmetric, as labeled fibers maintained a consistent orientation perpendicular to the long axis of the sulcus throughout their straight trajectory along the proximal sulcal bank without crossing the fundus.

*5.1.3 Intragyral Fibers*

In all cases, local intragyral fibers were observed. In case BMX, these fibers were observed as predominantly as transverse profiles within the superficial portions of the IPL white matter. Labeled fibers coursed rostrally to provide terminations in primary somatosensory cortex and caudally to provide terminations in cortical areas PFG and PGop. At the level of the injection site in case BMDH, intragyral fibers provided a columnar termination in cortical area PF on the lateral bank of the IPS. At the caudal extent of the injection site, columnar terminations were observed in cortical areas PFG and PG. Transverse fiber profiles, indicating a rostrocaudal orientation, were also observed coursing from the injection site within the gyral white matter of the IPL, providing terminations rostrally in the upper cortical layers of areas PF and 2, as well as caudally in cortical area PGop.

*5.1.4 Summary*

Short association fiber bundles at the rostral IPS illustrated two organizational principles. In case BMDH, a U-fiber interconnected the IPL with the medial IPS bank and adjoining SPL gyrus, traveled below the superficial white matter band, and was classified as asymmetric based on its oblique trajectory relative to the long axis of the sulcus and terminations caudal to the injection site. In case BMX, the absence of U-fibers was accompanied by L- and J-type connections, with the shorter L-type connection traveling superficially within the superficial band and terminating along the proximal sulcal bank, and the deeper J-type connection traveling farther around the fundus to terminate on the medial IPS bank, in accordance with Meynert's observation that shorter fibers occupy more superficial positions within the white matter whereas longer fibers travel deeper.

#### 5.2 Mid Intraparietal Sulcus

Six of the 14 parietal cases investigated had injections along the mid-portion of the IPS: three in the SPL and three in the IPL. In the mid-IPS region, the sulcus is deeper than in the rostral IPS described in the previous section, and multiple anatomically and physiologically defined areas are present along the banks and gyri. Across these six cases, U-shaped bundles were identified only in the three SPL-injected cases; no U-shaped bundles were observed in the three IPL-injected cases (Table 1).

#### Superior Parietal Lobule Cases

##### *5.2.1 Injection Site and Long-Range Fibers*

SPL injection sites were centered on cortical area PEc in cases BMP and BMT and on cortical area PGm, extending into cortical area PEc, in case BMD. These cases exhibited prominent rostrally directed cord fibers consisting of densely labeled fibers that traveled ventrally from the injection site within the core of the SPL white matter before curving beneath the fundus of the IPS (Supplementary Figure 8B, dashed black line) and splitting into two separate bundles. The more medial bundle continued medially, entering the splenium of the corpus callosum to reach the opposite hemisphere. The more ventrolateral bundle entered the superior aspect of the sagittal stratum and continued rostrally into the internal capsule, where it divided into thalamic and pontine components. In all three cases, a distinct group of cortico-striatal fibers separate from the cord coursed rostrally from the injection site, traveling through the FOF into Muratoff's bundle before continuing rostrally to terminate in the caudate nucleus. Some fibers continued laterally from Muratoff's bundle, passing through the superior portion of the internal capsule to terminate in the putamen, with additional fibers coursing through the external capsule to reach the claustrum. In cases BMP and BMT, an additional group of fibers coursed laterally through the corona radiata before entering the external capsule to reach the claustrum. Labeled fibers were also observed in long association fiber pathways (e.g., SLF I, cingulum bundle, FOF) in all three cases that were identifiable based on their deep position in the white matter and the orientation of their fibers.

*5.2.2 Short Association Fiber Bundles along the Intraparietal Sulcus*

Prominent U-shaped fiber bundles (Supplementary Figure 8B, dashed red line) were observed lateral to the cord at the depth of the IPS in all three cases. These rostrally directed fibers emerged from the injection site as predominantly transverse profiles, coursing ventrally through the SPL white matter before curving around the IPS fundus to enter the center of the IPL white matter. At the depth of the IPS, fiber profiles transitioned from transverse to longitudinal profiles within the plane of section (see Figure 17B), continuing dorsally toward the IPL gyral crown, where profiles progressively shifted to oblique and then transverse as fibers fanned out beneath the gyrus. Fibers crossed the GM-WM interface to terminate in cortical areas PG and PGop in case BMD, cortical area PG in case BMP (Supplementary Figure 8B, red arrow), and cortical area PGop in case BMT, with slight differences in termination sites between BMP and BMT reflecting differences in injection foci. In all three cases, these labeled fibers were traceable from the injection on the crown of the SPL, beneath the IPS, to the adjacent IPL gyral crown, supporting classification as U-fibers. However, the terminal fields in the IPL were notably rostral to the injection site (see Figure 10E), and the orientation of labeled fiber profiles throughout their trajectory indicated that the fibers did not cross the sulcus perpendicular to its long axis. Instead, they adopted a considerably oblique trajectory; these bundles were therefore classified as asymmetrical. Throughout their trajectory around the IPS, labeled fibers followed the contour of the IPS while avoiding the superficial white matter band (Supplementary Figure 8B, dashed yellow line), maintaining a consistent depth from the GM-WM border until approaching their cortical terminations. In case BMP, the superficial margin of the bundle was located at a mean depth of 236 μm from the GM-WM border (SD 22.5 μm; range 218-280 μm). Together, these features indicate that the U-fibers in all three SPL cases were deeper U-fibers with an asymmetric trajectory around the sulcus.

Shorter L-type connections (Supplementary Figure 8B, dashed teal line) were also identified in all three SPL cases. These bundles were observed as densely labeled strips of mixed fiber profiles descending ventrally from the injection site within the superficial white matter band along the medial IPS bank. In cases BMD and BMP, fibers terminated in cortical area PO in a columnar fashion caudal to the injection, whereas in case BMT, fibers extended slightly farther caudally to reach the same target. In all three cases, the remaining fibers continued rostrally within the superficial white matter band, providing columnar terminations in cortical area PEa along the medial IPS bank (Supplementary Figure 8B, teal arrowheads). Although terminal fields were similar across cases, minor variations in injection foci produced slight differences in trajectory. These bundles were positioned more superficially than the U-fibers described above. They were classified as asymmetric based on their oblique trajectory relative to the long axis of the IPS, as confirmed by examination of fiber profiles across multiple coronal sections and by the location of their terminations along the medial IPS bank, approximately 4 mm rostral to the injection site and further caudally.

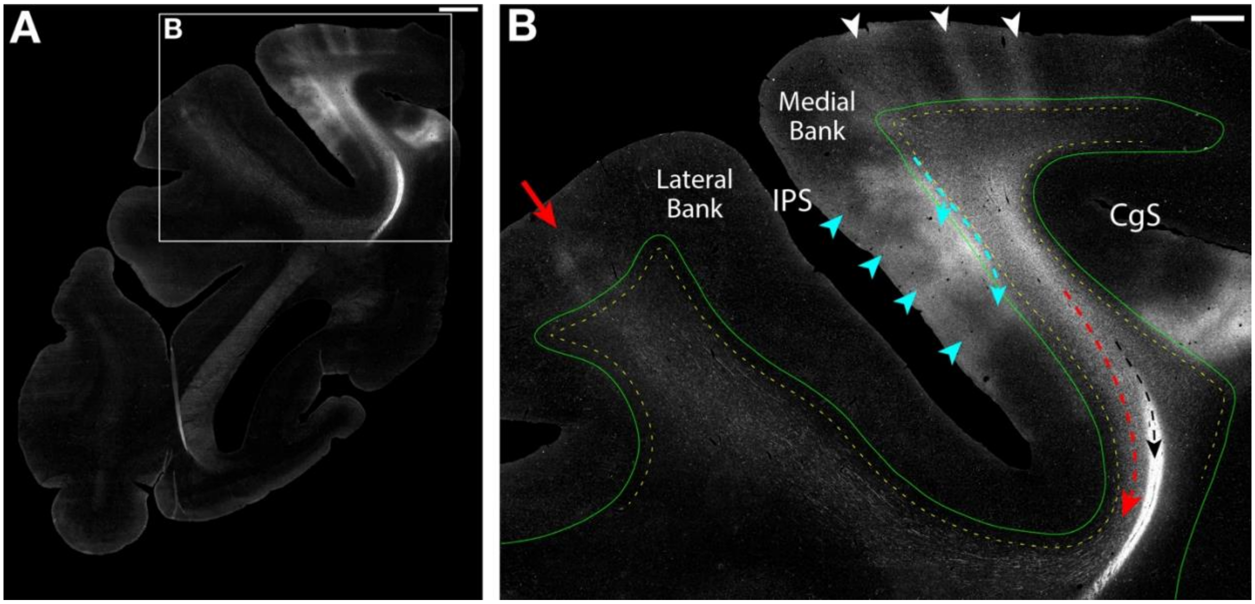


**Supplementary Figure 8:** Autoradiographs of short association fibers in the parietal lobe of the macaque monkey. A. Low magnification darkfield image ~4 mm rostral to an injection site in the superior parietal lobule (SPL) centered on cortical area PEc. B. Higher-magnification view of the boxed region in A showing multiple labeled fiber bundles, including a U-shaped bundle (dashed red line) coursing ventrally around the intraparietal sulcus (IPS), cord fibers within the deep white matter (dashed black line), and an L-type bundle (dashed teal line). The U-shaped bundle curves around the fundus of IPS, enters the center of the inferior parietal lobule white matter, and continues laterally before forming terminations on the gyral crown (red arrow). Throughout its trajectory, this bundle travels in deeper portions of the white matter and largely avoids the superficial white matter band (dashed yellow line) immediately subjacent to the gray matter-white matter (GM-WM) border (solid green line). In contrast, the L-type bundle courses rostrally from the injection site within the superficial white matter immediately deep to the GM-WM border and forms columnar terminations along the medial bank of the IPS (teal arrowheads). Additional columnar terminations are present on the crown of the SPL (white arrowheads). Images from case BMP. Scale bars: A = 2 mm; B = 1 mm. CgS, Cingulate Sulcus.

#### *5.2.3 Intragyral Fibers*

As in previous cases, labeled transverse fiber profiles corresponding to intragyral fibers were observed traveling within the white matter below the gyral crest, providing columnar terminations both rostral to their respective injection sites in cortical area PE (Supplementary Figure 8B, white arrowheads) and caudally in cortical area PEc. Columnar terminations were also observed across the SPL on the medial hemisphere in cortical area PGm in all three cases.

*5.2.4 Summary*

U-fibers were identified in all three SPL cases, interconnecting the SPL gyral crown with the adjacent IPL gyral crown across the IPS. In contrast to the prefrontal U-fibers, these bundles represented a different subpopulation of U-fibers that traveled below the superficial white matter band throughout their trajectory and were classified as asymmetric because their terminal fields in the IPL were notably rostral to the injection site rather than at the corresponding level across the sulcus. This interpretation is supported by the observed transverse fiber profiles in the SPL, which indicate an orientation parallel to the long axis of the sulcus and suggest that these U-fibers had an overall oblique trajectory. Shorter L-type connections were also identified in all three cases, coursing within the superficial white matter band along the medial IPS bank. Like the U-fibers, these L-type bundles followed an oblique trajectory and were classified as asymmetric. Together, both fiber types at the mid-IPS illustrate that U-fibers and shorter local connections can coexist beneath the same sulcus, with U-fibers traveling deeper and L-type connections more superficially, in accordance with Meynert's rule of a depth-connectivity relationship.

**Inferior Parietal Lobule Cases**

*5.2.5 Injection Site and Long-Range Fibers*

Of the three cases with IPL injections, one case (BML) had labeled fibers observed along the sulcal white matter of the IPS. In case BML, the injection (Supplementary Figure 9A-B, red asterisks) was located on the crown of the IPL, centered on the caudal portion of cortical area PG and extending into area Opt. The injection resulted in two parallel densely labeled cords that traveled medially through the deep white matter of the IPL (Supplementary Figure 9B, medial solid black line). The more dorsal cord continued medially traveling to and entering the splenium of the corpus callosum and the more ventral cord entered the sagittal stratum before continuing rostrally toward the internal capsule where it divides into a thalamic and pontine components. Other, more diffuse non-cord fibers were positioned medially and laterally to the cord. Cortico-striatal fibers traverse through the corona radiata to enter the FOF and then Muratoff's bundle to terminate in the caudate nucleus. The other contingent of striatal fibers descend from the core of the IPL into the external capsule and terminate in the claustrum and putamen. The majority of fibers lateral to the cord constitute long association fiber pathways (e.g., SLF I, II, ILF, MdLF) and were identified from other non-cord fibers based on their position in the deep white matter and stippled labeling (Supplementary Figure 9B, lateral solid black lines).

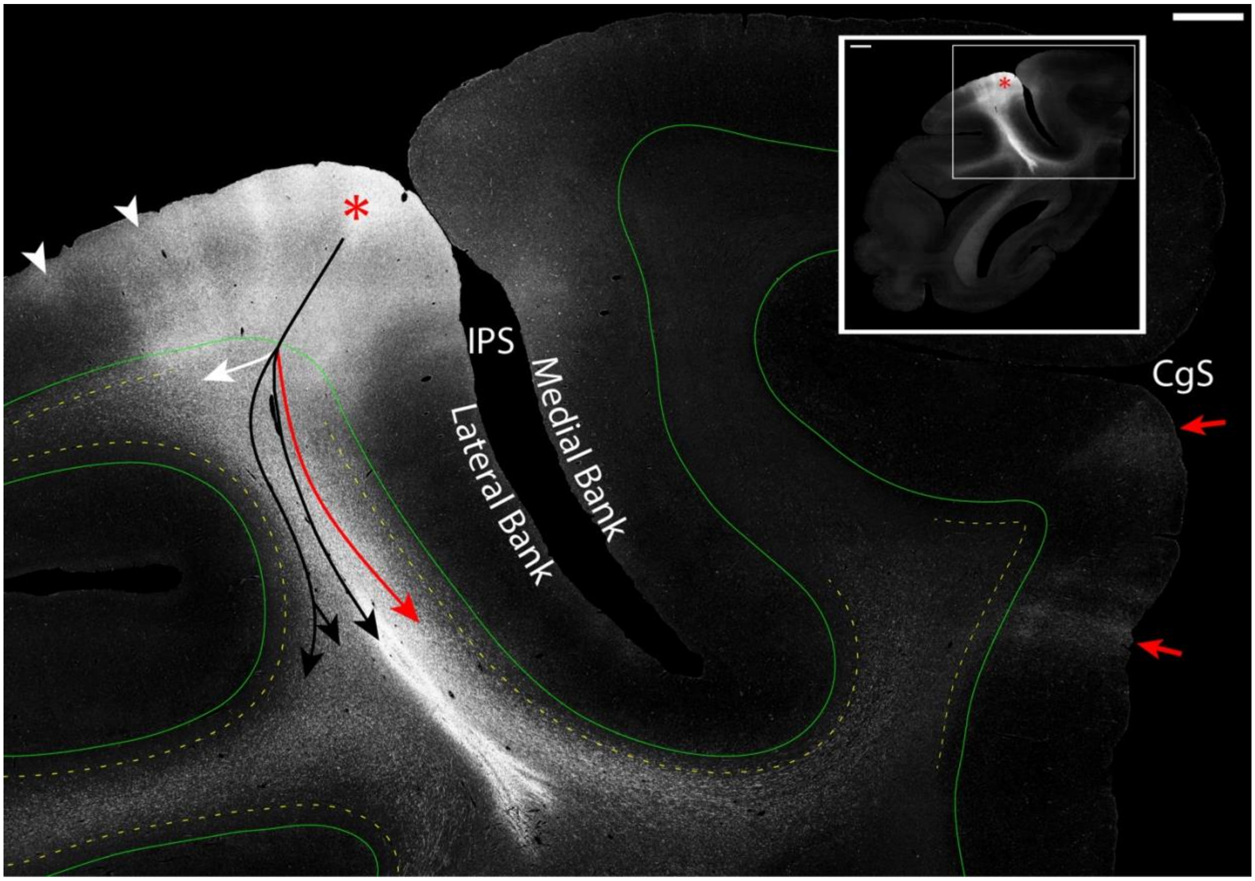


**Supplementary Figure 9:** Autoradiographs of longer association fibers below the mid portion of the intraparietal sulcus (IPS) that appear U-shaped and do not interconnect adjacent gyri. Main panel: Higher-magnification darkfield image from the region indicated in the inset showing multiple labeled fiber bundles, including a U-shaped bundle (solid red line) several labeled cord and long association fiber pathways (solid black lines), and intragyral fibers (solid white line) providing terminations on the injected gyrus (white arrowheads). Labeled fibers from the injection (red asterisk) course along the medial bank of the IPS and hook beneath the sulcal fundus, initially traveling along the opposing bank in a manner resembling a U-fiber. However, within the superior parietal lobule these fibers turn medially, traverse the gyral core, and form terminations on the medial hemispheric cortex (red arrows) below the cingulate sulcus (CgS). Throughout its trajectory, this bundle travels in deeper portions of the white matter, remaining below the superficial white matter band (dashed yellow line). Inset: Low-magnification darkfield image following an injection in the inferior parietal lobule centered in the caudal portion of cortical area PG and extending into area Opt (red asterisk). Images from case BML. Scale bars: main panel = 1 mm; inset = 2 mm.

#### *5.2.6 Fiber Bundles Along the Intraparietal Sulcus*

The majority of the remaining labeled fibers formed a prominent bundle coursing ventrally through the white matter core of the IPL (Supplementary Figure 9B, solid red line). At the level of the injection, the bundle consisted predominantly of longitudinal fiber profiles that coursed toward and hooked beneath the IPS fundus. Upon entering the SPL, labeled fibers briefly traveled along the opposing sulcal bank, resembling the U-shaped trajectory associated with U-fibers; however, rather than continuing toward the SPL crown, fibers turned medially and coursed obliquely across the gyral white matter core, diverging into medially and caudally directed components. The medially directed fibers provided columnar terminations in cortical area PGm, with some fibers continuing dorsally to reach cortical area PEci (Supplementary Figure 9B, red arrows), whereas the caudally directed fibers terminated in more caudal portions of cortical area PGm. A similar organization was observed rostral to the injection site, where transverse profiles within the center of the IPL white matter hooked around the IPS fundus before turning medially and traveling across the white matter to terminate in cortical area 31 on the medial surface of the hemisphere. The terminal field on the medial surface of the hemisphere therefore extended approximately 10 mm rostrocaudally. Throughout its trajectory, this bundle avoided the superficial white matter band (Supplementary Figure 9B, dashed yellow line), with the superficial margin at a mean depth of 260 μm from the GM-WM border (SD 30.0 μm; range 225-294 μm). Although this bundle adopted a U-shaped trajectory beneath the IPS, it did not terminate on the medial bank, fundus, or adjoining gyral crown. Rather than wrapping around the sulcus to the SPL crown as a U-fiber would, the bundle turned medially at the fundus to terminate on the medial surface of the hemisphere, constituting a longer association bundle rather than a local short association connection.

At the caudal extent of the injection, two additional fiber bundles were identified within the superficial white matter band: one along the IPS and one along the caudal STS, described below. Along the IPS, labeled fibers coursed ventromedially from the injection site along the lateral IPS bank as a densely labeled strip directly beneath the GM-WM border, consisting predominantly of longitudinal and oblique profiles. This bundle provided a series of columnar terminations in cortical area POa along the lateral IPS bank and a continuous strip of terminations extending into cortical area IPd at the sulcal fundus. Intracortical fibers fanning out through the cortical gray matter at the gyral crest were also associated with these terminations. Based on its straight trajectory from the injection site along the proximal sulcal bank, orientation perpendicular to the long axis of the sulcus, and terminations that did not extend beyond the IPS fundus, this bundle was classified as a asymmetric L-type connection.

*5.2.7 Short Association Fiber Bundles along the Superior Temporal Sulcus*

At the caudal extent of the injection in case BML, a labeled fiber bundle was identified beneath the caudal STS. At this level, the sulcus is relatively shallow and borders fewer anatomically and physiologically distinct cortical areas than at its mid-portion. Although case BML was selected as a parietal IPS case, the relatively large injection involved cortical area Opt on the gyrus dorsal to the STS, producing a labeled fiber bundle beneath the caudal STS alongside the IPS fibers described above. This bundle consisted of mixed longitudinal and oblique fiber profiles coursing ventromedially from the injection site and curving around the STS to enter the preoccipital gyrus. Terminations were observed in the upper cortical layers of area Opt along the STS bank and in areas V4d and DP on the opposing gyrus at approximately the same level as the injection. As the bundle originated from a gyral injection site, adopted a U-shaped trajectory, and terminated on the adjacent gyrus, it was classified as a U-fiber. Labeled fibers maintained a consistent orientation perpendicular to the local long axis of the sulcus, with terminations on the corresponding portion of the opposing gyrus at the same rostrocaudal level as the injection. Depth measurements confirmed that the bundle occupied the superficial white matter band throughout its trajectory (deep margin: mean 265 μm, SD 26.0 μm; range 230-288 μm). This bundle was therefore classified as a superficial or 'true' U-fiber, similar to the prefrontal U-fibers.

*5.2.8 Intragyral Fibers*

The remaining non-cord fibers that traveled laterally from the injection site continued within the white matter beneath the crown of the IPL and terminated in cortical areas PG and PGop within the gyrus at the level and immediately rostral to the injection site in columnar terminations (Supplementary Figure 9B, white arrowheads). These labeled fibers were predominantly observed as transverse and oblique profiles. The majority of these local intragyral fibers maintained a relatively fixed and deep position in the white matter until they became close to their termination site, at which point the fibers crossed through the most superficial portion of the white matter, and traveled for a short distance before entering the cortex.

*5.2.9 Summary*

Case BML illustrated three distinct organizational patterns. A prominent bundle adopted a U-shaped trajectory beneath the IPS but turned medially at the fundus to terminate on the medial hemisphere rather than the adjacent gyral crown, traveling below the superficial white matter band and constituting a longer association bundle rather than a U-fiber. A symmetric L-type connection was also identified along the lateral IPS bank, coursing within the superficial white matter band and terminating at the fundus without progressing to the opposing bank. Finally, the large injection, which extended into cortical area Opt, produced a U-fiber beneath the caudal STS, traveling within the superficial white matter band, terminating on the corresponding portion of the opposing preoccipital gyrus at approximately the same rostrocaudal level as the injection, and was classified as a superficial or 'true' U-fiber. Thus, this case demonstrates that U-fiber presence reflects the specific connectivity of the injected cortical area rather than the sulcus adjacent to which the case was selected.

### 6. Preoccipital Region

In the posterior parietal lobe, often referred to as the preoccipital region (Yeterian and Pandya, 2010), we examined the five remaining cases. Similar to the mid-portion of the IPS, the sulcus is deeper than in more rostral areas. However, at this level, the IPS is also operculated as it joins the lunate sulcus, and thus the morphology of the sulcus differs from that observed more rostrally. Across the five cases, no U-fibers interconnecting adjacent gyri were observed. However, shorter L- and J-type connections were present, along with deeper fiber bundles that appeared U-shaped but did not interconnect adjacent gyri.

#### *6.1 Injection Site and Long-Range Fibers*

The two representative preoccipital cases (BMEQ, BMFA) had injection sites in the caudal SPL centered on the dorsal portion of cortical area PO, extending rostrally into the caudal portion of cortical area PGm (Supplementary Figure 10B, red asterisks). In case BMEQ, the injection also extended slightly into cortical area PEa across the gyral white matter (Supplementary Figure 10E, red asterisk). Both cases exhibited a similar organization of labeled fiber bundles. The injection produced a prominent rostrally directed cord coursing ventrally through the SPL white matter core (Supplementary Figure 10E, dashed black line), following a trajectory similar to that observed in the mid-IPS SPL cases. After crossing the IPS fundus, the cord divided into two bundles: a medial bundle entering the corpus callosum and a lateral bundle entering the sagittal stratum before continuing into the internal capsule where it divided into thalamic and pontine components. Corticostriatal fibers coursed through the FOF into Muratoff's bundle, terminating in the caudate nucleus and putamen, with additional fibers coursing through the external capsule to terminate in the claustrum. Stippled transverse profiles consistent with long association fiber systems (e.g., SLF I, cingulum bundle), were also observed in the medial aspect of the SPL deep white matter.

#### *6.2 Short Association Fiber Bundles along the Intraparietal Sulcus*

In both cases, the most prominent short association fiber bundle adopted a U-shaped configuration beneath the IPS (Supplementary Figure 10C, F; dashed red lines), projecting from the medial surface of the hemisphere to the IPL crown. Labeled fibers emanated rostrally from the injection sites as predominantly transverse profiles, indicating an out-of-plane orientation parallel to the long axis of the IPS, and coursed ventrally through the SPL white matter before curving beneath the IPS. In case BMFA, a small group of sparsely labeled fibers was observed at the level of the injection, coursing beneath the fundus in deeper portions of white matter. These fibers transitioned to longitudinal profiles before fanning out within the IPL to terminate on the gyral crown in cortical areas Opt and DP. In both cases, the bulk of labeled fibers continued rostrally as transverse profiles before transitioning to longitudinal profiles as they curved around the fundus (Supplementary Figure 10C, F; dashed red lines). They then continued dorsolaterally through the center of the IPL white matter before progressively shifting to oblique and then transverse profiles beneath their cortical terminations. In case BMEQ, columnar terminations were observed in cortical area PG on the IPL crown and lateral IPS bank (Supplementary Figure 10C, red arrows). In case BMFA, terminations were present in cortical area Opt on the gyral crown and cortical area MST on the STS bank (Supplementary Figure 10F, red arrows), with additional terminations in cortical area PG further rostrally.

Although these bundles adopted a U-shaped trajectory beneath the IPS similar to the deeper U-fibers observed in the SPL cases, they originated from injections on the medial hemisphere rather than the gyral crown and did not interconnect adjacent gyri. Thus, these labeled bundles were not classified as U-fibers. Rather, these fiber bundles instead represent the reciprocal to the fiber bundle described in the IPL (case BML). Both bundles followed an oblique trajectory relative to the long axis of the sulcus and did not terminate at the same rostrocaudal level as the injection, consistent with an asymmetric organization. The relationship between these bundles and the superficial white matter band was partially obscured by shorter superficial bundles at the fundus (Supplementary Figure 10C, F; large dashed yellow line). In case BMEQ, where the superficial bundle did not progress beyond the fundus, the superficial band was identifiable along the lateral IPS bank as a region devoid of label (Supplementary Figure 10C, dashed yellow line). In case BMFA, where the superficial bundle hooked around the fundus, the two bundles were distinguished by their fiber profiles, the superficial bundle as a densely labeled band of transverse profiles and the deeper bundle as predominantly longitudinal profiles along the lateral IPS bank.
The shorter superficial fiber bundles identified in both cases were classified as L- and J-type connections. In case BMEQ, two distinct bundles were observed at different levels along the IPS. Immediately rostral to the injection, labeled fibers coursed ventrally within the superficial portions of the white matter beneath the GM-WM border along the medial IPS bank as transverse profiles, maintaining this superficial position as they hooked around the fundus and transitioned to longitudinal profiles within the IPL, with columnar terminations on the lateral IPS bank caudally in

cortical area V3A and more rostrally in area POa. Further rostrally, a second bundle was observed as transverse profiles within the superficial white matter band along the medial IPS bank (Supplementary Figure 10C, dashed teal line), providing columnar terminations in cortical area PEa along the medial bank and area IPd at the fundus (Supplementary Figure 10C, teal arrowheads). As this bundle did not progress past the fundus, it was classified as an L-type connection. In case BMFA, the superficial bundle followed a similar trajectory, coursing rostrally from the injection as transverse profiles within the superficial portions of the white matter along the medial IPS bank and providing terminations in the superior portions of cortical area PEa. Sparsely labeled fibers curved beneath the fundus, shifting to longitudinal profiles before terminating on the lateral IPS bank in cortical area V4A. Further rostrally, this bundle (Supplementary Figure 10F, dashed magenta lines) persisted within the superficial white matter band, with fibers transitioning to longitudinal profiles as they curved beneath the fundus before fanning out along the lateral IPS bank to provide terminations in the inferior portions of cortical area PEa on the medial bank and cortical area POa on the lateral bank (Supplementary Figure 10F, magenta arrowheads).

#### *6.3 Intragyral Fibers*

Similar to previous cases, intragyral fibers were observed coursing rostrally and caudally from the injection sites in both cases. The caudally directed intragyral fibers in both cases are observed as predominantly transverse fiber profiles that provide terminations in the primary visual cortex (e.g., V1 and V2). At the level of the injection site terminations are observed in cortical areas PO in both cases and area PEc in case BMFA. The majority of terminations consistent with intragyral fibers in both cases were observed rostral to the injection site, providing a series of columnar terminations along the medial hemisphere in cortical areas PGm and PO (Supplementary Figure 10F, white arrowheads).

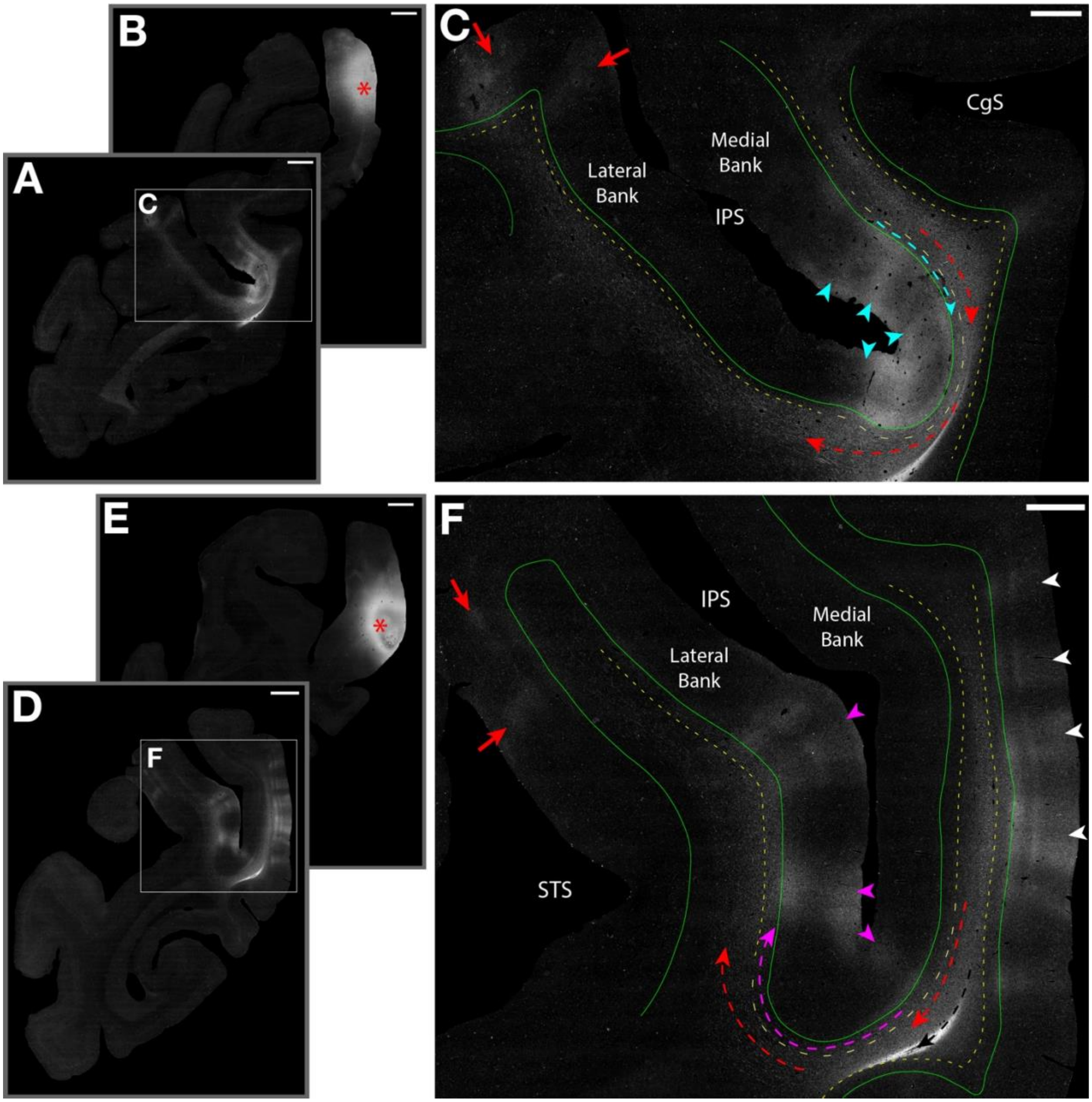


**Supplementary Figure 10:** Autoradiographs of association fibers below the mid portions of the intraparietal sulcus (IPS) that appear U-shaped but originate from the medial hemisphere. A-B. Low-magnification darkfield images are arranged rostral (A) to caudal (B) and ~10 mm apart with an injection site (A: red asterisk) on the medial hemisphere of the superior parietal lobule (SPL). C. Higher-magnification view of the boxed region in A showing multiple labeled fiber bundles. Labeled fibers with a U-shape (dashed red line) course rostrally within the SPL and then curve around the IPS forming columnar terminations on the opposing gyral crown and lateral bank of IPS (red arrows). This bundle remains in deeper portions of the white matter and avoids the superficial white matter band (dashed yellow line) immediately beneath the gray matter-white matter (GM-WM) border (solid green line). An L-type bundle (dashed teal line), courses rostrally within the superficial white matter band immediately deep to the GM-WM border and forms columnar terminations along the medial bank and fundus of the IPS (teal arrowheads). D-E. Low-magnification darkfield images are arranged rostral (D) to caudal (E) and ~10 mm apart with an injection site (E: red asterisk) on the medial hemisphere of the SPL. F. Higher-magnification view of the boxed region in D showing multiple labeled fiber bundles. The densely labeled cord (dashed black line) is located below the superficial band. Labeled fibers with a U-shape (dashed red line) follow a similar trajectory to the other case, coursing rostrally with the SPL before curving around the IPS providing terminations to the medial bank of the superior temporal sulcus (STS) (red arrows). This bundle avoids the superficial white matter band (dashed yellow line) immediately beneath the GM-WM border (solid green line). The

J-type bundle (dashed magenta line) also courses rostrally from the injection site and curves around the fundus of the IPS but instead, travels within the superficial white matter band immediately deep to the GM-WM border and forms columnar terminations near the fundus and on the lateral IPS bank (magenta arrowheads). A series of columnar terminations are observed along the medial hemisphere (white arrowheads). Panels A-C are from case BMEQ; panels D-F are from case BMFA (reflected for direct comparison between preoccipital cases). The superficial white matter band line for both cases is manually delineated based on the labeling patterns. Scale bars: A-B, D-E = 2 mm; C, F = 500 µm. CgS, Cingulate sulcus.

#### *6.4 Summary*

In both preoccipital cases, labeled bundles adopted U-shaped trajectories beneath the IPS but did not interconnect adjacent gyri, and were therefore not classified as U-fibers. These bundles traveled in deeper portions of the white matter, avoiding the superficial white matter band throughout their trajectory around the sulcus. In contrast, shorter J- and L-type connections were identified in both cases coursing within the superficial white matter band along the medial IPS bank and around the sulcus, with terminations limited to the sulcal banks and fundus. Together, these cases reinforce the organizational principle that shorter fiber bundles terminating locally along a sulcus travel within the superficial white matter band, while bundles projecting to more distant targets course in deeper portions of the white matter and avoid this band until approaching their cortical terminations.

### 7. Abbreviations

FOF, fronto-occipital fasciculus

GM-WM, gray matter-white matter

ILF, inferior longitudinal fasciculus

IPL, inferior parietal lobule

IPS, intraparietal sulcus

MdLF, middle longitudinal fasciculus

Pro, proisocortex

ProM, proisocortical motor cortex

SII, secondary somatosensory cortex

SLF, superior longitudinal fasciculus

SPL, superior parietal lobule

STS, superior temporal sulcus

### 8. References

Catani, M., Dell'acqua, F., Vergani, F., Malik, F., Hodge, H., Roy, P., Valabregue, R., & Thiebaut de Schotten, M. (2012). Short frontal lobe connections of the human brain. Cortex; a

Journal Devoted to the Study of the Nervous System and Behavior, 48(2), 273–291. https://doi.org/10.1016/j.cortex.2011.12.001

DeFelipe, J., Conley, M., & Jones, E. G. (1986). Long-range focal collateralization of axons arising from corticocortical cells in monkey sensory-motor cortex. The Journal of Neuroscience: The Official Journal of the Society for Neuroscience, 6(12), 3749–3766. https://doi.org/10.1523/jneurosci.06-12-03749.1986

Jones, E. G., Coulter, J. D., & Hendry, S. H. (1978). Intracortical connectivity of architectonic fields in the somatic sensory, motor and parietal cortex of monkeys. The Journal of Comparative Neurology, 181(2), 291–347. https://doi.org/10.1002/cne.901810206

Künzle, H. (1978). Cortico-cortical efferents of primary motor and somatosensory regions of the cerebral cortex in Macaca fascicularis. Neuroscience, 3(1), 25–39. https://doi.org/10.1016/0306-4522(78)90151-3

Mufson, E. J., & Pandya, D. N. (1984). Some observations on the course and composition of the cingulum bundle in the rhesus monkey. The Journal of Comparative Neurology, 225(1), 31–43. https://doi.org/10.1002/cne.902250105

Petrides, M., & Pandya, D. N. (1984). Projections to the frontal cortex from the posterior parietal region in the rhesus monkey. The Journal of Comparative Neurology, 228(1), 105–116. https://doi.org/10.1002/cne.902280110

Petrides, M., & Pandya, D. N. (2006). Efferent association pathways originating in the caudal prefrontal cortex in the macaque monkey. The Journal of Comparative Neurology, 498(2), 227–251. https://doi.org/10.1002/cne.21048

Petrides, Michael, & Pandya, D. N. (1988). Association fiber pathways to the frontal cortex from the superior temporal region in the rhesus monkey. The Journal of Comparative Neurology, 273(1), 52–66. https://doi.org/10.1002/cne.902730106

Petrides, Michael, & Pandya, D. N. (2009). Distinct parietal and temporal pathways to the homologues of Broca's area in the monkey. PLoS Biology, 7(8), e1000170. https://doi.org/10.1371/journal.pbio.1000170

Schmahmann, J. D., & Pandya, D. N. (1989). Anatomical investigation of projections to the basis pontis from posterior parietal association cortices in rhesus monkey. The Journal of Comparative Neurology, 289(1), 53–73. https://doi.org/10.1002/cne.902890105

Schmahmann, J. D., & Pandya, D. N. (1991). Projections to the basis pontis from the superior temporal sulcus and superior temporal region in the rhesus monkey. The Journal of Comparative Neurology, 308(2), 224–248. https://doi.org/10.1002/cne.903080209

Schmahmann, J. D., & Pandya, D. N. (1992). Course of the fiber pathways to pons from parasensory association areas in the rhesus monkey. The Journal of Comparative Neurology, 326(2), 159–179. https://doi.org/10.1002/cne.903260202

Schmahmann, J. D., & Pandya, D. N. (1993). Prelunate, occipitotemporal, and parahippocampal projections to the basis pontis in rhesus monkey. The Journal of Comparative Neurology, 337(1), 94–112. https://doi.org/10.1002/cne.903370107

Schmahmann, J. D., & Pandya, D. N. (1997). Anatomic organization of the basilar pontine projections from prefrontal cortices in rhesus monkey. The Journal of Neuroscience: The Official Journal of the Society for Neuroscience, 17(1), 438–458. https://doi.org/10.1523/jneurosci.17-01-00438.1997

Schmahmann, Jeremy D., & Pandya, D. (2006). Fiber pathways of the brain. Oxford University Press. https://doi.org/10.1093/acprof:oso/9780195104233.001.0001

Schmahmann, Jeremy D., Rosene, D. L., & Pandya, D. N. (2004). Motor projections to the basis pontis in rhesus monkey. The Journal of Comparative Neurology, 478(3), 248–268. https://doi.org/10.1002/cne.20286

Seltzer, B., & Pandya, D. N. (1989). Frontal lobe connections of the superior temporal sulcus in the rhesus monkey. The Journal of Comparative Neurology, 281(1), 97–113. https://doi.org/10.1002/cne.902810108

Siwek, D. F., & Pandya, D. N. (1991). Prefrontal projections to the mediodorsal nucleus of the thalamus in the rhesus monkey. The Journal of Comparative Neurology, 312(4), 509–524. https://doi.org/10.1002/cne.903120403

Yamashita, A., & Arikuni, T. (2001). Axon trajectories in local circuits of the primary motor cortex in the macaque monkey (Macaca fuscata). Neuroscience Research, 39(2), 233–245. https://doi.org/10.1016/s0168-0102(00)00220-0

Yeterian, E. H., & Pandya, D. N. (1985). Corticothalamic connections of the posterior parietal cortex in the rhesus monkey. The Journal of Comparative Neurology, 237(3), 408–426. https://doi.org/10.1002/cne.902370309

Yeterian, E. H., & Pandya, D. N. (1991a). Corticothalamic connections of the superior temporal sulcus in rhesus monkeys. Experimental Brain Research, 83(2), 268–284. https://doi.org/10.1007/bf00231152

Yeterian, E. H., & Pandya, D. N. (1991b). Prefrontostriatal connections in relation to cortical architectonic organization in rhesus monkeys. The Journal of Comparative Neurology, 312(1), 43–67. https://doi.org/10.1002/cne.903120105

Yeterian, E. H., & Pandya, D. N. (1993). Striatal connections of the parietal association cortices in rhesus monkeys. The Journal of Comparative Neurology, 332(2), 175–197. https://doi.org/10.1002/cne.903320204

Yeterian, E. H., & Pandya, D. N. (1995). Corticostriatal connections of extrastriate visual areas in rhesus monkeys. The Journal of Comparative Neurology, 352(3), 436–457. https://doi.org/10.1002/cne.903520309

Yeterian, E. H., & Pandya, D. N. (1997). Corticothalamic connections of extrastriate visual areas in rhesus monkeys. The Journal of Comparative Neurology, 378(4), 562–585. https://doi.org/10.1002/(sici)1096-9861(19970224)378:4%253C562::aid-cne10%253E3.0.co;2-l

Yeterian, E. H., & Pandya, D. N. (1998). Corticostriatal connections of the superior temporal region in rhesus monkeys. The Journal of Comparative Neurology, 399(3), 384–402. https://doi.org/10.1002/(sici)1096-9861(19980928)399:3%253C384::aid-cne7%253E3.0.co;2-x

Yeterian, E. H., & Pandya, D. N. (2010). Fiber pathways and cortical connections of preoccipital areas in rhesus monkeys. The Journal of Comparative Neurology, 518(18), 3725–3751. https://doi.org/10.1002/cne.22420
